# The voltage and spiking responses of subthreshold resonant neurons to structured and fluctuating inputs: persistence and loss of resonance and variability

**DOI:** 10.1101/2021.06.14.448368

**Authors:** Rodrigo F. O. Pena, Horacio G. Rotstein

## Abstract

We systematically investigate the response of neurons to oscillatory currents and synaptic-like inputs and we extend our investigation to non-structured synaptic-like spiking inputs with more realistic distributions of presynaptic spike times. We use two types of chirp-like inputs consisting of (i) a sequence of cycles with discretely increasing frequencies over time, and (ii) a sequence having the same cycles arranged in an arbitrary order. We develop and use a number of frequency-dependent voltage response metrics to capture the different aspects of the voltage response, including the standard impedance (*Z*) and the peak-to-trough amplitude envelope (*V_ENV_*) profiles. We show that *Z*-resonant cells (cells that exhibit subthreshold resonance in response to sinusoidal inputs) also show *V_ENV_*-resonance in response to sinusoidal inputs, but generally do not (or do it very mildly) in response to square-wave and synaptic-like inputs. In the latter cases the resonant response using *Z* is not predictive of the preferred frequencies at which the neurons spike when the input amplitude is increased above subthreshold levels. We also show that responses to conductance-based synaptic-like inputs are attenuated as compared to the response to current-based synaptic-like inputs, thus providing an explanation to previous experimental results. These response patterns were strongly dependent on the intrinsic properties of the participating neurons, in particular whether the unperturbed *Z*-resonant cells had a stable node or a focus. In addition, we show that variability emerges in response to chirp-like inputs with arbitrarily ordered patterns where all signals (trials) in a given protocol have the same frequency content and the only source of uncertainty is the subset of all possible permutations of cycles chosen for a given protocol. This variability is the result of the multiple different ways in which the autonomous transient dynamics is activated across cycles in each signal (different cycle orderings) and across trials. We extend our results to include high-rate Poisson distributed current- and conductance-based synaptic inputs and compare them with similar results using additive Gaussian white noise. We show that the responses to both Poisson-distributed synaptic inputs are attenuated with respect to the responses to Gaussian white noise. For cells that exhibit oscillatory responses to Gaussian white noise (band-pass filters), the response to conductance-based synaptic inputs are low-pass filters, while the response to current-based synaptic inputs may remain band-pass filters, consistent with experimental findings. Our results shed light on the mechanisms of communication of oscillatory activity among neurons in a network via subthreshold oscillations and resonance and the generation of network resonance.

## 1 Introduction

Subthreshold (membrane potential) oscillations (STOs) in a variety of frequency ranges have been observed in many neuron types [1–40]. In brain areas such as the entorhinal cortex, the hippocampus and the olfactory bulb, the frequency of the STOs is correlated with the frequency of the networks of which they are part [1, 2, 26–28, 34, 41–44], thus suggesting STOs play a role in the generation of network rhythms [11, 26, 45], the communication of information across neurons in a network via timing mechanisms [6, 29, 46, 47], cross-frequency coupling in neurons where STOs are interspersed with spikes (mixed-mode oscillations; MMOs) [48–55], and the encoding of information [56, 57] and sensory processing [39]. STOs can be generated by cellular intrinsic or network mechanisms. In the first case, STOs result from the interplay of ionic currents that provide positive and slower negative effects (e.g., [3, 5, 58, 59]). (Examples of the former are the persistent sodium and calcium activation and examples of the latter are h-type hyperpolarization-activated mixed sodium-potassium, M-type slow potassium and calcium inactivation.) In the second case, STOs are generated in networks, but the individual cells cannot robustly oscillate when isolated (e.g., [20, 60, 61]).

The communication of oscillatory information among neurons in a network and across brain areas requires the generation of spiking patterns that are correlated with the underlying STOs (e.g., MMOs where spikes occur at the peak of the STO or at a consistent phase referred to this peak). It also requires the system to be able to respond to external inputs in such a way as to preserve the oscillatory information. Studies on the latter are typically carried out by using sinusoidal inputs. Subthreshold (membrane potential) resonance (MPR) refers to the ability of a system to exhibit a peak in their voltage amplitude response to oscillatory inputs currents at a preferred (resonant) frequency [62–65] (in voltage-clamp, the input is voltage and the output is current). MPR has been investigated in many neuron types both experimentally and theoretically [7–9, 17, 62–107] and it has been shown to have functional implications for the generation of network oscillations [92, 108].

The choice of sinusoidal inputs is based on the fact that for linear systems they can be used to reconstruct the response to arbitrary time-dependent inputs, and relatively good approximations can be obtained for mildly nonlinear systems. However, although neurons may be subject to oscillatory modulated inputs, the communication between neurons in a network occurs via synaptic connections whose waveforms are significantly different from pure sinusoidal functions. Synaptic inputs such as these associated with AMPA or GABA_*A*_ synaptic currents rise very fast (almost instantaneously) and then decay exponentially on a slower time scale. In contrast to sinusoidal inputs, the rise and decay of the periodic synaptic inputs occur over a small portion of the input periods for the smaller input frequencies. The gradual variation of the sinusoidal inputs causes the voltage response to reach the stationary regime after a very small number of cycles, while the abrupt changes in the synaptic inputs over a small time interval sequentially activate the autonomous transient dynamics at every cycle, and therefore is expected to produce different response patterns than these for sinusoidal inputs [109].

The main goal of this paper is to understand whether and under what conditions the presence of MPR in a neuron is predictive of the preferred frequency at which the neuron will spike in response to periodic presynaptic inputs when the input amplitude is increased above subthreshold levels.

From a dynamical systems perspective, single-cell sustained STOs can be in the limit cycle regime (robust to noise, driven by DC inputs) or in the fluctuation-driven regime (vanishing or decaying to an equilibrium in the absence of noise). Noisy STOs in the limit cycle regime reflect the stationary dynamics of the system in the absence of noise. In contrast, fluctuation-driven STOs reflect the autonomous transient dynamics (the transient dynamics of the underlying unperturbed system) [109]. The effects of the autonomous transient dynamics are captured by the system’s response to abrupt changes in constant inputs [109]. There, the values of the voltage and other variables at the end of a constant input regime become the initial conditions for the new one. By repeated activation of the autonomous transient dynamics, piecewise constant inputs with short-duration pieces and arbitrarily distributed amplitudes (not necessarily randomly distributed) are able to produce oscillatory responses [109]. Noise-driven oscillations are a limiting case of this mechanism where the input’s constant pieces have randomly distributed amplitudes and their durations approach zero. If the amplitudes are normally distributed, these piecewise constant inputs provide an approximation to Gaussian white noise [110]. Roughly speaking, each “kick” to the system by the noisy input operates effectively as an abrupt change of initial conditions to which the system responds by activating the transient time scales, and the voltage and other state variables evolve according to the vector field away from equilibrium (or stationary state). For example, noise-driven STOs [58, 109, 111, 112] can be generated when damped oscillations are amplified by noise, and this may extend to situations where the noiseless system exhibits overdamped oscillations (overshoots) [109].

Along these lines of the previous discussion, a series of dynamic clamp experiments [19] using artificially generated synaptic conductances and currents driven by high-rate presynaptic Poisson spike trains showed that medial entorhinal cortex layer II stellate cells (SCs) are able to generate STOs in response to current-based synapses, but not in response to conductance-based synaptic currents. SCs are a prototypical example of an intrinsic fluctuation-driven STO neuron [3, 5, 58, 113] and resonator [68]. In the response to current-based synaptic inputs, the STOs have similar frequencies and amplitudes as the spontaneous STOs [3, 5] and the resonant responses to sinusoidal inputs [68]. In response to conductance-based synaptic currents, STOs may still be present, but highly attenuated as compared to current-based synaptic inputs. Similar results were found in [114] for hippocampal CA1 OLM (oriens lacunosum-moleculare) cells and in [115, 116] for phase-response curves in subthalamic neurons.

This raises a seeming contradiction between the ability of the impedance (*Z*-) profile (curve of the voltage *V*-response normalized by the amplitude of the oscillatory inputs as a function of the input frequency) to predict the existence of STOs for arbitrary time-dependent inputs, in particular Gaussian white noise, and the absence of STOs for conductance-based synaptic inputs in response to Poisson-distributed spike trains whose effect on the target cells have been approximated by Gaussian white noise [117–122]. This can be partially explained by the fact that synaptic currents “add linearity” to the system, but fluctuation-driven STOs can be generated in linear systems and therefore one could expect only changes in amplitude and frequency. Another possible explanation is that while the *Z*-profile is independent of the input waveform, the voltage response power spectral density (PSD) is not and depends on the current input waveform. The expectation that the PSD be similar for current-based Gaussian white noise and Poisson-driven current-/conductance-based synaptic inputs would assume similarity between the different input types.

In this paper, we systematically address these issues in a broader context. Our results shed light on the mechanisms of communication of oscillatory activity among neurons in a network via subthreshold oscillations and resonance and the generation of suprathreshold and network resonance.

## 2 Methods

### 2.1 Models

In this paper, we use relatively simple biophysically plausible models describing the subthreshold dynamics of individual neurons subject to both additive and multiplicative inputs.

#### 2.1.1 Linear model: additive input current

For the individual neurons we use the following linearized biophysical (conductance-based) model

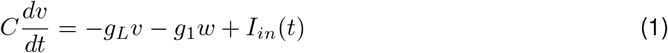

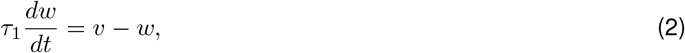

where *v* (mV) is the membrane potential relative to the voltage coordinate of the fixed-point (equilibrium potential) of the original model, *w* (mV) is the recovery (gating) variable relative to the gating variable coordinate of the fixed-point of the original model normalized by the derivative of the corresponding activation curve, *C* (*μ*F/cm^2^) is the specific membrane capacitance, *g_L_* (mS/cm^2^) is the linearized leak conductance, *g*_1_ (mS/cm^2^) is the linearized ionic conductance, *τ*_1_ (ms) is the linearized gating variable time constant and *I_in_*(*t*) (*μ*A/cm^2^) is the time-dependent input current. In this paper we consider resonant gating variables (*g*_1_ > 0; providing a negative feedback effect). We refer the reader to [63, 64] for details of the description of the linearization process for conductance-based models.

#### 2.1.2 Conductance-based synaptic input model: multiplicative input

To account for the effects of conductance-based synaptic inputs we extend the model (1)-(2) to include a synaptic current

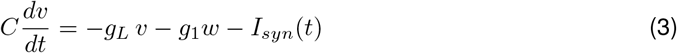

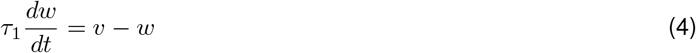

where

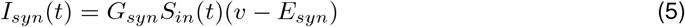

and *G_syn_* (mS/cm^2^) is the maximal synaptic conductance, *E_syn_* (mV) is the synaptic reversal potential (*E_ex_* for excitatory inputs and *E_in_* for inhibitory inputs) and *S_in_*(*t*) is the time-dependent synaptic input.

#### 2.1.3 *I*_Nap_ + *I*_h_ conductance-based model

To test our ideas in a more realistic model we will use the following conductance-based model combining a fast amplifying gating variable associated to the persistent sodium current *I_Nap_* and a slower resonant gating variable associated to the hyperpolarization-activated mixed cation (h-) current *I_h_*. The model equations for the subthreshold dynamics read

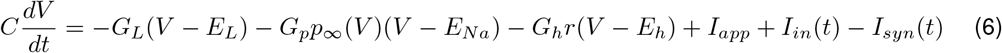

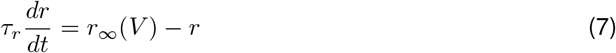

where

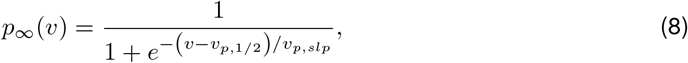

and

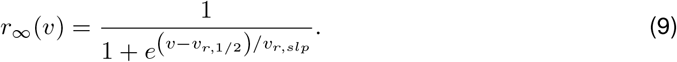

This model describes the onset of spikes, but not the spiking dynamics [58]. Spikes are added by including a voltage threshold (indicating the occurrence of a spike after its onset) and reset values *V_rst_* and *r_rst_* for the participating variables.

Unless stated otherwise, we use the following parameter values: *v*_*p*,1/2_ = −38 mV, *v_p,slp_* = 6.5 mV, *v*_*r*,1/2_ = −79.2 mV, *v_r,slp_* = 9.78 mV, *C* =1 *μ*F/cm^2^, *E*_L_ = −65 mV, *E*_Na_ = 55 mV, *E*_h_ = −20 mV, *g*_L_ = 0.5 mS/cm^2^, *g_p_* = 0.5 mS/cm^2^, *g*_h_ = 1.5 mS/cm^2^, *I*_app_ = −2.5 *μ*A/cm^2^, and *τ_r_* = 80 ms.

### 2.2 Input functions: periodic inputs and realistic waveforms

The input functions *I_in_*(*t*) and *S_in_* we use in this paper have the general form

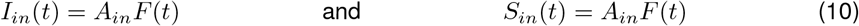

#### 2.2.1 Chirp-like input functions: increasingly ordered frequencies

We will use the three chirp-like input functions *F*(*t*) shown in Fig. 1-a. The sinusoidal chirp-like function (Fig. 1-a1) consists of a sequence of input cycles with discretely increasing frequencies over time (Fig. 1-a4). We use integer frequencies in the range 1 – 100 Hz. These chirp-like functions are a modification of the standard chirp function [69] where the frequency of the sinusoidal input increases (or decreases) continuously with time [69]. Sinusoidal inputs of a single frequency and sinusoidal chirps with monotonically and continuously increasing (or decreasing) frequencies with time have been widely used to investigate the resonant properties of neurons both *in vitro* and *in vivo* [69, 78, 123].

**Figure 1:**
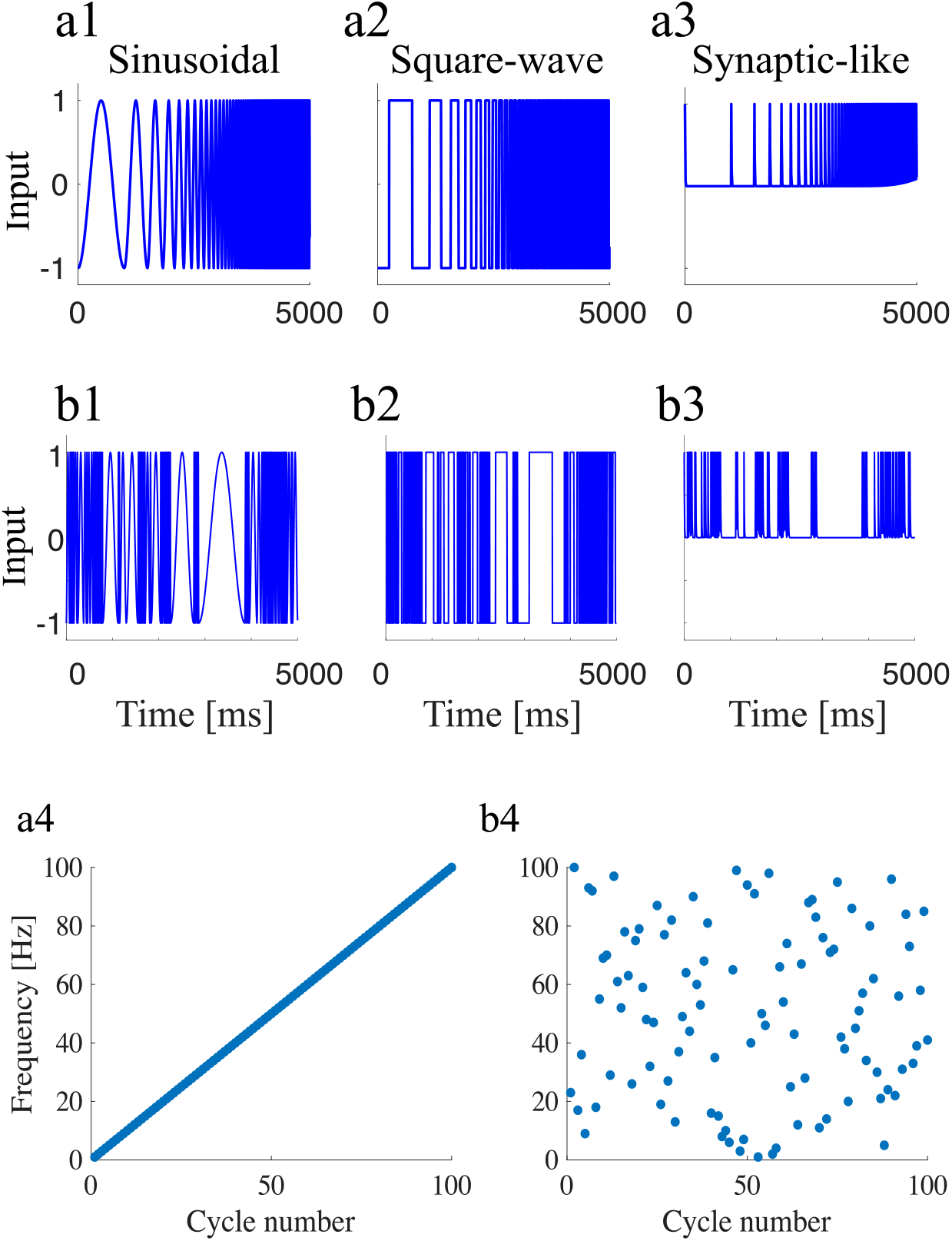
Chirp-like input functions. (a1) Sinusoidal chirp-like input. (a2) Square-wave chirp-like input. (a3) Synaptic-like chirp input. (b) Same input functions as in (a) but with arbitrarily ordered frequencies. (a4) Increasingly ordered frequencies used to construct (a). (b4) Arbitrarily ordered frequencies used to construct (b). All panels show frequencies in the range 1–100 Hz.

The square-wave (Fig. 1-a2) and synaptic-like (Fig. 1-a3) chirp-like functions are constructed in the same manner as the sinusoidal one by substituting the sinusoidal functions by square waves (duty cycle = 0.5) and exponentially decreasing functions with a characteristic decay time *τ*_Dec_, respectively. We refer to them as sinusoidal, square-wave, and synaptic-like inputs or chirps, respectively (and we drop the “chirp-like”).

The discretely changing chirp-like functions we use here are a compromise between tractability and the ability to incorporate multiple frequencies in the same input signal. They combine the notion of input frequency with the notion of transition between different frequencies in the same signal. Also, note that the square-wave inputs are an intermediate between sinusoidal and synaptic-like inputs in the sense that square-wave inputs have abrupt transitions as synaptic-like inputs but are closer in shape to the sinusoidal input that is changed gradually.

#### 2.2.2 Chirp-like input functions: arbitrarily ordered frequencies

To examine the variability of the cell’s response to the chirp signals described above and to capture the fact that information does not necessarily arrive in a regularly ordered manner, we will use modified versions of these chirp inputs where the cycles are rearranged in an arbitrary order (Fig. 1-b). The regularly (Fig. 1-a4) and arbitrarily (Fig. 1-b4) ordered input signals have exactly the same cycles (one cycle for each frequency value within the considered range) and therefore the same frequency content.

#### 2.2.3 Poisson distributed spikes and white Gaussian noise

To test the oscillatory responses to more realistic inputs we use spike-trains with distributed spikes following a homogeneous Poisson process with rate *ν*. Each input spike evokes a synaptic-like input function as described above. In addition, we use an additive Gaussian noise current 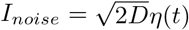 where *η*(*t*) is a Gaussian white noise input with zero mean and unit variance (*I_in_*(*t*) has zero mean and variance 2*D*). Unless stated otherwise, *ν* = 1000 Hz and *D* = 1 (additional information is provided in the figure captions).

### 2.3 Output Metrics

#### 2.3.1 Impedance (amplitude) profile

The impedance (amplitude) profile is defined as the magnitude of the ratio of the output (voltage) and input (current) Fourier transforms

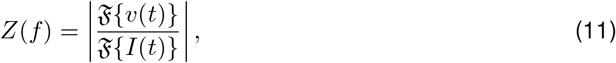

where 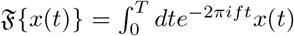. In practice, we use the Fast Fourier Transform algorithm (FFT) to compute 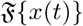. Note that *Z*(*f*) is typically used as the complex impedance, a quantity that has amplitude and phase. For simplicity, here we used the notation *Z*(*f*) for the impedance amplitude.

#### 2.3.2 Voltage and impedance (amplitude) envelope profiles

The upper and lower envelope profiles 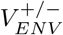 are curves joining the peaks and troughs of the steady state voltage response as a function of the input frequency *f*. The envelope impedance profile is defined as [65, 105]

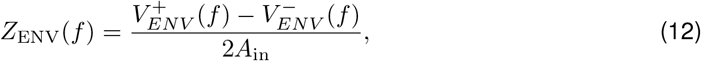

where *A*_in_ is the input amplitude. For linear systems, *Z*_ENV_(*f*) coincides with *Z*(*f*).

#### 2.3.3 Voltage power spectral density

In the frequency-domain, we compute the power spectral density (PSD) of the voltage as the absolute value of its Fourier transform 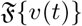. We will refer to this measure as PSD or *V*_PSD_.

#### 2.3.4 Firing rate (suprathreshold) response

We compute the firing rate response of a neuron by counting the number of spikes fired within an interval of length *T* and normalizing by *T*

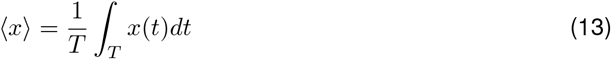

where the neural function *x* is given by

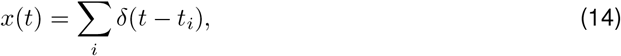

and *t_i_* are the spike times within the considered interval.

### 2.4 Numerical simulations

We used the modified Euler method (Runge-Kutta, order 2) [124] with step size Δ*t* = 0.01 ms. All neural models and metrics, including phase-plane analysis, were implemented by self-developed Matlab routines (The Mathworks, Natick, MA) and are available in https://github.com/BioDatanamics-Lab/impedance_input_dependent.

## 3 Results

### 3.1 Transient and steady-state neuronal responses to abrupt versus gradual input changes

The properties of the transient responses of dynamical systems to external inputs depend on the intrinsic properties of the target cells, the initial conditions of the participating variables and the nature of the attractors (assumed to exist). The complexity of the autonomous transient dynamics, defined as the transient response to abrupt changes in constant inputs, increases with the model complexity. For example, for the simplest, passive neuron (a one-dimensional system), the voltage *V* evolves monotonically towards the new equilibrium value determined by a constant input. Two-dimensional neurons having a restorative current with slow dynamics (e.g., *I_h_, I_M_*, *I_CaT_* inactivation) may display overshoots and damped oscillations (Fig. 2), which can be amplified by fast regenerative currents (e.g., *I_Nap_, I_Kir_*, *I_CaT_* activation), and are more pronounced the further away the initial conditions are from the equilibrium (not shown) and the more abrupt is the input change. Here we review the dependence of the transient response properties of relatively simple models with the properties of the input and discuss some of the implications for the steady-state responses of the same systems to periodic inputs.

**Figure 2:**
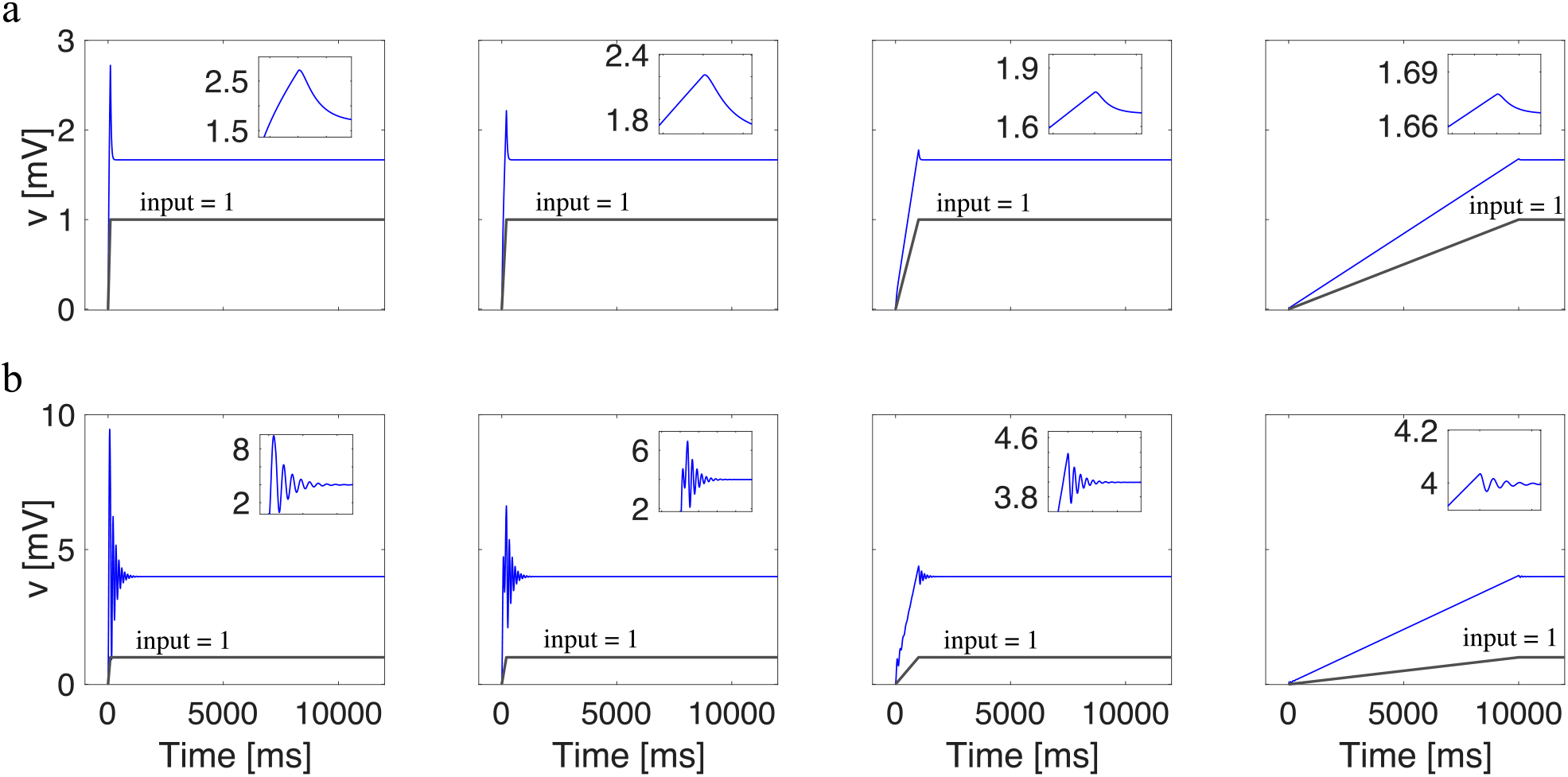
Transient response to input decreases as the input changes from abrupt to gradual. Input current increases from 0 to 1 (black solid line) but the strength of the transient decreases in every column. First row: Example with overshoot, parameters are *g*_L_ = 0.2 and *g*_1_ = 0.4. Second row: Example with subthreshold oscillations, parameters are *g*_L_ = 0.00025 and *g*_1_ = 0.25. The insets show a zoom-in on the transients.

#### 3.1.1 The strength of the transient responses to input changes decreases as the input changes transition from abrupt to gradual

An abrupt change in the input current (e.g., step DC input) can be interpreted as causing a sudden translation of the equilibria (for the voltage and other state variables) in the phase-space diagram from its baseline location (e.g., Fig. 3-a, *I* = 0, intersection between the *V*- and *w*-nullclines, solid-red and green, respectively) to the to a new location determined by the DC value (e.g., Fig. 3-a, *I* = 1, intersection between the displaced *V*- and *w*-nullclines, dashed-red and green, respectively). The values of these state variables prior to the transition become the initial conditions with respect to this new equilibrium. Therefore, the voltage responses to abrupt changes in the input currents are expected to exhibit overshoots and damped oscillations (Fig. 3, insets, and Fig. 2, left), which are more pronounced the stronger the input (not shown). As the change in input current becomes more gradual, the transient effects become more attenuated (Fig. 2, middle) and eventually the voltage response becomes almost monotonic (Fig. 2, right).

**Figure 3:**
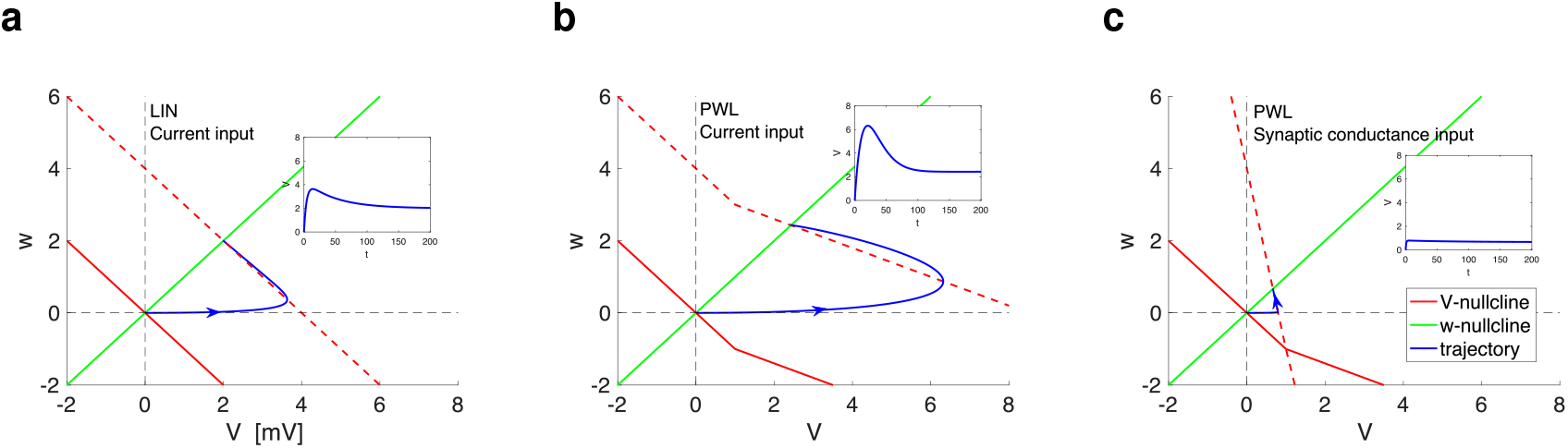
Nonlinear transient response amplifications and attenuations in current- vs. conductance-based inputs. Phase-plane diagrams for *I* = 1. The solid-red curve represents the *V*-nullcline (*dv/dt* = 0) for *I* = 0, the dashed-red curve represents the *V*-nullcline (*dv/dt* = 0) for *I* =1, the solid-green curve represents the *w*-nullcline (*dw/dt* = 0) for *I* = 0, the solid-blue curve represents the trajectory, and the dashed-gray lines are marking the point where the trajectory initially starts at (0, 0) (the fixed-point for *I* = 0), converging to the fixed-point for *I* =1. The insets show the *V* traces. The 2D linear system exhibits an overshoot in response to step-constant inputs and resonance in response to oscillatory inputs [63, 64, 105]. **a.** Linear (LIN) model described by eqs. (1)–(2). **b.** Current-based piecewise linear (PWL) model described by 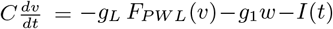 and 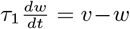 where *F_PWL_*(*v*) = *v* for *v* < *v_c_* and *F_PWL_*(*v*) = *v_c_*+*g_c_*/*g_L_*(*v–v_c_*) for *v* > *v_c_*. **c.** Conductance-based piecewise linear (PWL) model described by 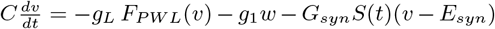 and 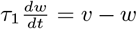 where *F_PWL_*(*v*) = *v* for *v* < *v_c_* and *F_PWL_*(*v*) = *v_c_* + *g_c_*/*g_L_*(*v* – *v_c_*) for *v* > *v_c_* (with *S* substituted by *I*). We used the following parameter values: *C* = 1, *g_L_* = 0.25, *g*_1_ = 0.25, *τ*_1_ = 100 (same as in Fig. 4), *v_c_* = 1 and *g_c_* = 0.1.

This transition in the strength of the transient responses is expected since a monotonic input change can be approximated by a sequence of smaller step input changes of increasing (constant) magnitude, each one producing a transient response, which becomes smaller the larger the number of steps (the smaller the step size) since the initial conditions for each step in the partition are very close to the corresponding (new) steady state. Therefore, for input transitions between the same constant values, but with different slopes, the amplitude of the transient response becomes more attenuated the more gradual the transition since a larger partition is required in order to keep the step size constant.

#### 3.1.2 Nonlinear amplification of the transient and steady-state response to constant inputs

Certain types of nonlinearities have been shown to amplify the response of neuronal systems (and dynamical systems in general) to the same input. This is reflected in both the responses to constant and oscillatory inputs. For illustrative purposes, in Figs. 3-b and -c we use a piecewise linear (PWL) model obtained from the linear model (1)-(2) by making the *v*-nullcline a continuous PWL function. It was shown in [105] that this type of model displays nonlinear amplification of the voltage response to sinusoidal inputs and captures similar phenomena observed in nonlinear models, in particular these having parabolic-like *V*-nullclines describing the subthreshold voltage dynamics [65].

Figs. 3-a and -b show the superimposed phase-plane diagrams for the PWL model (b) and the linear (LIN) model (a) from which it originates for a constant input amplitude *I* = 0 (solid-red, baseline) and *I* =1 (dashed-red). The *w*-nullcline is unaffected by changes in *I*. The trajectory (blue), initially at the fixed-point for *I* = 0, converges towards the fixed-point for *I* =1. For low enough values of *I* (lower than in Figs. 3-a and -b) the trajectory remains within the linear region (the trajectory does not reach the *V*-nullcline’s “breaking point” value of *V*) and therefore the dynamics are not affected by the nonlinearity. In both cases (panels a and b), the response exhibits an overshoot. The peak occurs when the trajectory is able to cross the *V*-nullcline. Because the *V*-nullcline’s “right piece” has a smaller slope than the “left piece”, the trajectory is able to reach larger values of *V* before turning around. This amplification is particularly stronger for the transient dynamics (initial upstroke) than for the steady-state response. Nonlinear response amplifications in this type of system are dependent on the time scale separation between the participating variables. For smaller values of *τ*_1_ this nonlinear amplification is reduced and although the system is nonlinear, it behaves quasi-linearly [65, 105].

#### 3.1.3 Attenuation of the transient and steady-state response of conductance-based versus current-based (constant) synaptic inputs

From the phase-plane diagram in Fig. 3-c we see that increasing values of *I* (here we have a conductance and a driving-force in the model) reduces the nonlinearity of the *V*-nullcline (dashed-red) and increases (in absolute value) its slope. Both phenomena oppose the response amplification (blue) and the overshoot becomes much less prominent. The triangular region (bounded by the *V*-axis, the displaced *V*-nullcline (dashed-red) and the *w*-nullcline (green)) is reduced in size as compared to the current-based inputs (panel b) and therefore the response is reduced in amplitude. Moreover, because the displaced *V*-nullcline in panel c is more vertical than the baseline *V*-nullcline, the size of the overshoot in response to constant inputs is reduced and, in this sense, the responses become quasi-1D. As a consequence, the initial portion of the transient responses to abrupt changes in input is reduced in size and the oscillatory response to PWC inputs is also attenuated and the resonant peak disappears [109].

#### 3.1.4 Implications for the neuronal responses to structured (periodic) and non-structured inputs: hypotheses and questions

An immediate consequence of these observations is the prediction that a system’s responses to square-wave and sinusoidal inputs of the same frequency (and duty cycle) will be qualitatively different, and these differences will depend on the stability properties of the unperturbed cells (e.g., stable nodes versus foci, overshoots versus damped oscillations). The steady-state response to periodic inputs can be interpreted as a sequence of transient responses to input changes. These transient effects (autonomous transient dynamics) are expected to be prominent in the steady-state responses to square-wave inputs, but not in the steady-state responses to sinusoidal inputs. Sinusoidal and square-wave inputs are representative of gradually and abruptly changing signals, respectively, and are amenable for comparison. The result of these comparisons sheds light on more realistic signals such as synaptic-like ones. To test these ideas we will use the chirp-like input currents shown in Fig. 1 (see Section 2.2.1).

A second immediate consequence of the observations referred to above is the finding that a system’s amplitude response to piecewise constant (PWC) inputs having the same set of constant pieces arranged in different order is variable with respect to each other [109]. At the population level (same cell receiving a number of input signals consisting of permutations of the order of the same constant pieces of the same baseline signal), the properties of this variability crucially depend on the cell’s autonomous transient dynamics. These reflect the multiple ways in which the cell responds to a given constant piece input from the values determined by the responses to the previous piece in the inputs signal, which change across trials. Interestingly, this phenomenon does not require the constant piece amplitudes to be randomly distributed, but they can be generated by a deterministic rule consisting of a baseline input pattern (e.g., increasing order of amplitudes) and a subset of all possible permutations of the constant piece amplitude order. We analyzed this in detail in the companion paper [109].

The issues discussed above raise a number of questions. First, whether and under what conditions the frequency-preference properties of a system’s response to sinusoidal inputs are predictive of the response properties of the same system to other types of inputs. While the Fourier theorem guarantees that the latter can be reconstructed from the former if properly normalized, it does not guarantee that the two will have the same waveform and the same frequency-dependence properties using metrics that depend on these waveforms since the normalization factors (related to the input) may have different frequency dependencies. Second, whether and under what conditions the differences between the preferred frequency-band response of sinusoidal and non-sinusoidal inputs, if they exist, persist in the spiking regime. Given that the communication between neurons occurs via synaptic interactions, the failure of the responses to synaptic-like inputs to replicate the frequency-preference properties in response to sinusoidal inputs would indicate that the latter, although useful for the reconstruction of signals, does not have direct implications for the spiking dynamics. Third, whether and under what conditions the frequency-preference properties of a system’s response to structured (deterministic) inputs are predictive of the responses of the same system to unstructured (noisy) inputs. Fourth, whether and under what conditions the oscillatory (intrinsic) and resonant properties of cells result from the very brief initial transients of their autonomous dynamics. Fifth, how does the variability of a cell’s response to different input trials is processed by the feedback effects operating at the cell level. We address these issues in the next sections.

### 3.2 Subthreshold resonance in response to sinusoidal inputs is captured by the impedance and voltage envelope profiles

#### 3.2.1 Subthreshold resonance

A cell is said to exhibit subthreshold resonance if its voltage amplitude response to subthreshold oscillatory inputs peaks at a preferred (resonant) frequency (Figs. 4-a and 5-a). These responses are typically measured by computing the impedance *Z*, defined as the quotient of the power spectra of the output and input (see Methods). In current clamp, the input is current and the output is voltage. In controlled experiments and simulations, the unperturbed cells are in equilibrium in the absence of the oscillatory inputs. In response to constant inputs, resonant cells may be non-oscillators, typically exhibiting an overshoot, or exhibit oscillatory behavior (e.g., damped oscillations) (e.g., see Figs. 4-b1 and 5-b1, respectively, where the behavior can be observed in the first cycle). Therefore, resonance is not uncovering an oscillatory property of the unperturbed cell, but rather it is a property of the interaction between the cell and the oscillatory inputs [105]. Hence, it is not clear whether and under what conditions subthreshold resonance persists in the presence of other types of periodic inputs with non-sinusoidal waveforms.

**Figure 4:**
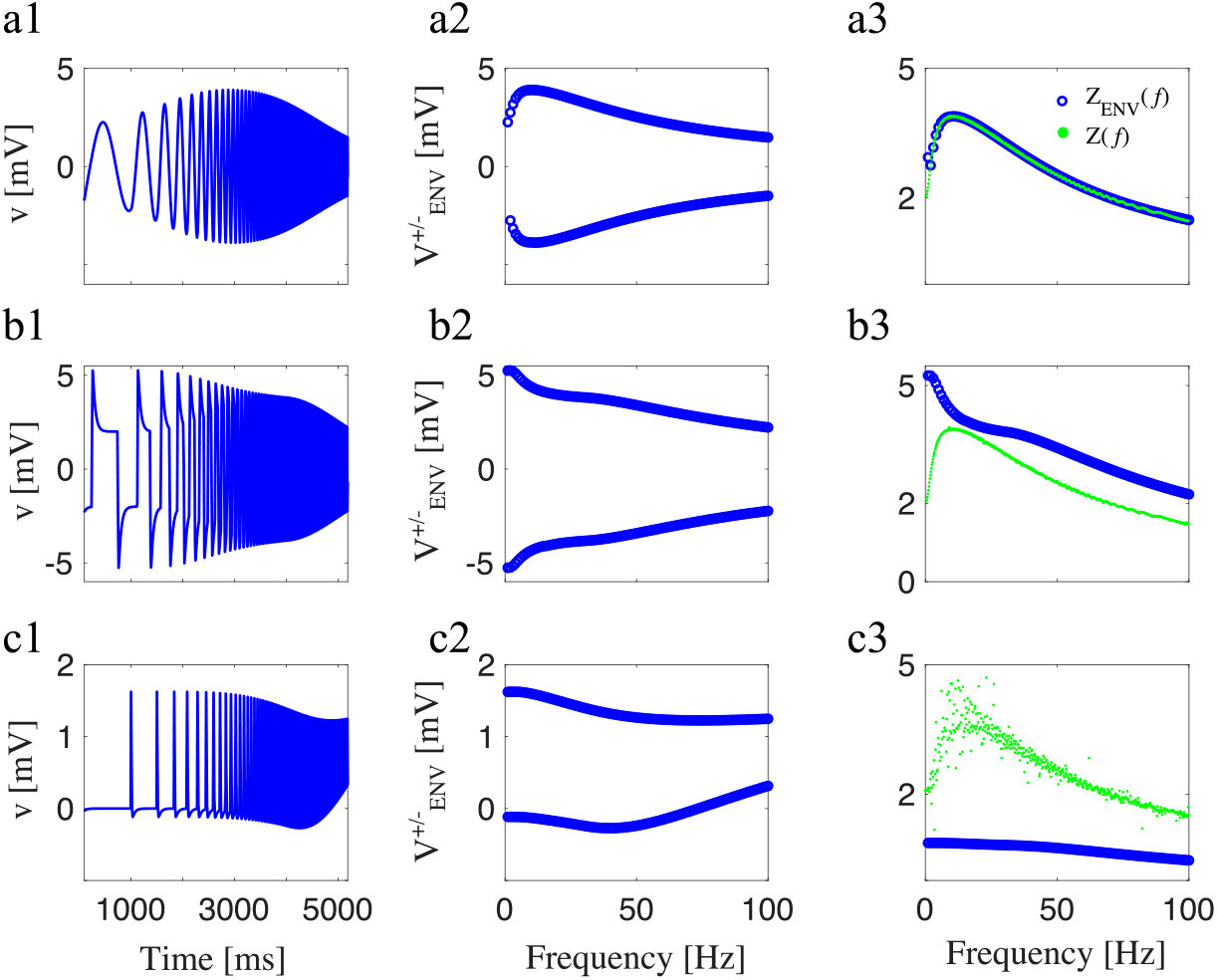
Neuronal response upon application of three different inputs (linear model, overshoot). (a) Sinusoidal chirp. (b) Square-wave chirp. (c) Synaptic-like chirp. (a1,b1, and c1) voltage traces. (a2, b2, and c2) Voltage-response envelopes in the frequency-domain. (a3, b3, and c3) *Z*(*f*) (frequency-content) and *Z_ENV_*(*f*) (envelope) impedance. We used the following parameter values: *C* = 1, *g*_L_ = 0.25, *g*_1_ = 0.25, *τ*_1_ = 100 ms, and *A*_in_ = 1 (same model as in Fig. 3a).

**Figure 5:**
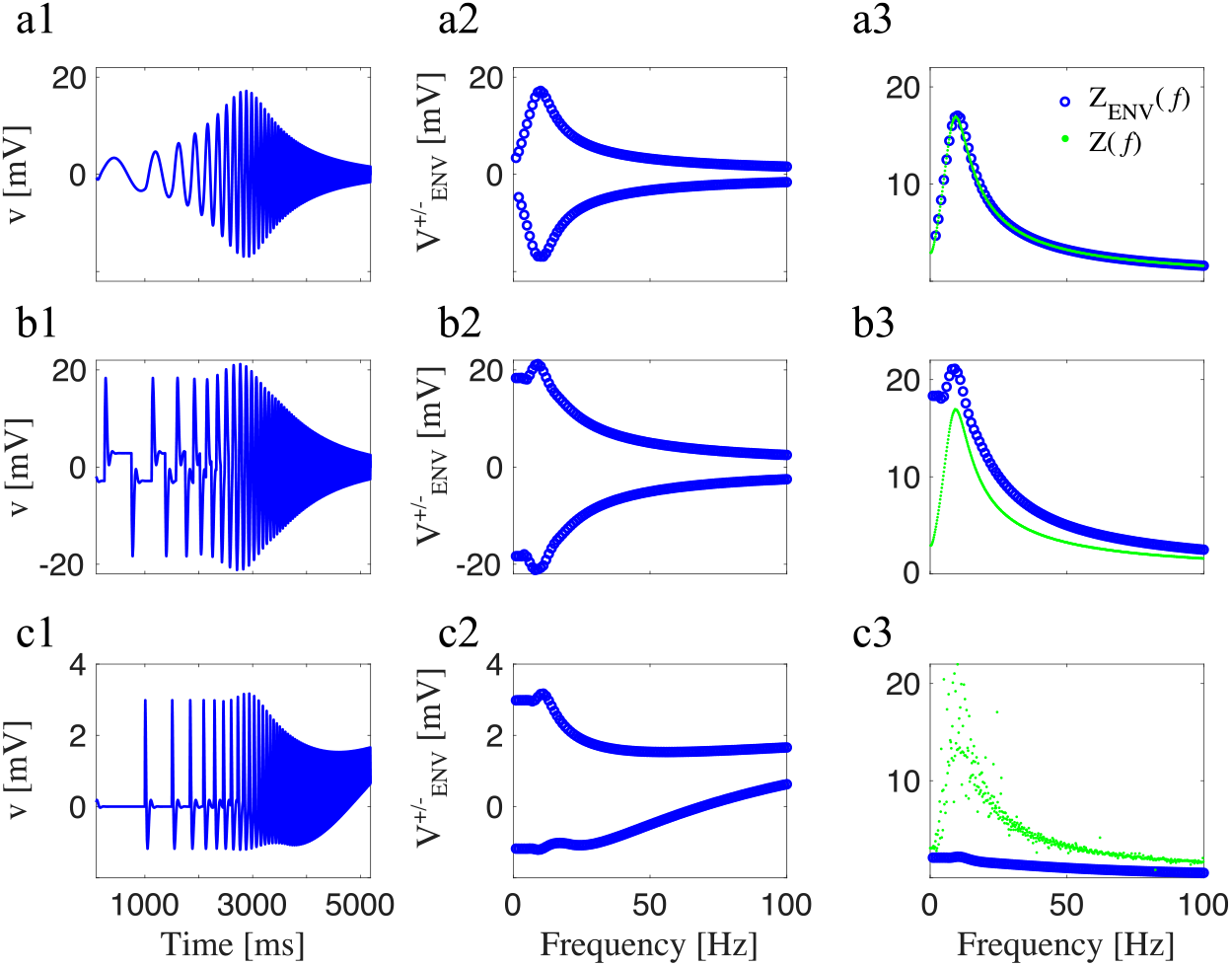
Neuronal response upon application of three different inputs (linear model, subthreshold oscillations). (a) Sinusoidal chirp. (b) Square-wave chirp. (c) Synaptic-like chirp. (a1,b1, and c1) voltage traces. (a2, b2, and c2) Voltage-response envelopes in the frequency-domain. (a3, b3, and c3) *Z*(*f*) (frequency-content) and *Z_ENV_*(*f*) (envelope) impedance. We used the following parameter values: *C* =1, *g*_L_ = 0.05, *g*_1_ = 0.3, *τ*_1_ = 100 ms, and *A*_in_ = 1.

In principle, there are various metrics one could use to characterize the frequency response profiles of neurons (and dynamical systems in general). The impedance *Z*-profile (curve of the impedance amplitude as a function of the input amplitude) measures the signal frequency content (Fig. 4-a3, green). The upper and lower envelope profiles 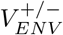 (Fig. 4-a2) capture the stationary peaks and troughs of the voltage response, respectively as a function of the input frequency. The peak profiles, in particular, are a relevant quantity since spikes are expected to occur at the response peaks as the input amplitude crosses threshold (the voltage response to this amplitude crosses the voltage threshold). The envelope impedance *Z_ENV_* profiles (Fig. 4-a3, blue), consisting of the stationary peak-to-trough amplitude normalized by the input amplitude as a function of the input frequency and serves to connect and compare between the two previous profiles.

For sinusoidal inputs, the *Z*-profile in response to chirp inputs typically coincides with the *Z_ENV_*-profile computed by using sinusoidal inputs of a constant frequency (over a range of input frequencies). This remains true for the sinusoidal chirp-like input we use here (Fig 4-a). It is always true for linear systems [63, 64] and certain nonlinear systems (e.g., [125]). In other words, the frequency content of the voltage response (green) is reflected by the voltage upper and lower envelope response profiles and the response to non-stationary chirp-like inputs coincides with the stationary response to sinusoidal inputs of a single frequency. This is a direct consequence of the fact that the input changes are gradual.

#### 3.2.2 Communication of the preferred frequency responses from the sub- to the supra-threshold regimes

The frequency-dependent suprathreshold response patterns to periodic inputs result from the interplay of the frequency-dependent subthreshold voltage responses to the same inputs and the spiking mechanisms. The subthreshold resonant frequency is communicated to the suprathreshold regime when neurons selectively fire action potentials in response to oscillatory inputs only at frequencies within a small enough range around the subthreshold resonant frequency. This type of evoking resonance can be obtained for input amplitudes sightly above these producing only subthreshold responses, for example for neurons for which the spiking response to oscillatory inputs can be thought of as spikes mounted on the corresponding subthreshold responses. Evoked spiking resonance captures a selective coupling between the oscillatory input and firing, and it has been observed experimentally and theoretically [69, 126] and the underlying dynamic mechanisms have been investigated in detail [127]. A related measure of the communication of the subthreshold resonant frequency to the suprathreshold regime is that of firing rate resonance [63] where the firing rate in response to oscillatory inputs peaks at or within a small range around the subthreshold resonance frequency. We note that subthreshold resonance does not necessarily imply evoked spiking resonance [69], evoked spiking resonance may be observed as the input amplitude crosses threshold, but lost for higher input amplitudes [127], the firing rate (or spiking frequency) at the firing rate (evoked) resonant frequency band is not necessarily the same as that frequency band [63, 126, 127], and evoked spiking resonance may be occluded in the presence of spontaneous firing, a situation likely to occur *in vivo*. When the spontaneous (or intrinsic) firing frequency is relatively regular, the associated time scale may dominate over the subthreshold resonant time scale and determine the firing rate resonant frequency [63]. A third form of preferred frequency response to oscillatory inputs is the so-called output spiking resonance [127] where the spiking frequency response to oscillatory inputs remains within a relatively narrow range independently of the input frequency range. The output spiking resonant frequency and the subthreshold resonant frequency are not necessarily the same, but the mechanisms that give rise to both are dynamically related [127].

While there is no guarantee that subthreshold resonance implies any of the various types of supra-threshold resonance, the communication of the resonant frequency to the suprathreshold regime is favored when the neuron’s upper envelope profile 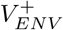 exhibits a peak at the subthreshold resonant frequency (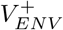 resonance). For the examples in Figs. 4-a and 5-a, the models, supplemented with a voltage threshold for spike generation and a reset mechanism, will exhibit evoked spiking resonance in response to sinusoidal inputs at the subthreshold resonant frequency band, and this is well captured by both the *Z* and *Z_ENV_* profiles. However, while this remains true for a larger class of systems, we note that this is not necessarily the case for nonlinear systems exhibiting, for instance if the 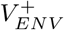 and 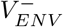 are asymmetric with respect to the equilibrium voltage [128]. For example, a cell that is an upper envelope low-pass filter (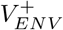 is a decreasing function of the input frequency), but a lower envelope band-pass filter (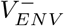 has a trough at an intermediate input frequency) will show a peak in the impedance profile *Z* and therefore will be considered resonant, but this will not necessarily be reflected in the spiking response since the lower frequencies will be communicated better to the spiking regime than the intermediate frequencies as the input amplitude increases above threshold, in particular, these within the subthreshold resonant frequency band.

This together with our discussion in the previous section raises the question of whether the responses of resonant cells to non-sinusoidal periodic inputs will also show a preferred frequency response in the resonant frequency band and whether the *Z*- and *V_ENV_*-profiles exhibit the same filtering properties. This has implications for the frequency-dependent supra-threshold response patterns to periodic inputs since the communication between neurons occurs via synaptic interactions, which exhibit abrupt changes as compared to the sinusoidal inputs, raising in turn the possibility of a competition between *Z*- and *V_ENV_*-profiles in determining the spiking frequency filtering properties.

### 3.3 Resonant cells do not necessarily show 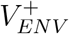 resonance in response to chirp-like square-wave inputs

Figs. 4-b1 and -b2 illustrate that (*Z*) resonant cells (see Fig. 4-a) may not exhibit envelope bandpass filter in responses to square-wave inputs. The 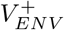 resonant response for sinusoidal inputs (Figs. 4-a1 and -a2) is lost for square-wave inputs and, consequently, these inputs would produce spiking activity preferentially at the lowest frequencies (no evoked spiking resonance) for input current amplitudes above threshold. The absence of 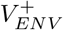-resonance does not imply the absence of *Z*-resonant frequency content. In fact, the power spectra for the responses to sinusoidal and square-wave inputs (Fig. 4-b3, green) are very similar to the power spectra for sinusoidal inputs (Figs. 4-a3, green) and all show *Z*-resonance (see schematic explanation in Fig. S1). However, this *Z*-resonance is not reflected in the *V* response and therefore it does not have a direct effect on the communication of the subthreshold frequency content to the spiking regime.

Figs. 5-b1 and -b2 shows a representative case where 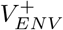 resonance is still present for square-wave inputs, but the resonance amplitude 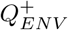 (defined as the quotient of the values of 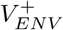 at the peak and at *f* = 0) is very small as compared to 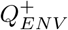 in response to sinusoidal inputs (Fig. 5-a2). In these cases, the subthreshold resonant frequency will be communicated to the spiking regime, but only for a small range of input amplitudes as compared to the responses to sinusoidal inputs (Fig. 5-a), above which spiking would occur for the lowest frequencies. The frequency content of the voltage response (Fig. 5-b3, green) is not reflected by the 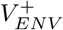 and 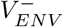 response profiles (Fig. 5-b2) and consequently by the *Z_ENV_* profiles (Fig. 5-b3, blue).

The main difference between the two cases presented in Figs. 4 and 5 is the type of autonomous transient dynamics of the two cells. For the parameter values used in Fig. 4 the equilibrium for the isolated cell is a stable node (real eigenvalues, no intrinsic damped oscillations) and the cell displays overshoot transient responses to input changes (e.g., Fig. 2-a), while for the parameter values used in Fig. 5, the equilibrium for the isolated cell is a stable focus and the cell displays damped oscillations in response to input changes (e.g., Fig. 2-b). Biophysically, the transition from stable nodes to stable foci is associated with an increase in the levels of the amplifying currents. In Fig. 5, this is reflected as a decrease in the linearized conductance *g_L_*, which contains information about fast amplifying currents such as *I_Nap_* present in the original biophysical models [63, 64].

The persistence of 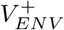 resonance in cells having stable foci is due to a combination of the (damped) oscillatory response after the abrupt input increase/decrease and summation. More specifically, the response to the abrupt input changes (square-wave or synaptic) has two regimes: a relatively large amplitude response, reflecting the biophysical amplification levels, and a smaller amplitude response reflecting the stability properties of the equilibrium *V_eq_*. The location of the voltage response *V* right before the arrival of the input from the next cycle determines the response amplitude to this input and this location depends on the stability properties of *V_eq_*. When *V_eq_* is a node, the voltage response *V* decreases below *V_eq_* immediately after the abrupt input change, and then returns to *V_eq_*. When the input from the next cycle arrives, *V* is below *V_eq_*. In contrast, when *V_eq_* is a focus, the oscillatory voltage response may be above *V_eq_* when the input from the next cycle arrives, and therefore, because it starts at a higher value, *V* reaches higher values. This depends on the frequency of the damped oscillations and the input frequency. If the input frequency is too low, then the damped oscillations die out before the next input arrives, while if the input frequency is high enough, then the damped oscillations are close to their first peak. If the input frequency is higher, then the value that *V* has when the next input arrives is lower because is further away from the first peak. For still higher input frequencies, the standard summation takes over.

Figs. S2 and S3 show similar results for the nonlinear conductance-based *I_h_* + *I_Nap_* model (the linear models used for Figs. 4 and 5 can be considered as linearized versions of this *I_h_* + *I_Nap_* model). For the lower levels of the *I_Nap_* conductance *G_p_* the *I_h_* + *I_Nap_* shows no 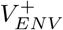 resonance, while 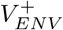 resonance persists for the higher levels of *G_p_*, consistent with the transition of the equilibrium from a stable node to a stable focus.

### 3.4 Dependence of 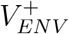 resonance in response to synaptic-like input currents on the current sign and the cell’s intrinsic properties

#### 3.4.1 *Z*-resonant cells do not necessarily show 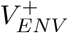 resonance in response to excitatory synaptic-like inputs currents

As for the square-wave inputs described above, 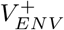 is absent when *V_eq_* is a node (Figs. 4-b1 and -b2) and present when *V_eq_* is a focus (Figs. 5-b1 and -b2), but with a smaller 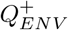 than the response to sinusoidal inputs (Fig. 5a). Also similarly to square-wave inputs, the absence of 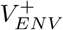-resonance does not imply the absence of *Z*-resonant frequency content; The power spectra of the responses to sinusoidal, square-wave and synaptic-like inputs are very similar and all show *Z*-resonance (compare Figs. 4-b3 and -5-b3, green, with Figs. 4-a3 and -5-a3, green), and therefore this (*Z*-) resonance may have no direct effect on the communication of the subthreshold frequency content to the spiking regime (since the frequency-dependent properties that govern the generation of spikes are captured by 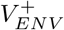 profiles and not on the frequency content captured by the *Z*-profiles).

In contrast to the responses to square-wave inputs, both the 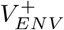 and 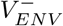 responses to excitatory synaptic-like inputs exhibit a trough before increasing due to summation, which is more pronounced in Fig. 4c (*V_eq_* is a node) than in Fig. 5c (*V_eq_* is a focus). The generation of these troughs are the result of the interplay of the accumulation of synaptic inputs and the intrinsic properties of the cell reflected by the transient responses to individual inputs (overshoots, damped oscillations), and occur at a different frequency than the *Z*-resonant frequency. More specifically, for the lower frequencies in Fig. 4-c, *V* exhibits a sag before returning to a vicinity of *V_eq_*. The value *V* reaches before the arrival of the next input serves as the initial condition for the next cycle. As the input frequency increases, these initial conditions are lower than for the previous cycles since the periods decrease, and therefore *V* returns to an even lower value after the synaptic input wears off. As the input frequency increases further, standard summation takes over and the combination of summation and the higher frequency input creates the high-pass filter *V_ENV_* patterns with an amplitude that decreases with frequency. This phenomenon is watered down when *V_eq_* is a focus because the amplification associated to the presence of damped oscillations as compared to overshoots causes the voltage response troughs at every cycle to reach lower values when *V_eq_* (Fig. 5c) is a focus than when *V_eq_* is a node (Fig. 4c).

Figs. S2 and S3 show similar results for the nonlinear conductance-based *I_h_* + *I_Nap_* model.

#### 3.4.2 Z-resonant cells show 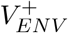 resonance in response to inhibitory synaptic-like inputs currents in a *V_eq_*-stability-dependent manner

For inhibitory synaptic-like input, the 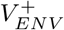 and 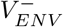 responses are qualitatively inverted images of the ones described above. For linear cells, in particular, the responses for excitatory and inhibitory synaptic-like inputs are symmetric with respect to *V_eq_* (= 0). The most salient feature is the presence of a peak in both 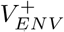 and in 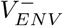 (Figs. S4-a1 and -b1 and Figs. S4-a2 and -b2), indicating the occurrence of 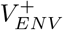 resonance at a frequency, which is different from the *Z*-resonant frequency (Figs. S4 c1 and c2). The mechanism of generation of this 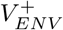 band-pass filters is similar to the one described for the troughs in excitatory synaptic-like inputs and involves a combination of summation and intrinsic properties of the cell, reflected in the properties of the transient response of the cells to individual inputs. More specifically, the summation acts as a low-pass filter and the effects of the transient responses to individual neurons, associated with the presence of *Z* resonance, act as a high-pass filter. We emphasize that the 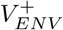 and *Z* resonances are significantly different. We also emphasize that 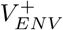 resonance is not significant when *V_eq_* is a focus (Fig. S4-b2) since the amplification associated to the presence of damped oscillations referred to above obstructs the envelope high-pass filtering component.

Figs. S5 and S6 show similar results for the nonlinear conductance-based *I_h_* + *I_Nap_* model.

#### 3.4.3 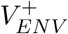 low- and high-pass filtering properties of synaptic-like currents

The responses to synaptic-like inputs are affected by the summation effect, which depends on the synaptic decay time *τ*_Dec_. Figs. S7 and S8 illustrate the transition of the response (middle and right panels) to excitatory synaptic-like inputs (left panels) for representative values of *τ*_Dec_ (including these used in Figs. 4-c and 5-c). In all cases, the frequency content measured by the impedance *Z* (Figs. S7 and S8, green) remains almost the same. The summation effects, which increases as *τ*_Dec_ increases, strengthens the low-pass filter properties of the *Z_ENV_* response. The results for inhibitory synaptic-like inputs are symmetric to these in Figs. S7 and S8 with respect to *V_eq_* (= 0) (not shown), and therefore, increasing values of *τ*_Dec_ strengthens the low-pass filter properties of *Z_ENV_*.

### 3.5 The autonomous transient dynamic properties are responsible for the poor upper envelope 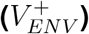 resonance (or lack of thereof) exhibited by (*Z*-) resonant cells in response to non-sinusoidal chirp-like input currents

As discussed above, the differences between the 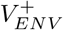 response patterns to square-wave/synaptic-like and sinusoidal inputs and the differences between the 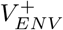 response patterns to different types of synaptic-like inputs (excitatory, inhibitory) are due to the different ways in which individual cells transiently respond to abrupt and gradual input changes, which operate at every cycle. Both and the corresponding sinusoidal inputs share the primary frequency component determined by the period (see Fig. S1). However, the sinusoidal input is gradual and causes a gradual response without the prominent transients (overshoots and damped oscillations) observed for the square-wave input, which together with the summation phenomenon produces *V_ENV_* peaks. The responses to square wave inputs, in particular for the lower frequencies, reach a steady-state value as the responses to sinusoidal inputs do, but in contrast to the latter, the voltage envelope for the former is determined by the transient peaks. These transients appear to be “getting in the way” of the voltage response to produce *V_ENV_* resonance. However, they are not avoidable. In fact, overshoots in non-oscillatory systems are an important component of the mechanism of generation of resonance in response to sinusoidal inputs [65, 105] as are damped oscillations.

For comparison, Fig. S9 shows the responses of a passive cell (*g*_1_ = 0) to the three types of inputs. Passive cells exhibit neither overshoot nor damped oscillatory transient responses to abrupt input changes, but monotonic behavior. As expected, this cell does not exhibit resonance in response to oscillatory inputs, but a low-pass filter response in both *Z* and *V_ENV_* (Figs. S9-a). The envelope response to square-wave inputs is also a low-pass filter (Fig. S9-b), though it decays slower with increasing values of the input frequency since, because of the waveform, the squarewave input stays longer at its maximum value than the sinusoidal input at each cycle. In contrast to Figs. 4 and 5, the cell’s response to synaptic-like inputs is a *V_ENV_* high-pass filter (Fig. S9-c) due to summation and the lack of interference by the transient effects. Fig. S10 shows similar results for a two-dimensional linear model with a reduced value of the negative feedback conductance *g*_1_ where overshoots and damped oscillations are not possible. The analogous results for inhibitory synaptic-like inputs are presented in Figs. S4 (rows 3 and 4).

### 3.6 Current- and conductance-based synaptic-like inputs produce qualitatively different voltage responses and synaptic currents

The synaptic-like inputs considered so far are additive current inputs. However, realistic synaptic currents involve the interaction between the synaptic activity and the postsynaptic voltage response. In biophysical models, the synaptic currents terms consist of the product of synaptic conductances and the voltage-dependent driving force (Eq. 3). Because the voltage response contributes to the current that produces this response, the frequency-dependent response profiles for current- and conductance-based inputs may be qualitatively different.

#### 3.6.1 *Z*-resonant cells do not show 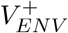 resonance in response to conductance-based excitatory synaptic-like inputs, but they do show troughs in the 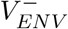 response

Figs. 6 and 7 show representative comparative examples for the two *Z*-resonant cells (in response to sinusoidal inputs) discussed in Figs. 4 (stable node) and 5 (stable focus), respectively. For the parameter values corresponding to Fig. 4 (no 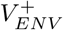 resonance in response to current-based synaptic-like inputs, Fig. 6-b, blue), the conductance-based synaptic current shows a peak in the upper envelope (Fig. 6-a2), but not in the voltage response (Fig. 6-b, red), which, instead, shows a trough as for the current-based synaptic input (Fig. 6-b, blue). In spite of the similarities between the two profiles, the *Z_ENV_* profile for the conductance-based input shows a peak (Fig. 7-c1, red), but this peak does not reflect a true *Z_ENV_* preferred frequency response.

**Figure 6:**
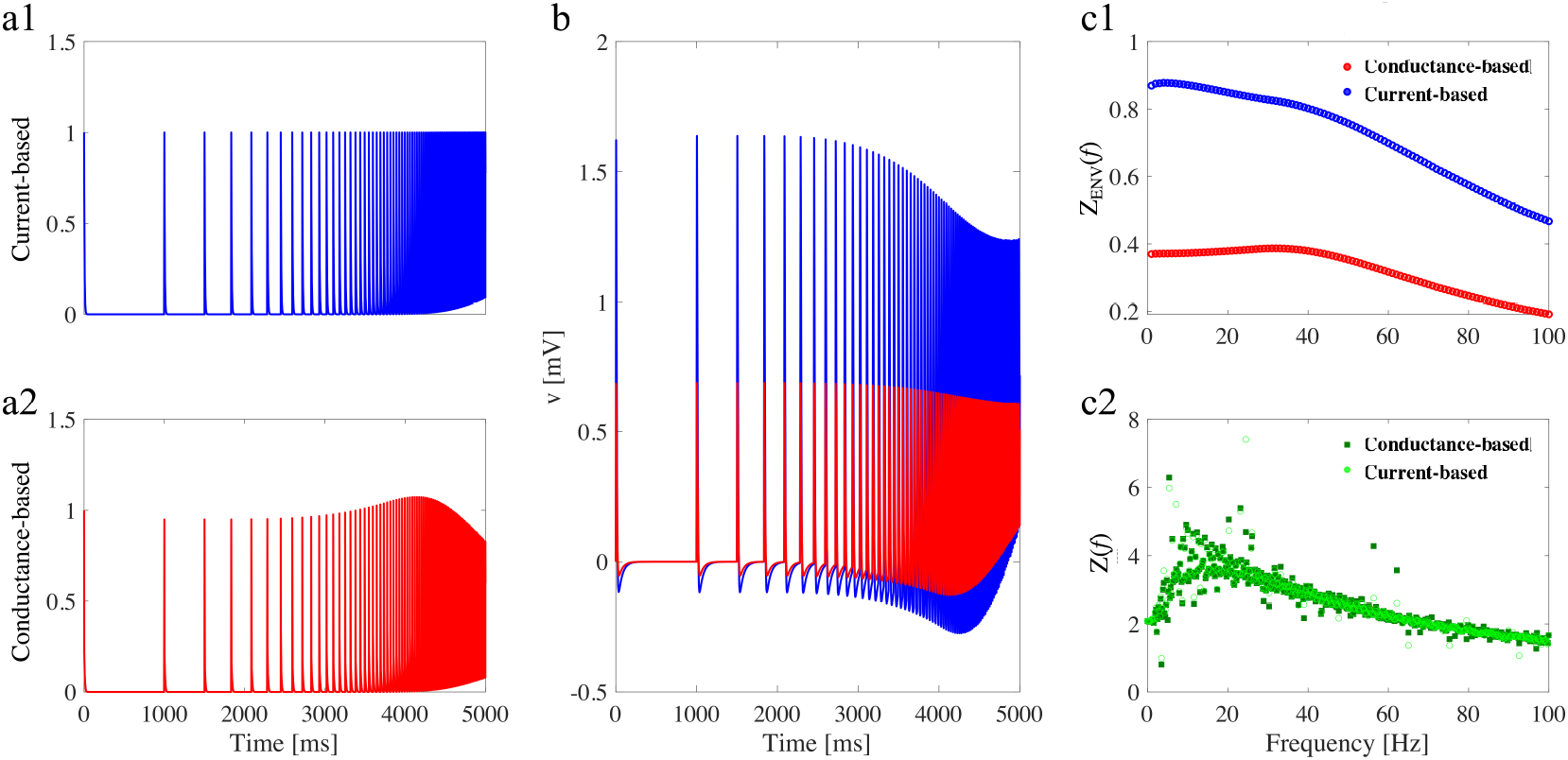
Comparison between conductance-based and current-based inputs (linear model, overshoot). (a) Input currents (see Eqs. 1 and 3). (a1) Current-based input. (a2) Conductance based input. (b) Voltage traces (same colors as in a1 and a2). (c1) *Z_ENV_*(*f*) (envelope) impedance. (c2) *Z*(*f*) (frequency-content). We used the following parameter values: *C* =1, *g*_L_ = 0.25, *g*_1_ = 0.25, *τ*_1_ = 100 ms, and *A*_in_ = 1, same model as in Fig. 4.

**Figure 7:**
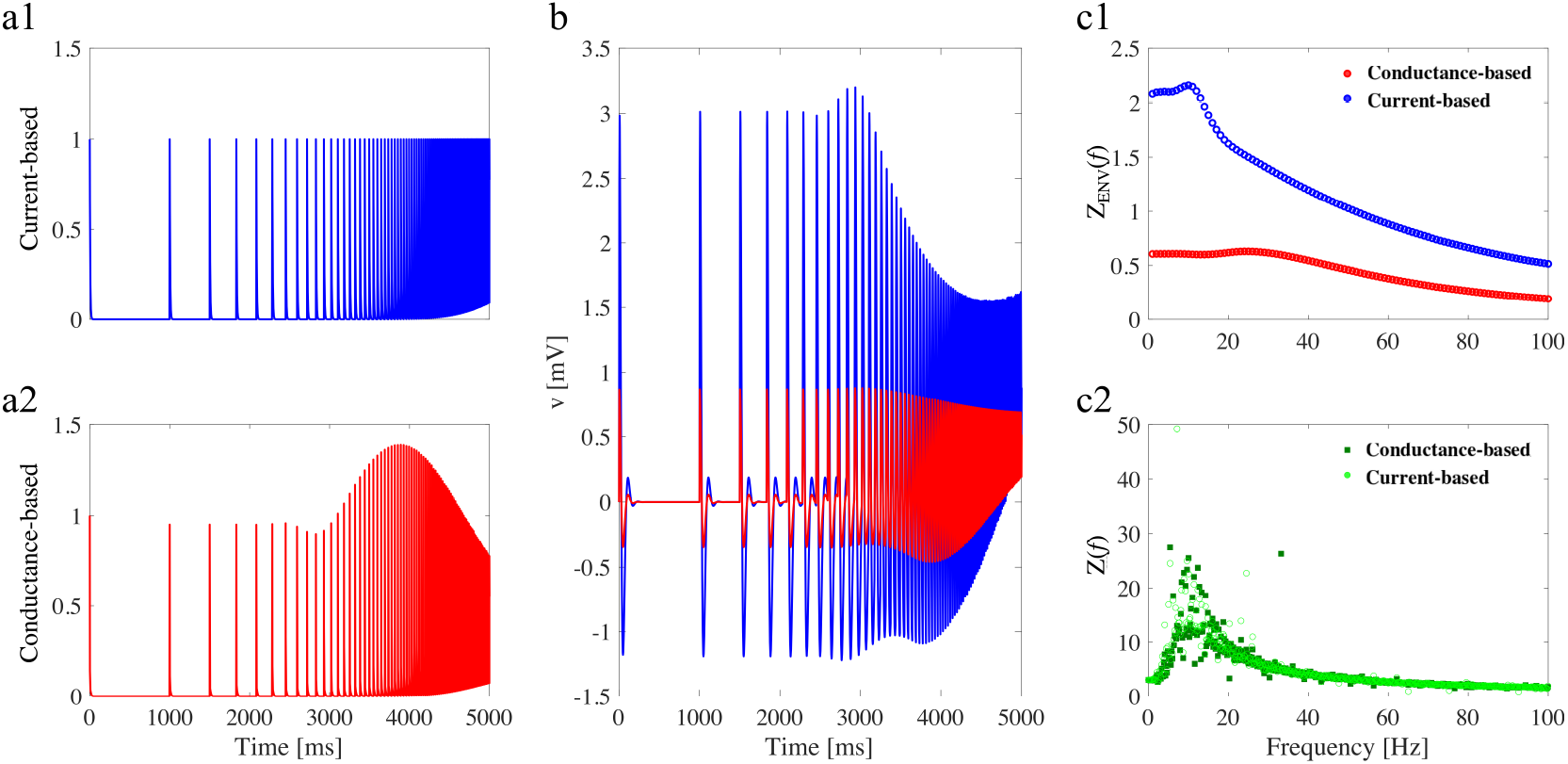
Comparison between conductance-based and current-based inputs (linear model, subthreshold oscillations). (a) Input currents (see Eqs. 1 and 3). (a1) Current-based input. (a2) Conductance based input. (b) Voltage traces (same colors as in a1 and a2). (c1) *Z_ENV_*(*f*) (envelope) impedance. (c2) *Z*(*f*) (frequency-content). We used the following parameter values: *C* =1, *g*_L_ = 0.05, *g*_1_ = 0.3, *τ*_1_ = 100 ms, and *A*_in_ = 1, same model as in Fig. 5.

For the parameter values corresponding to Fig. 5 (mild 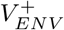 resonance in response to current-based synaptic-like inputs, Fig. 7-b, blue), the response to conductance-based synaptic inputs is similar to that in Fig. 6, but more amplified. In particular, there is no 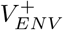 resonance in response to conductance-based synaptic-like input (Fig. 7-b, red). In both cases, the cells show *Z* resonance (Figs. 6-c2 and 7-c2).

For comparison, Fig. S11 shows the result of repeating the protocols used above for a passive cell (*g*_1_ = 0, same as Fig. S9). The frequency response patterns are the standard *Z*- and *Z_ENV_* low-pass filters and the expected 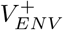 high-pass filter.

#### 3.6.2 *Z*-resonant cells show 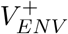 resonance in response to conductance-based inhibitory synaptic-like inputs

Figs. S12, S13, and S14 shows the result of repeating the protocols described above (Figs. 6, 7 and S11, respectively) using synaptic-like inhibitory conductance-based inputs. The *Z*- and *Z_ENV_*-profiles are qualitatively similar, except for the relative magnitudes of the synaptic- and conductance-based *Z_ENV_* that are inverted. The 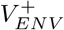 profile shows a significant (resonant) peak when the cell has a node (Fig. S12-b), which is almost absent when the cell has a focus (Fig. S13-b), but *Z_ENV_* has a peak when the cell is a focus (Fig. S13-c1), while it is a low-pass filter when the cell has a node (Fig. S12-c1).

Together, these results and the result from the previous section shows that the frequency content (in terms of *Z*) of resonant cells persists in response to synaptic-like current- and conductance-based inputs, but the 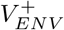 responses are different between synaptic-like current- and conductance-based inputs, and these differences depend on whether the cell has a node or a focus and whether the synaptic-like input is excitatory or inhibitory. Of particular interest are the peaks in the conductance-based synaptic inputs (Figs. 6-a2 and 7-a2, red).

### 3.7 Amplitude variability in response to chirp-like inputs with arbitrarily distributed cycles results from the transient response properties of the autonomous system

In the previous sections we used discretely changing frequencies (chirp-like or, simply, chirps, see Section 2.2.1) as a compromise between tractability and the ability to incorporate multiple frequencies in the same input signal, and we extended this type of inputs to waveforms with more realistic time-dependent properties. In all the cases considered so far, the chirp-like input cycles were “regularly” ordered in the sense that the input frequency monotonically increases with the cycle number (and with time). In this section, we move one step forward and consider chirp-like inputs where the cycles are arbitrarily ordered (see Section 2.2.2 and, Fig. 1-b) in an attempt to capture the fact that information does not necessarily arrive in a regularly ordered manner while keeping some structure properties (sequence of oscillatory cycles), which ultimately allows for a conceptual understanding of the responses.

Each trial consists of a permutation of the order of the cycles using the regularly ordered cycles as a reference. The regularly and arbitrarily ordered input signals have exactly the same cycles (one cycle for each frequency value within some range) and therefore the same frequency content. The corresponding responses are expected to have roughly the same frequency content as captured by the *Z*-profiles within the range of inputs considered. However, we expect the voltage responses to have different frequency-dependent *V* responses, captured by the peak-and-trough envelopes 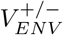. The differences between the *V* responses for two different inputs (different cycle orders) are due to the different ways in which the autonomous transient dynamics are activated across cycles for these inputs as the result of the transition between cycles. The values of the participating variables at the end of one cycle become the initial conditions for the subsequent cycle.

#### 3.7.1 Emergence of the amplitude variability

In Section 3.2.1 (Figs. 4-a and 5-a) we showed that (*Z*-) subthreshold resonance is well captured by the 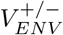 profiles in response to chirp-like sinusoidal inputs via the *Z_ENV_*-profiles (difference between the 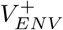 and 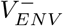 profiles). In Section 3.5 we argued that the transient response properties of the autonomous (unforced) cells (overshoots, damped oscillations, passive monotonic increase/decrease) are responsible for the (frequency-dependent) differences between the 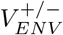 profiles and the *Z*-profiles in response to both square-wave and synaptic-like chirp-like inputs, and for the (frequency-dependent) differences among the 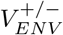 profiles in response to the three types of chirp-like inputs.

The ordered chirp-like input signals produced 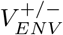 profiles with gradual amplitude variations along with the input frequency range (and a very small number of increasing and decreasing portions). The peaks and troughs for each frequency are determined by two parameters: the values of the variables at the beginning of the corresponding cycle and the duration of the cycle (the intrinsic properties of the cell are the same for all input frequencies), which in turn determines the initial values of the variables in the next cycle. The monotonic increase of the input frequency causes a gradual change in these parameters along the frequency axis, which in turn causes gradual changes in the 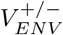 profiles.

Because of this dependence of the values of the variables at the beginning of each cycle with the values of these variables at the end of the previous cycle, we reasoned that the voltage response to chirp with arbitrarily distributed cycles in time will exhibit non-regularly distributed peak and troughs, leading to amplitude variability in the 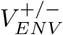 profiles, while producing at most minimal changes in the *Z* profiles (as compared to the responses to input signals with order cycles) within the frequency range considered. Moreover, this variability will depend on the type and properties of the autonomous transient dynamics of the participating cells. The arbitrary distribution in the order of the cycles in the input signal is achieved by considering one permutation of the regularly ordered signal (signal with regularly ordered cycles). The randomness in the input signals lies in the choice of a subset of all possible permutations for the considered trials.

Our results are presented in Figs. 8 and 9 for a cell exhibiting an overshoot (Fig. 8; stable node; same parameter values as in Fig. 4) and damped oscillations (Fig. 9; stable focus; same parameter values as in Fig. 5) in response to step-constant inputs. For sinusoidal and square-wave chirplike inputs, the output 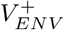 and 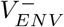 frequencies were computed as the differences between two consecutive troughs and two consecutive peaks, respectively, normalized so that the resulting frequencies have units of Hz. The 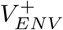 and 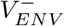 profiles consist of the sequence of maxima and minima for each frequency (dots superimposed to the *v* time courses in the left columns) and include the damped oscillations for the lower frequencies (e.g., Fig. 9-b1). For the synaptic-like chirp inputs, we used the 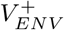 and 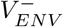 profiles consisting of the sequence of maxima and minima for each input frequency and do not include the damped oscillations for the lower frequencies (e.g., shown in Fig. 9-c1, but not present in Fig. 9-c2).

**Figure 8:**
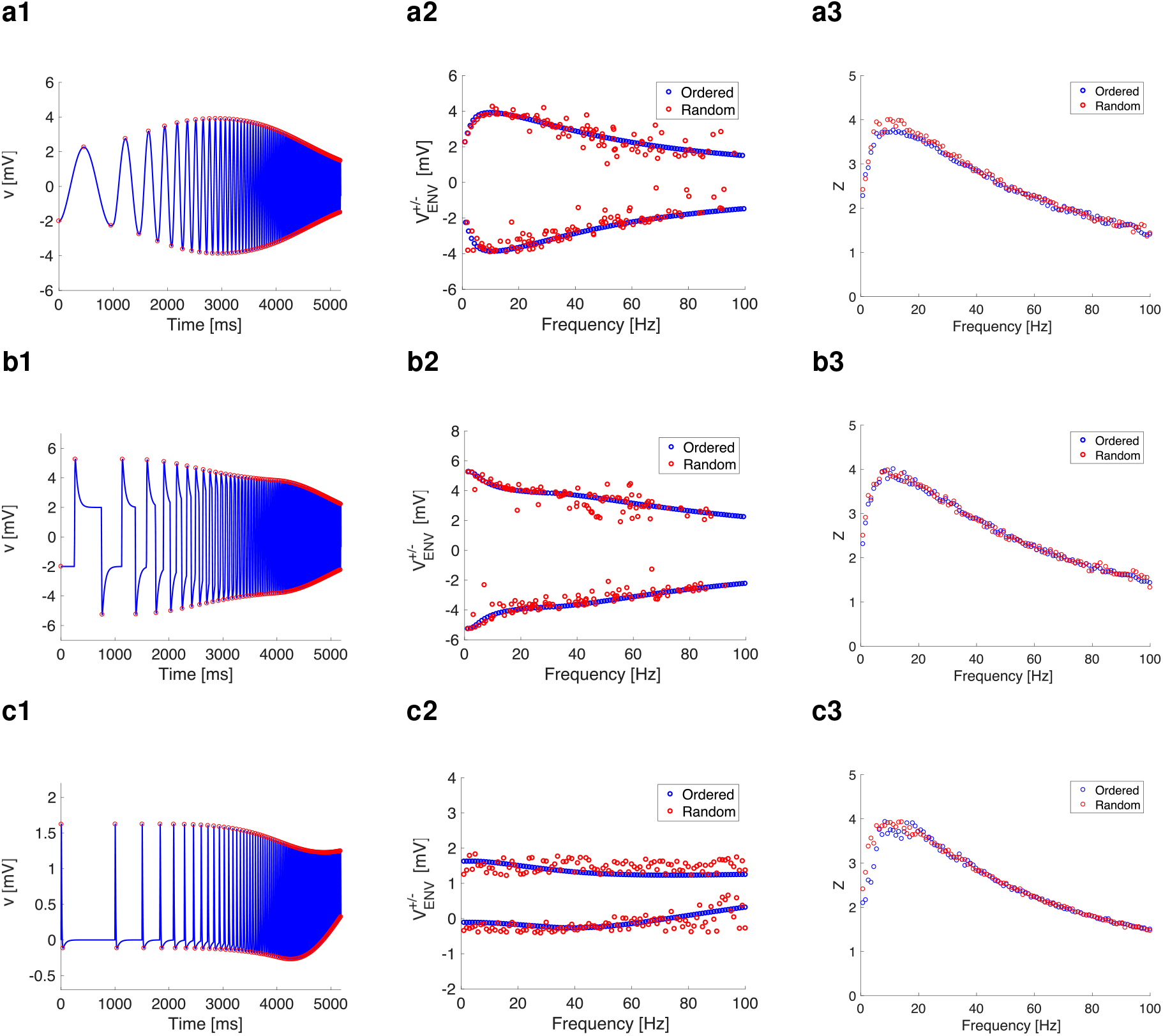
Comparison of neuronal response between ordered and random input (linear model, overshoot). (a) Sinusoidal chirp. (b) Square-wave chirp. (c) Excitatory synaptic-like chirp. (a1, b1, and c1) voltage traces with peaks and troughs marked by red circles. (a2, b2, and c2) Voltage-response envelopes in the frequency domain (blue is ordered input; red is random input as in Fig. 1b). (a3, b3, and c3) *Z*(*f*) (frequency-content) for ordered and shuffled inputs. In this 2D linear model, we used the following parameter values: *C* = 1, *g*_L_ = 0.25, *g*_1_ = 0.25, *τ*_1_ = 100 ms, and *A*_in_ = 1. Same model as in Fig. 4.

**Figure 9:**
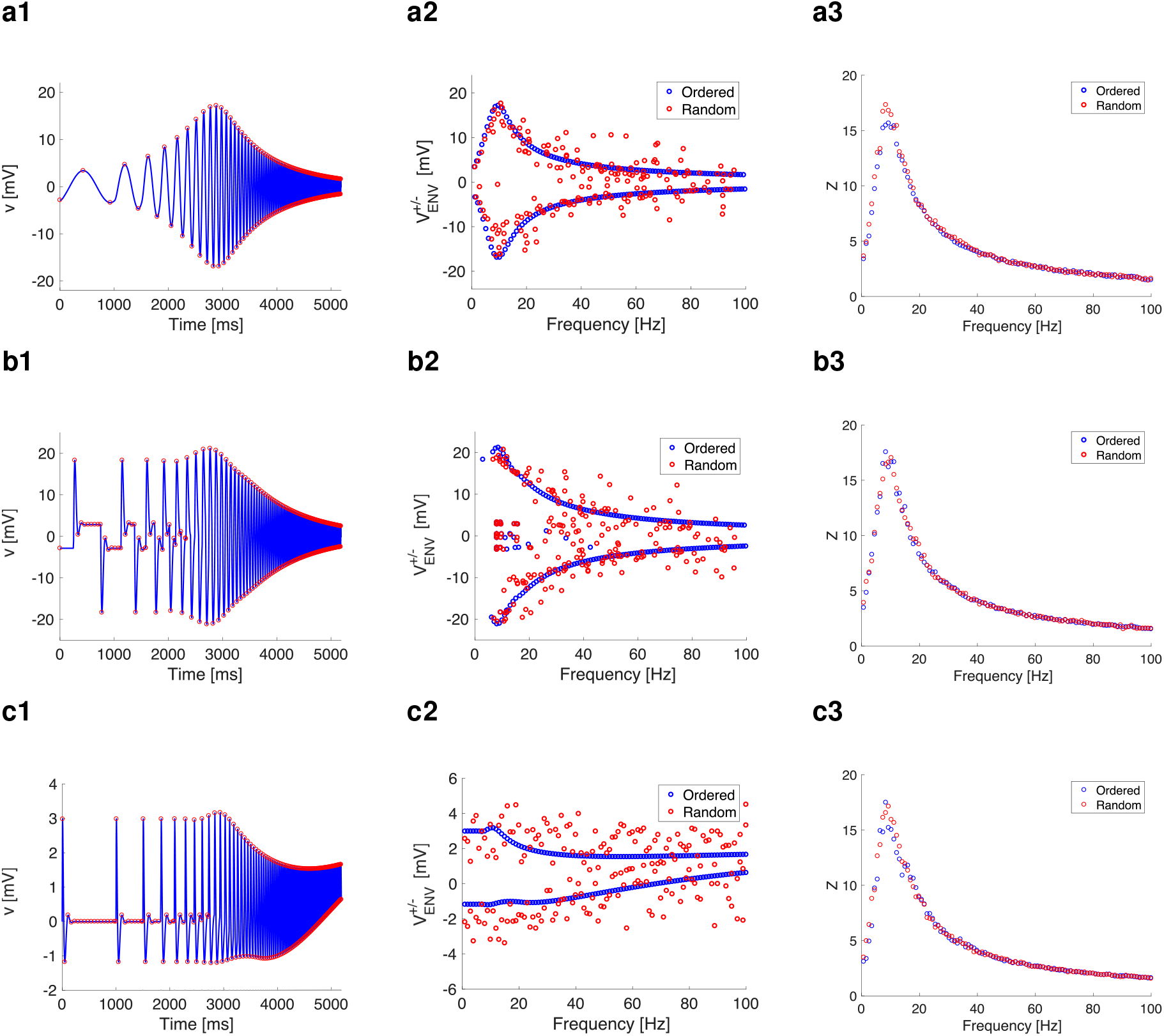
Comparison of neuronal response between ordered and random input (linear model, subthreshold oscillations). (a) Sinusoidal chirp. (b) Square-wave chirp. (c) Excitatory synaptic-like chirp. (a1, b1, and c1) voltage traces with peaks and troughs marked by red circles. (a2, b2, and c2) Voltage-response envelopes in the frequency domain (blue is ordered input; red is random input as in Fig. 1b). (a3, b3, and c3) *Z*(*f*) (frequency-content) for ordered and shuffled inputs. In this 2D linear model, we used the following parameter values: *C* = 1, *g*_L_ = 0.05, *g*_1_ = 0.3, *τ*_1_ = 100 ms, and *A*_in_ = 1. Same model as in Fig. 5.

We use the (regularly changing) responses to inputs with regularly ordered cycles (blue) as a reference for the variability of the responses to arbitrarily ordered cycles (red). In both Figs. 8 and 9 the responses to the input with randomly ordered cycles (red) have random amplitudes organized around (sinusoidal and square-wave; panels a and b) or in a vicinity (synaptic-like panel c) of the responses to regularly ordered cycles. The amplitude response variability is stronger for higher frequencies than for the lower frequencies, since the responses for the former are more affected by changes in the initial conditions at the corresponding cycles. Importantly, the variability is stronger for cells having stable foci (exhibiting transient damped oscillations; Fig. 9) than for cells having stable nodes (exhibiting transient overshoots; Fig. 8), reflecting the higher complexity of the latter cells’ autonomous part. In all cases considered, the *Z* profiles remain almost unaffected by the order of cycles within the input frequency range. For comparison and completeness, Figs. S15 and S16 show similar graphs for a passive cell and synaptic-like inhibition, respectively. An important observation common to the responses of the three types of cells to synaptic-like inputs is the identification of the summation effects in the generation of 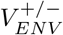 resonances. For example, Fig. S15-c2 (passive cell) shows that the summation effect in response to regularly ordered cycles (blue) disappears in the responses to randomly ordered cycles (red). Furthermore, the 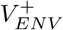 resonance in response to regularly ordered synaptic-like inhibitory inputs (Fig. S16-a2, blue) also disappears in the responses to randomly ordered synaptic-like inhibitory inputs (Fig. S16-a2, red).

Together these results show that the disruption of the regular order of a set of basic input signals, while the basic signals and their shapes remain unchanged, is translated into the amplitude variability of the response as compared to the responses to the regularly ordered sequence of signals, and this variability results from the properties of the transient dynamics of the (unforced) cells receiving the input.

#### 3.7.2 Dependence of the amplitude response distribution variance with the cycle frequency and the properties of the receiving cell

Here we focus on synaptic-like inputs since they are the most realistic signals cells receive and we considered both current- and conductance-based synaptic-like inputs. We used the model (1)-(2) for current-based inputs and the model (3)-(4) for conductance-based inputs. We use the same input signal for both (*I*(*t*) = *S*(*t*)).

In order to quantify the variability of the voltage response envelopes (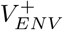 and 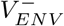) to arbitrarily ordered chirp-like inputs we considered a number of trials (*N_trials_* = 100) and computed the cycle-by-cycle variance for the corresponding peaks 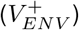 and troughs 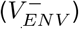. Our results for some representative cases are presented in Fig. 10. In all cases, 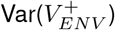 (blue) is less variable across frequencies than 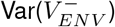 (red). The latter significantly increases for the higher frequencies. This high variability is associated with the phenomenon of summation observed in the regularly ordered cell. In other words, while summation is not observed in the responses to arbitrarily distributed cycles (e.g., Figs. 8-c, 9-c, and S15-c), it is translated into a high response variability. In Fig. 10-a, the transition from the P-cell (passive cell) to the N-cell (node cell) is due to a small increase in *g*_1_ and therefore the Var patterns are similar. The transition from the N-cell to the F-cell (focus cell) involves changes in both *g*_L_ and *g*_1_ in order to maintain *f_res_* within the same (small) range. Both 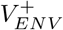 and 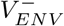 are significantly larger for the F-cell than for the N-cell, consistent with Figs. 8 and 9. The amplification of the initial portion of the transient response to constant inputs caused by differences in cell type is translated into a higher response variability. In Fig. 10-b, the transition from the P-cell to the N-cell to the F-cell is due to an increase only in *g*_1_ (at the expense of having values of *f_res_* distributed on a longer range than in panels a). The Var magnitudes are similar among the different cases. Together, these results reflect the fact that changes in the levels of the positive feedback effects (captured by the parameter *g_L_* in linear models) have stronger effects on the response variability than changes in the negative feedback effect (captured by the parameter *g*_1_).

**Figure 10:**
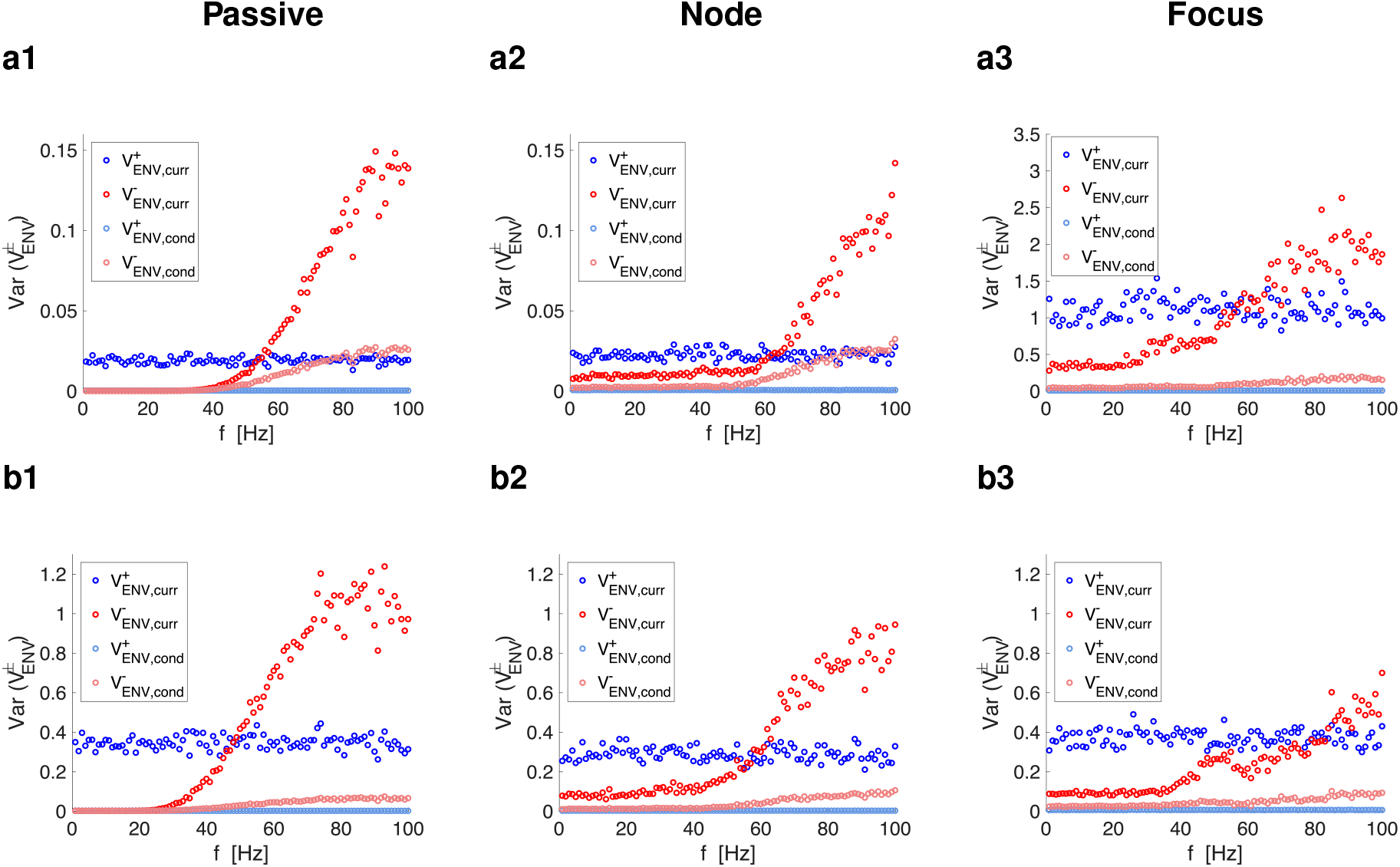
Peak and trough envelope (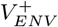 and 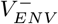) variability in response to synaptic-like chirp-like inputs with arbitrarily distributed cycles for current- and conductance-based models. We used the linear model (1)-(2). Each trial (*N_trials_* = 100) consists of a permutation of the cycle orders using as reference the ordered input patterns in Figs. 8-c to S15-c. The blue and red curves represent the variances across trials for 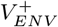 and 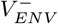 in response to synaptic-like current-based inputs. The light-blue and light-coral curves represent the variances across trials for 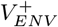 and 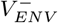 in response to synaptic-like conductance-based inputs. **Column 1.** Passive cells. **Column 2.** Node (N-) cells. **Column 3.** Focus (F-) cells. **a1.** *g_L_* = 0.25 and *g*_1_ = 0 (*f_nat_* = *f_res_* = 0). **a2.** *g_L_* = 0.25 and *g*_1_ = 0.25 (*f_nat_* = 0 and *f_res_* = 9). **a3.** *g_L_* = 0.05 and *g*_1_ = 0.3 (*f_nat_* = 8.1 and *f_res_* = 8). **b1.** *g_L_* = 0.1 and *g*_1_ = 0 (*f_nat_* = *f_res_* = 0). **b2.** *g_L_* = 0.1 and *g*_1_ = 0.2 (*f_nat_* = 0 and *f_res_* = 7). **b3.** *g_L_* = 0.1 and *g*_1_ = 0.8 (*f_nat_* = 12.3 and *f_res_* = 14). We used the additional parameter values: *C* = 1, *τ*_1_ = 100, *A_in_* = 1, *G_syn_* = 1, *E_syn_* = 1.

#### 3.7.3 The envelope response variabilities are stronger for current-than for conductance-based synaptic-like inputs

The 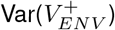 and 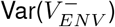 for the conductance-based synaptic-like inputs follow a similar pattern as these for the current-based inputs (Fig. 10, light-blue and light-coral), but the magnitudes for the former are lower than these for the latter inputs, consistent with the attenuation of the initial portion of the transient response to conductance-based constant synaptic inputs as compared to current-based constant synaptic inputs discussed in Section 3.1.2. These relationships persist when the 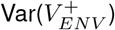 and 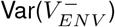 patterns are normalized by the amplitude of the response of the first synaptic-like input in the ordered patterns (a metrics that takes into account the effects of the differences in parameter values by the relative magnitude of their responses to the same input pattern).

### 3.8 Intrinsic oscillations evoked by Gaussian white noise may be lost or attenuated in response to synaptic-like inputs with arbitrarily ordered frequencies

#### 3.8.1 Oscillations (and lack of thereof) in response to synaptic-like chirp inputs

Fig. 11-a (blue) corresponds to the same parameter values as Figs. 8-c, 9-c, and S15-c. The most salient feature is the low-pass filter in the middle panel (a2), while the corresponding *Z*-profile shows a band-pass filter (Figs. 8-c). For the parameter values in Fig. 11-b2 both the PSD-profile and *Z*-profile (not shown) display a band-pass filter, but the resonant peak in the *Z*-profile is more pronounced than in the PSD-profile and relatively bigger in comparison to the values of the same quantities at *f* = 0. This reflects the different ways in which the autonomous transient dynamics can be evoked by different types of input patterns, leading to significantly different results. This is not unexpected from the sets of parameter values in this graphs (panels a2 and b2) since the autonomous cell has resonance (in response to oscillatory inputs, *f_res_* > 0) but not intrinsic damped oscillations (*f_nat_* = 0). For the other panels in Fig. 11 the *Z*-profile is a relatively good predictor of the PSD-profiles. These correspond to low-pass filters (panels a1 and b1) and strong band-pass filters (panels a3 and b3; *f_res_* > 0 and *f_nat_* > 0).

**Figure 11:**
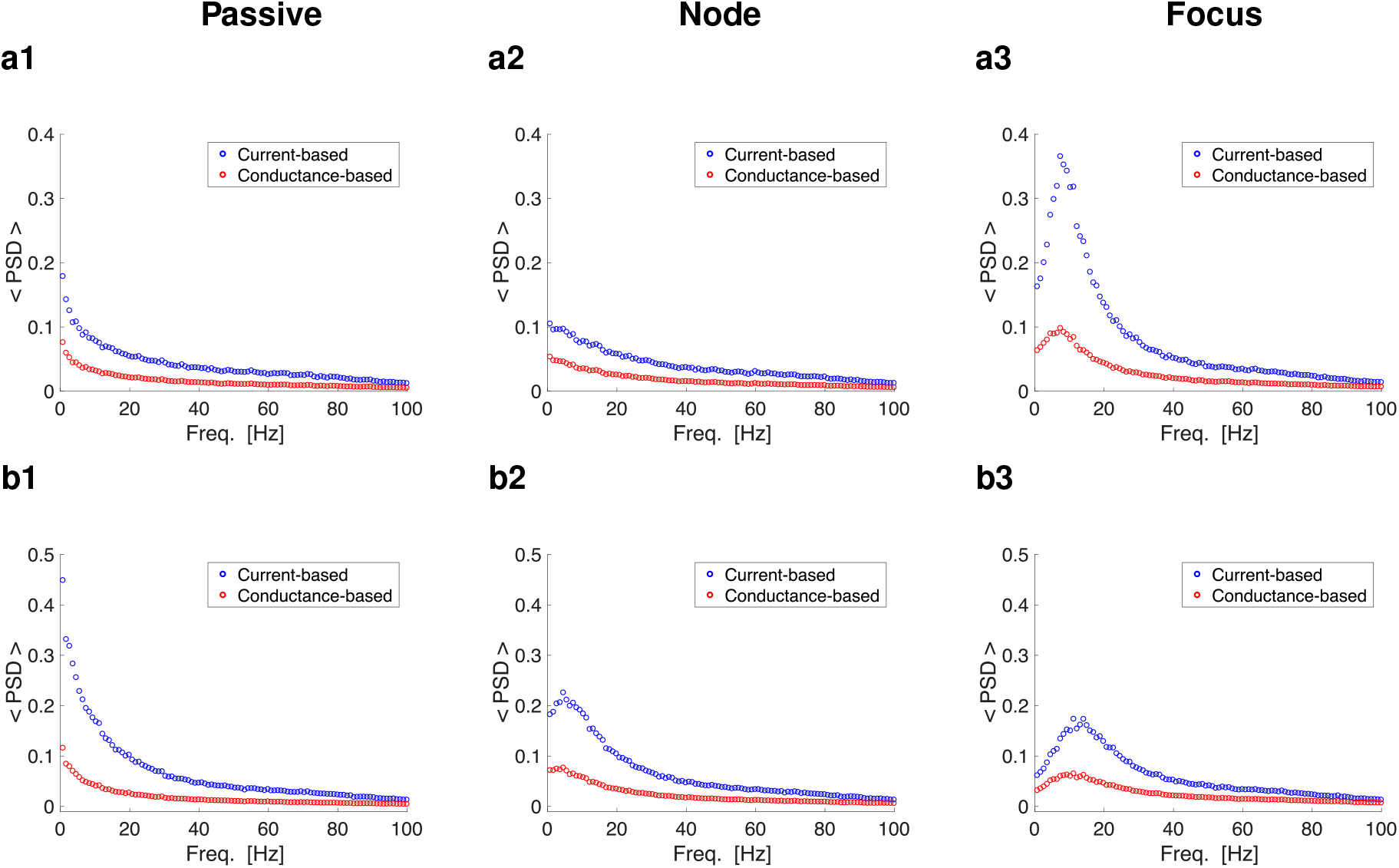
Average PSD for the *V* response to synaptic-like chirp-like inputs with arbitrarily distributed cycles for current- and conductance-based models. For current-based synaptic-like inputs we used eqs. (1)–(2). For conductance-based synaptic-like inputs we used the linear component of eqs. (3)–(4). The parameter values are as in Fig. 10. Each trial (*N_trials_* = 100) consists of a permutation of the cycle orders using as reference the ordered input patterns in Figs. 8-c to S15-c. The blue and red curves represent the < *PSD* > for the responses to synaptic-like current- and conductance-based inputs, respectively. **Column 1.** Passive cells. **Column 2.** Node (N-) cells. **Column 3.** Focus (F-) cells. **a1.** *g_L_* = 0.25 and *g*_1_ =0 (*f_nat_* = *f_res_* = 0). **a2.** *g_L_* = 0.25 and *g*_1_ = 0.25 (*f_nat_* = 0 and *f_res_* = 9). **a3.** *g_L_* = 0.05 and *g*_1_ = 0.3 (*f_nat_* = 8.1 and *f_res_* = 8). **b1.** *g_L_* = 0.1 and *g*_1_ = 0 (*f_nat_* = *f_res_* = 0). **b2.** *g_L_* = 0.1 and *g*_1_ = 0.2 (*f_nat_* = 0 and *f_res_* = 7). **b3.** *g_L_* = 0.1 and *g*_1_ = 0.8 (*f_nat_* = 12.3 and *f_res_* = 14). We used the additional parameter values: *C* = 1, *τ*_1_ = 100, *A_in_* = 1, *G_syn_* = 1 *E_syn_* = 1.

#### 3.8.2 The oscillatory voltage responses are stronger for current-than for conductance-based synaptic-like inputs

This is readily seen by comparing the blue and red curves in Fig. 11. These results are inherited from the responses of linear systems to synaptic current- and conductance-based constant inputs discussed in Section 3.1.2, and they can be understood in terms of our phase-plane diagrams discussion (compare Figs. 3-a and -c). Importantly, while in Fig. 11-a2 both responses show a low-pass filter, in Fig. 11-b2, the response to conductance-based synaptic inputs is close to a low-pass filter, while the response to current-based synaptic inputs is a well developed band-pass filter. In both cases, the autonomous cells have a stable node, exhibiting resonance, but not intrinsic (damped) oscillations.

#### 3.8.3 Oscillations (and lack of thereof) in response to Poisson distributed synaptic-like inputs

Poisson distributed inputs have in principle a very different structure than the synaptic-like chirp inputs we discussed above. However, there is a natural transition between the two types of patterns. Roughly speaking, synaptic-like chirp patterns can be first extended to include a larger number of frequencies (not necessarily integer), and more than one cycle for each input frequency according to some distribution. Therefore one expects the results discussed above to extend to Poisson distributed inputs. This is supported by the fact that for high enough Poisson input rates *ν*, synaptic-like inputs approximate constant inputs (this is true for Poisson distributed pulses with amplitude *g* → 0 and rate *v* → ∞), and therefore one should expect the voltage response PSD to approach these for constant inputs reflecting overshoots (low-pass filters) and damped oscillations (band-pass filters). However, for cells exhibiting overshoots there is a conflict between the response pattern “dictated” by the *Z*-profile (band-pass filter) and the low-pass filter pattern in response to constant inputs. For these cases, we expect the response pattern to be highly sensitive to the interplay of the Poisson rate *v* and model parameters. For other rates we expect a departure from the overall behavior described above, but less pronounced.

Our results are presented in Fig. 12 (see also Figs. S17 to S20). The blue and red curves correspond to the *V* responses to synaptic-like current- and conductance-based inputs respectively. The solid curves are smoothed versions of the dotted ones (to which they are superimposed). The green dots/solid curves correspond to the *V* responses to white noise and are used as a reference for comparison. The blue and red dashed curves are rescaled versions of the corresponding dot/solid curves so that they match the values of the green curves at *f* = 1. In all cases, the responses to synaptic-like inputs are attenuated as compared to the responses to white noise. The level of attenuation increases with increasing values of the input Poisson rate (*ν*) as we discussed below. Low-pass filters (panels a1 and b1) and strong band-pass filters (panels a3 and b3, F-cell) remain so with some variations. The responses of N-cells (panels a2 and b2) vary according to the input rate *ν*. Consistent with our results discussed above, for the cells that have resonance but not intrinsic (damped) oscillations (panels a2 and b2) and *ν* = 1000 (Fig. S19), the response can be either a low-pass filter or a band-pass filter depending on the model parameters, in particular the level of the (amplifying) positive feedback effect that increases with decreasing values of *g_L_* (compare panels a2, for *g_L_* = 0.25, and b2, for *g_L_* = 0.1). This remains the case for *ν* = 500 (Fig. S18) for the conductance-based response, but not for the current-based response (a band-pass filter emerges in panel a2). For *v* = 100 and *v* =10 (Figs. S17 and S20), both the current- and conductance-based responses show a band-pass filter.

**Figure 12:**
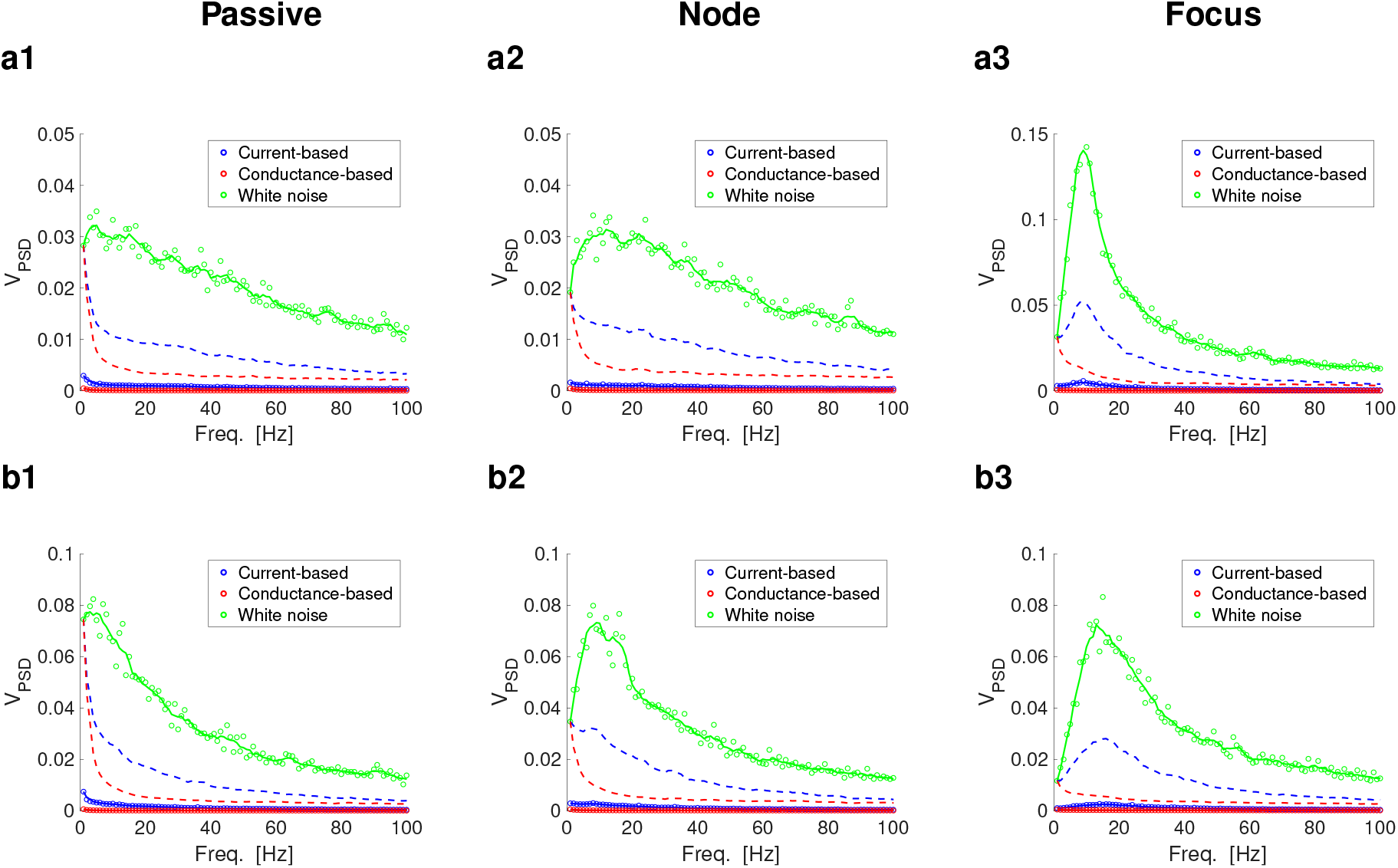
PSD for the *V* response to Poisson synaptic inputs trains (excitatory rate = 1000 Hz) for current- and conductance-based models. For current-based synaptic-like inputs we used eqs. (1)–(2). For conductance-based synaptic-like inputs we used the linear component of eqs. (3)–(4). The parameter values are as in Figs. 10 and 11. Poisson inputs (refractory time = 0.2 ms) were generated for a total duration of 1,000,000 ms. White noise had a variance 2*D* with *D* = 1. Blue dots and solid curves represent the PSD in response to current-based synaptic-like inputs. Red dots and solid curves represent the PSD in response to conductance-based synaptic-like inputs. Green dots and solid curves represent the PSD in response to white noise. The solid curves are a smoothed version (“moving”, 13 points) of the corresponding dots. The dashed curves are rescaled versions of the dots/solid curves to **Column 1.** Passive cells. **Column 2.** Node (N-) cells. **Column 3.** Focus (F-) cells. **a1.** *g_L_* = 0.25 and *g*_1_ =0 (*f_nat_* = *f_res_* = 0). **a2.** *g_L_* = 0.25 and *g*_1_ = 0.25 (*f_nat_* = 0 and *f_res_* = 9). **a3.** *g_L_* = 0.05 and *g*_1_ = 0.3 (*f_nat_* = 8.1 and *f_res_* = 8). **b1.** *g_L_* = 0.1 and *g*_1_ = 0 (*f_nat_* = *f_res_* = 0). **b2.** *g_L_* = 0.1 and *g*_1_ = 0.2 (*f_nat_* = 0 and *f_res_* = 7). **b3.** *g_L_* = 0.1 and *g*_1_ = 0.8 (*f_nat_* = 12.3 and *f_res_* = 14). We used the additional parameter values: *C* = 1, *τ*_1_ = 100, *A_in_* = 1, *G_syn,ex_* = 1, *E_syn,ex_* = 1, *τ*_Dec_ = 25.

The results described above persist when the input synaptic train consist of both excitatory and inhibitory synaptic-like inputs (e.g., Fig. S21).

The emergence of band-pass filters in response to synaptic-like inputs is strongly dependent on the synaptic input decay time *τ*_Dec_. For values of *τ*_Dec_ (= 25 ms) larger than in Figs. S17 to S21 and not realistic for fast synapses the band-pass filters are attenuated for F-cells (compare Figs. 12-a3 and -b3 with Figs. S19-a3 and -b3) and the N-cells show low-pass filters (compare Figs. 12-a2 and -b2 with Figs. S19-a2 and -b2).

#### 3.8.4 The oscillatory voltage responses are stronger for current-than for conductance-based synaptic-like inputs and biophysically plausible *in vivo* input rates

This is readily seen in Figs. 12 and S19 for the realistic values of *in vivo* input rates (*ν* = 1000) also used in controlled experiments [19]. For lower values of *ν* (Fig. S17 for *ν* = 100, Fig. S18 for *ν* = 500, Fig. S20 for *ν* = 10) and when synaptic inhibition is incorporated (Figs. S21 for excitatory *v* = 1000 and inhibitory *ν* = 500) the relative magnitudes of the current- and conductance-based responses depends on the input frequency regardless of whether the response has a low- or a bandpass filter. For even lower values of *ν* (Figs. S17 for *ν* = 10), the conductance-based response is stronger than the current-based response).

Together these results and the results o the previous Sections shed some light on the implications of the experimental findings in [19] where the intrinsically generated subthreshold oscillations observed in medial entorhinal cortex layer II stellate cells (SCs) have been shown to be strongly attenuated by current-based synaptic-like inputs and absent (or almost absent) in response to conductance-based synaptic-like inputs. Our findings suggest that STOs in SCs are generated by noise-dependent mechanisms in the presence of subthreshold resonance with at most strongly damped intrinsic oscillations [58], but not sustained limit cycle oscillations [129] in the presence of noise variability.

## 4 Discussion

Subthreshold (membrane potential) oscillations (STOs) have been observed in many neuron types in a variety of brain areas and have been argued to be functionally important for the generation of brain rhythms, sensory processing, encoding of information, communication of information via timing mechanisms and cross-frequency coupling (see more details and references in the Introduction).

Intrinsically generated STOs in single neurons require the presence of relatively slow restorative currents providing a negative feedback effect (currents having a resonant gating variable) and are amplified by fast regenerative currents providing a positive feedback effect (currents having an amplifying gating variable) (see Section 3.1 for more details). From a dynamical systems perspective, sustained STOs can be generated by limit cycle mechanisms or be noise-driven. In the latter case, the noiseless system may exhibit either damped oscillations (F-cells; the equilibrium has complex eigenvalues) or even overshoots (N-cells; the equilibrium has real eigenvalues) in response to abrupt changes in constant inputs. The interaction between Gaussian white noise and these autonomous transient dynamics may create sustained STOs [58, 109].

Neurons are subject to fluctuating inputs from a large number of synaptic currents generated by action potentials whose collective dynamics can be modeled as a high-rate Poisson process. Due to its high-rate, this synaptic noise has been approximated by Gaussian white noise or Ornstein-Uhlenbeck processes [130] (low-pass filtered versions of Gaussian white noise) [117–122]. Recent experimental results [19] on medial entorhinal cortex SCs, a prototypical intrinsic STO neuron [3, 5] and resonator [68], using artificially generated current- and conductance-based synaptic inputs driven by high-rate presynaptic Poisson spike trains, showed that STOs are still present in response to current-based synaptic inputs, but absent or strongly attenuated in response to conductance-based synaptic inputs. This would suggest that in realistic conditions the STO properties of SCs are not communicated to the network regime via synaptic mechanisms. On the other hand, in SCs and other cell types exhibiting STOs, the frequency of the STOs has been found to be correlated with the frequency of the networks in which they are embedded [1, 2, 26–28, 34, 41–44], suggesting intrinsic STOs in individual neurons may play, at least, an indirect role in the generation of network oscillations.

These issues are part of the more general question of how the response of neurons to periodic inputs (and to external inputs in general) depends on the interplay of the neuronal intrinsic properties and the properties of the input. Typical experiments on subthreshold and suprathreshold resonance use sinusoidal inputs, which change gradually with time. These studies are motivated by the fact that the resulting patterns can be used for the reconstruction of the system’s response to arbitrary time-dependent inputs under certain assumptions on both the input and the system (e.g., quasi-linearity). However, systems are not necessarily close to linearity and neuronal communication occurs via relatively fast synapses (e.g., AMPA and GABA_*A*_), which change more abruptly. These abrupt input changes evoke the autonomous intrinsic dynamics (damped oscillations or overshoots), which are occluded in response to gradual input changes. As a result, periodic (and also nonperiodic) sequences of sinusoidal and synaptic inputs are expected to produce different patterns and therefore the impedance profile will not be a good predictor of the voltage response to trains of synaptic inputs under general assumptions.

We set out to clarify these issues in a broader context. To develop the main set of ideas, we used a relatively simple neuronal model, the linearization of conductance-based models subject to additive current-based inputs and multiplicative conductance-based synaptic inputs. We then tested these ideas using a conductance-based model. We used three representative waveforms over a range of frequencies: sinusoidal, synaptic-like and square-wave (duty cycle equal to 0.5). Sinusoidal inputs are typically used to uncover the preferred oscillatory responses to external inputs as discussed above. Synaptic-like inputs represent the realistic ways in which communication between neurons occurs. Square-wave inputs can be considered as an intermediate between the first two. Sinusoidal and square-wave inputs share the waveform skeleton (they have the same frequency content except for the high frequency associated with the abrupt changes between phases), but sinusoidal inputs change gradually. Square-wave and synaptic inputs involve abrupt changes between minima and maxima, but the active part of the synaptic-like waveforms is independent of the period for a relatively large range of input frequencies. In addition, we used chirp-like (sinusoidal) inputs with discretely changing frequencies in order to be able to incorporate multiple frequencies in the same signal and we extended these chirps to include square- and synaptic-like waveforms.

We developed the notion of the peak/trough voltage envelope profiles 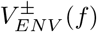 and the peak-to-trough impedance profiles *Z_ENV_*(*f*) as metrics to investigate the frequency-dependent voltage responses to periodic inputs in addition to the (standard) impedance amplitude profiles *Z*(*f*) and the corresponding voltage PSD (computed using Fourier transforms of the whole signal). Because the upper (peak) envelope is the most important quantity regarding the communication of information to the suprathreshold regime, we often refer to it indistinctly as *V_ENV_* or 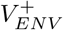. The differences between *V_ENV_* (or 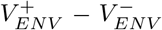) and the voltage PSD are due to the signal structure (*V_ENV_* captures only the envelope of the voltage response). The differences between *Z_ENV_* and *Z* capture the effect of the input signal (*Z_ENV_* is normalized by the input signal’s amplitude *A_in_*, while *Z* is normalized by the amplitude of its PSD). The differences between the *V_ENV_* and *Z_ENV_* profiles are due to the asymmetries in the voltage responses.

We showed that cells that exhibit resonance in response to sinusoidal inputs (*Z*-resonant cells) also show resonance in the *V_ENV_*- and *Z_ENV_*-responses to sinusoidal chirp inputs independently of whether they were N-cells or F-cells. This was expected given the gradual increase of the sinusoidal waveforms, but it served as a baseline for comparison with the other input types. For suprathreshold input amplitudes within some range, the frequency properties of *Z*-resonant cells in response to sinusoidal inputs are communicated to the suprathreshold regime in the form of evoked spiking resonance [127] or firing-rate resonance [63]. In contrast, *Z*-resonant N-cells are *V_ENV_* and *Z_ENV_* low-pass filters and *Z*-resonant F-cells have mild *V_ENV_* and *Z_ENV_* resonant properties. In other words, the cells’ subthreshold frequency-dependent properties, are not necessarily communicated to the spiking regime in response to non-sinusoidal inputs. The 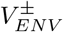 patterns in response to these two types of inputs are dominated by the autonomous intrinsic dynamics and this is particularly strong for the lower frequencies (longer periods) where the overshoots and damped oscillations can be prominent. In response to sinusoidal inputs, the autonomous transient dynamics develops gradually and their contribution to the 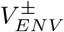 patterns remains occluded.

We used these protocols to compare the response of these neuron types to current- vs. conductance-based synaptic-like inputs. In all cases, the response to conductance-based inputs was attenuated as compared to the response to current-based inputs. Each one of the metrics produced different results, capturing different aspects of the voltage responses to these two types of inputs and their relationship with the input signals. For N-cells the responses to both current- and conductance-based inputs are *V_ENV_* low-pass filters. In contrast, for F-cells the *V_ENV_* responses to current-based inputs were band-pass filters, while the responses to conductance-based inputs were low-pass filters. These band-pass filters were generated as the result of the interplay of the autonomous transient dynamics (damped oscillations) and summation.

The *Z_ENV_* profiles tell a different story. For N-cells, the *Z_ENV_* responses to current-based inputs are low-pass filters, while they are band-pass filters for conductance-based inputs. For F-cells, the *Z_ENV_* responses to both current- and conductance-based inputs are band-pass filters. In both cases, the *Z_ENV_* band-pass filters in response to conductance-based inputs reflect troughs in the 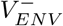 profiles rather than a real preferred voltage response. The *Z* profiles tell yet a different story. In all cases considered (N- and F-cells, current- and conductance-based inputs), the *Z* profiles are band-pass filters. For passive cells, where the autonomous transient dynamics are relatively simple (monotonic increase or decrease), the response patterns are dominated by summation and, while *Z_ENV_* and *Z* are low-pass filters, *V_ENV_* are high-pass filters.

In order to understand the contribution of the autonomous intrinsic dynamics to the generation of variability in the neuronal response patterns to external inputs, we used the three types of chirp-like inputs with arbitrarily ordered cycles. These inputs are an intermediate step between the regularly ordered and the fully irregular chirp-like inputs. The prototypical example of the latter (and the one we had in mind) are the synaptic-like inputs generated in response to spike-trains with Poisson-distributed spike times. The inputs have the same cycles for all trials, and hence the same frequency content, but each trial corresponded to a different permutation of the order of the cycles. The only source of uncertainty was the subset of all possible permutations of the cycle period. The differences in the voltage responses for cycles with the same period across trials were due to the differences in the initial conditions across trials for the same period. More specifically, for a given period (*T_k_*) the previous cycle has different periods across trials and therefore different voltage values at the end of these periods, which become the initial conditions for period *T_k_*. The variability of these initial conditions across trials involves not only the voltage but the (hidden) recovery variables. Our results demonstrated the emergence of variability of the voltage responses across trials for all input waveforms inherited from this mechanism. This variability was stronger for the F-cells than for the N-cells considered and, again, it did not require stochastic input fluctuations, but it was the result of the multiple different ways in which the inputs evoked the autonomous intrinsic dynamics.

The average voltage PSD (< *PSD* >) responses for F-cells were band-pass filters for both current- and conductance-based synaptic-like inputs with arbitrarily order periods. For N-cells, in contrast, the < *PSD* > responses to current-based input were low-pass filters, while the < *PSD* > responses to conductance-based inputs were low-pass filters or mild band-pass filters. This is consistent with the results in [19] and previous results showing that STOs in SCs are noise-driven [58, 113] (but see [129]). Our protocols consisted of the response of one cell type to variable inputs. More research is needed to understand the effects of variability across cells using some baseline attribute to all of them (e.g., same resonant properties or the same noise-driven oscillation properties).

Armed with these results, we compared the voltage responses (*V_PSD_*) of these cell types to high-rate Poisson distributed current- and conductance-based synaptic inputs and additive Gaussian white noise (noise-driven oscillations). The *V_PSD_*-profiles in response to both current- and conductance-based synaptic inputs were attenuated with respect to the response to white noise. The *V_PSD_*-profiles in response to current-based synaptic inputs were low-pass filters for F-cells and low-pass filters (or mild band-pass filters) for N-cells. The *V_PSD_*-profiles in response to conductance-based synaptic inputs were low-pass filters for all cell types. This is, again, consistent with the results in [19] and suggests that in contrast to the noise-driven oscillations that emerge in both F- and N-cells, the current-based synaptic-like Poisson-driven oscillation requires a stronger intrinsic oscillatory structure. These results also show that the responses to synaptic-like high-rate Poisson-driven inputs are not necessarily captured by the response to additive Gaussian white noise in contrast to standard assumptions. More research is needed to establish the conditions under which oscillations emerge in response to synaptic-like inputs. Representative examples show that oscillatory responses for current- and conductance-based synaptic-like inputs emerge for both F-and N-cells for lower Poisson rates. More research is also needed to establish the conditions under which the Gaussian white noise approximation provides good approximations to high-rate synaptic inputs.

The question arises whether the lack of oscillatory responses to synaptic-like inputs (almost complete for conductance-based and partial for current-based) implies the lack of communication of the intrinsic (noise-driven) oscillatory and resonant properties to the suprathreshold regime. While this requires a detailed analysis and is beyond the scope of this paper, we conducted a number of simulations to explore a few representative cases. We used the biophysical (conductance-based) *I_h_* + *I_Nap_* model (6)-(7) having two-dimensional subthreshold dynamics (Figs. S22 to S25) and compared them with the results using an integrate-and-fire model (Fig. S26) for which the subthreshold dynamics is one-dimensional. The results are mixed, but an important common theme is that the responses show output firing rate resonance (the response firing remains within a relatively small bounded range) even when the subthreshold resonance is not present. The most salient cases are shown in Fig. S22-a and -b. This phenomenon is absent in the absence of the intrinsic oscillatory dynamics for the leaky integrate-and-fire model (Fig. S22-d, Fig. S26). These results strongly depend on the input amplitude (Fig. S23 to S26). These results suggest that the oscillatory properties of individual neurons may be occluded at the subthreshold level, but they are still communicated to the suprathreshold regime. One may potentially contrast these observations against the cancellation of frequency-dependent redundant sensory stimuli in electric fish [131–134]. In such experiments, chirps create responses that are significantly enhanced compared with slower beats. This type of experimentation would certainly benefit from future discussions that derive from our work given the nature of their oscillatory inputs.

Our study is focused on a specific type of resonance in response to deterministic periodic inputs (in the form of chirps) and leaves out the response of systems to stochastic inputs that may lead to stochastic and coherence resonance [135–151] (and references therein). These are related, but different phenomena, which may be affected by the use of non-sinusoidal inputs of the form we use here. More research is required to understand these issues.

The synaptic-like inputs we use in this paper have constant amplitude across cycles. Previous work showed that synaptic short-term plasticity (STP, depression and facilitation) can lead to the emergence of temporal filters [152–154] and synaptic resonance [153, 155]. Additionally, STP has been shown to play a role in modulated intrinsic subthreshold oscillations [156–158]. Future work should consider the effects of adding STP to the protocols we used here. However, this may require the use of full periodic signals instead of chirps to capture phenomena mentioned above. A step in this direction has been done in [154].

Simply put, under rather general circumstances, neurons that exhibit subthreshold resonance in response to sinusoidal inputs would exhibit spiking resonance in response to the same inputs (for small enough values of the input amplitude), but would not exhibit spiking resonance for other types of inputs, in particular for synaptic-like waveforms, which are the ones that operate in networks. This includes the difference between the responses to current- and conductance-based inputs. These issues have been overlooked in the literature and, often, claims are made about the implications of resonance in response to sinusoidal inputs for spiking activity in realistic setups. Our results generate a number of predictions that can be tested experimentally *in vitro* using the dynamic clamp technique [159, 160] or *in vivo* using optogenetic tools [123, 161–163].

## Acknowledgments

This work was partially supported by the National Science Foundation grant DMS-1608077 (HGR).

## Appendix

### A Intrinsic and resonant oscillatory properties of 2D linear systems

Consider

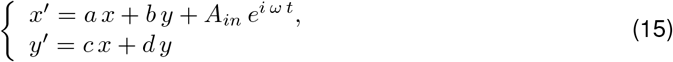

where *a, b, c* and *d* are constants, *ω* = 2*πf*/1000 > 0 is the input frequency and *A_in_* ≥ 0 is the input amplitude. The prime sign represents the derivative with respect to *t*. The units of *t* are ms and the units of *f* are Hz.

#### A.1 Intrinsic oscillations

The characteristic polynomial for the corresponding homogeneous system (*A_in_* = 0) is given by

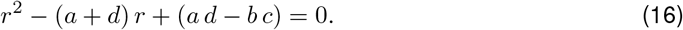

The eigenvalues are given by

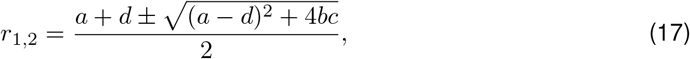

and the natural (intrinsic) frequency of the (damped) oscillations (in Hz if *t* has units of ms) is given by

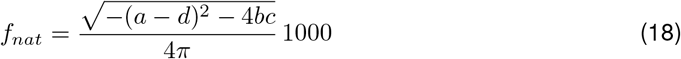

assuming (*a* – *d*)^2^ + 4*bc* < 0.

#### A.2 Resonance and the impedance amplitude profile

The impedance amplitude profile *Z*(*ω*) for system (15)-(16) is the magnitude

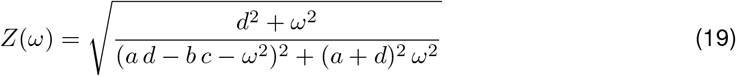

of the complex valued coefficient of the particular solution to the system

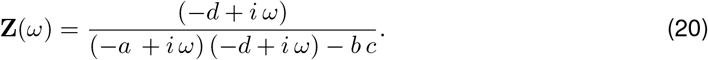

For 1D system, these quantities are given, respectively, by

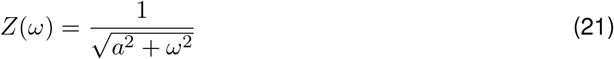

and

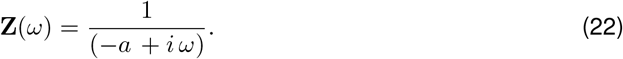

The resonance frequency *f_res_* (in Hz if *t* has units of ms) is the frequency at which *Z* reaches its maximum

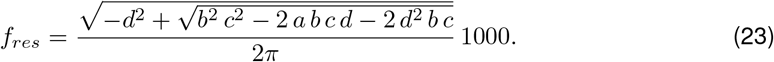

## Supplementary Material

**Figure S1:**
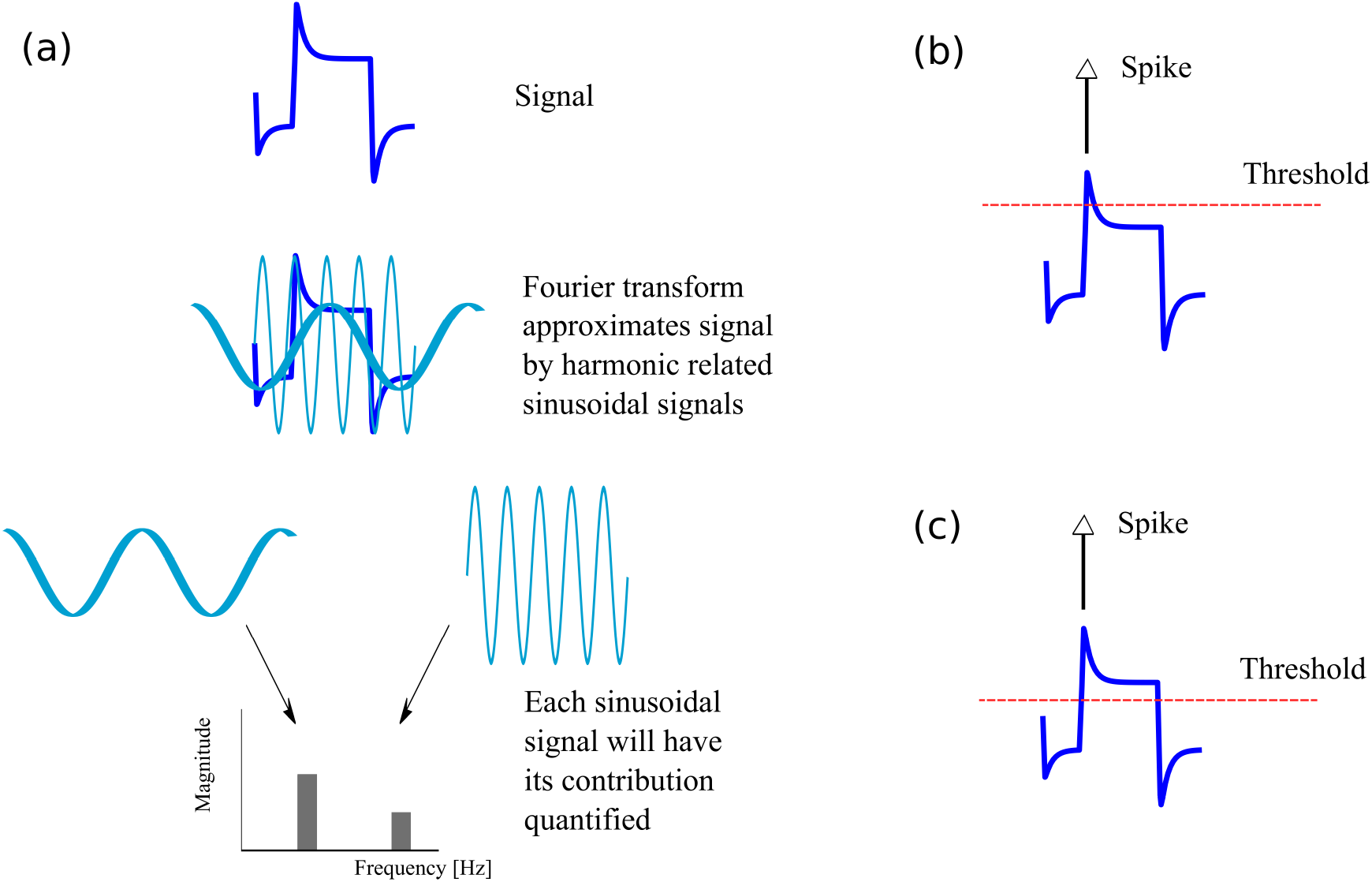
A schematic explanation for differences found between impedance computed from Fourier transforms (*Z*(*f*)) or from peaks and troughs (*Z*_ENV_). (a) The schematic voltage response signal is approximated in a Fourier Transform by the superposition of sinusoidal waves. These waves have their contribution quantified in the signal’s spectrum magnitude. Transients may contribute less than other parts of the signal. (b) Spikes appear when the voltage reaches the threshold in the model. Transients may reach the threshold easier independently of their contribution to the magnitude of the Fourier domain. (c) Even if the threshold is lower, the emitted spike will still be dependent on the transients as they may come earlier.

**Figure S2:**
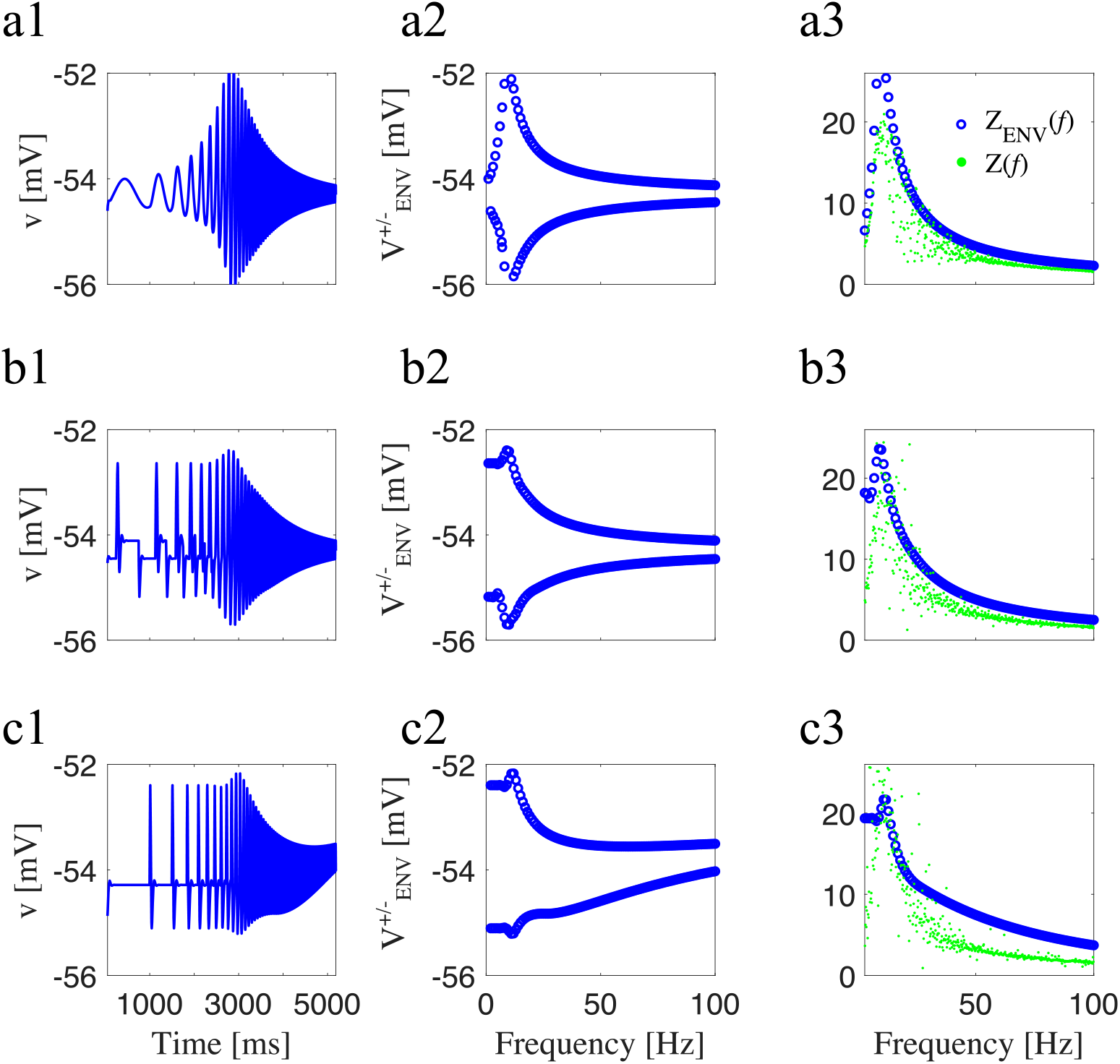
Neuronal response upon application of three different inputs (*I*_Nap_+*I*_h_ conductance-based F-cell model). Same parameters as in Fig S22(a) and Fig. S23. (a) Sinusoidal chirp. (b) Square-wave chirp. (c) Synaptic-like chirp. (a1,b1, and c1) voltage traces. (a2, b2, and c2) Voltage-response envelopes in the frequency-domain. (a3, b3, and c3) *Z*(*f*) (frequency-content) and *Z_ENV_*(*f*) (envelope) impedance. Parameter values are: *v*_*p*,1/2_ = −38 mV, *v_p,slp_* = 6.5 mV, *v*_*r*,1/2_ = −79.2 mV, *v_r,slp_* = 9.78 mV, *C* = 1 *μ*F/cm^2^, *E*_L_, = −65 mV, *E*_Na_ = 55 mV, *E*_h_ = −20 mV, *g*_L_ = 0.5 mS/cm^2^, *g_p_* = 0.5 mS/cm^2^, *g*_h_ = 1.5 mS/cm^2^, *I*_app_ = −2.5 *μ*A/cm^2^, *τ_r_* = 80 ms, *A*_in_ = 0.1 in (a), *A*_in_ = 0.07 in (b), and *A*_in_ = 0.5 in (c).

**Figure S3:**
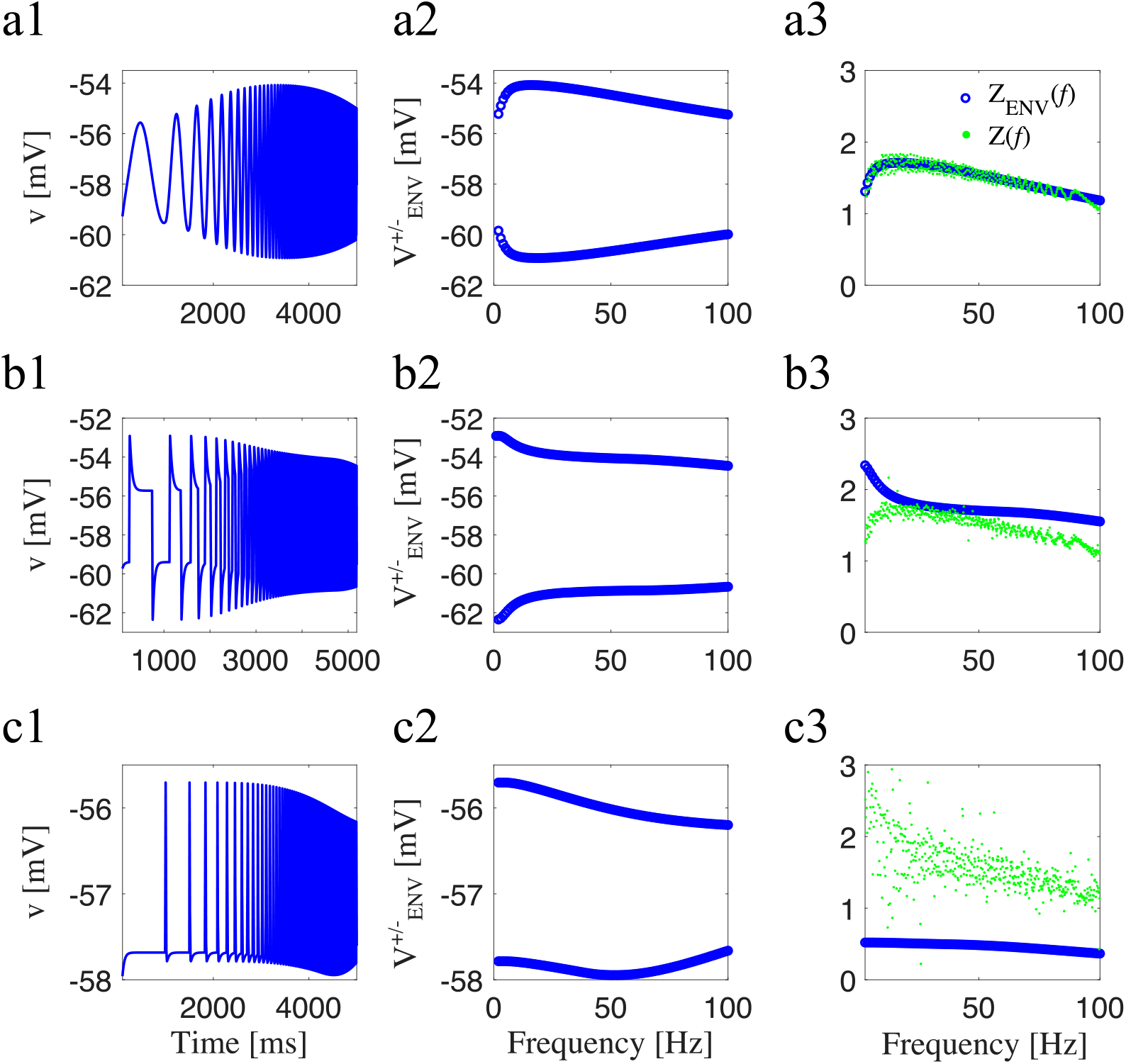
Neuronal response upon application of three different inputs (*I*_Nap_ + *I*_h_ conductance-based N-cell model). Same parameters as in Fig S22(b) and Fig. S4. (a) Sinusoidal chirp. (b) Square-wave chirp. (c) Synaptic-like chirp. (a1,b1, and c1) voltage traces. (a2, b2, and c2) Voltage-response envelopes in the frequency-domain. (a3, b3, and c3) *Z*(*f*) (frequency-content) and *Z_ENV_*(*f*) (envelope) impedance. Parameter values are: *v*_*p*,1/2_ = −38 mV, *v_p,slp_* = 6.5 mV, *v*_*r*,1/2_ = −79.2 mV, *v_r,slp_* = 9.78 mV, *C* = 1 *μ*F/cm^2^, *E*_L_ = −65 mV, *E*_Na_ = 55 mV, *E*_h_ = −20 mV, *g*_L_ = 0.5 mS/cm^2^, *g_p_* = 0.1 mS/cm^2^, *g*_h_ = 1.5 mS/cm^2^, *I*_app_ = −2.5 *μ*A/cm^2^, *τ_r_* = 80 ms, and *A*_in_ = 2.

**Figure S4:**
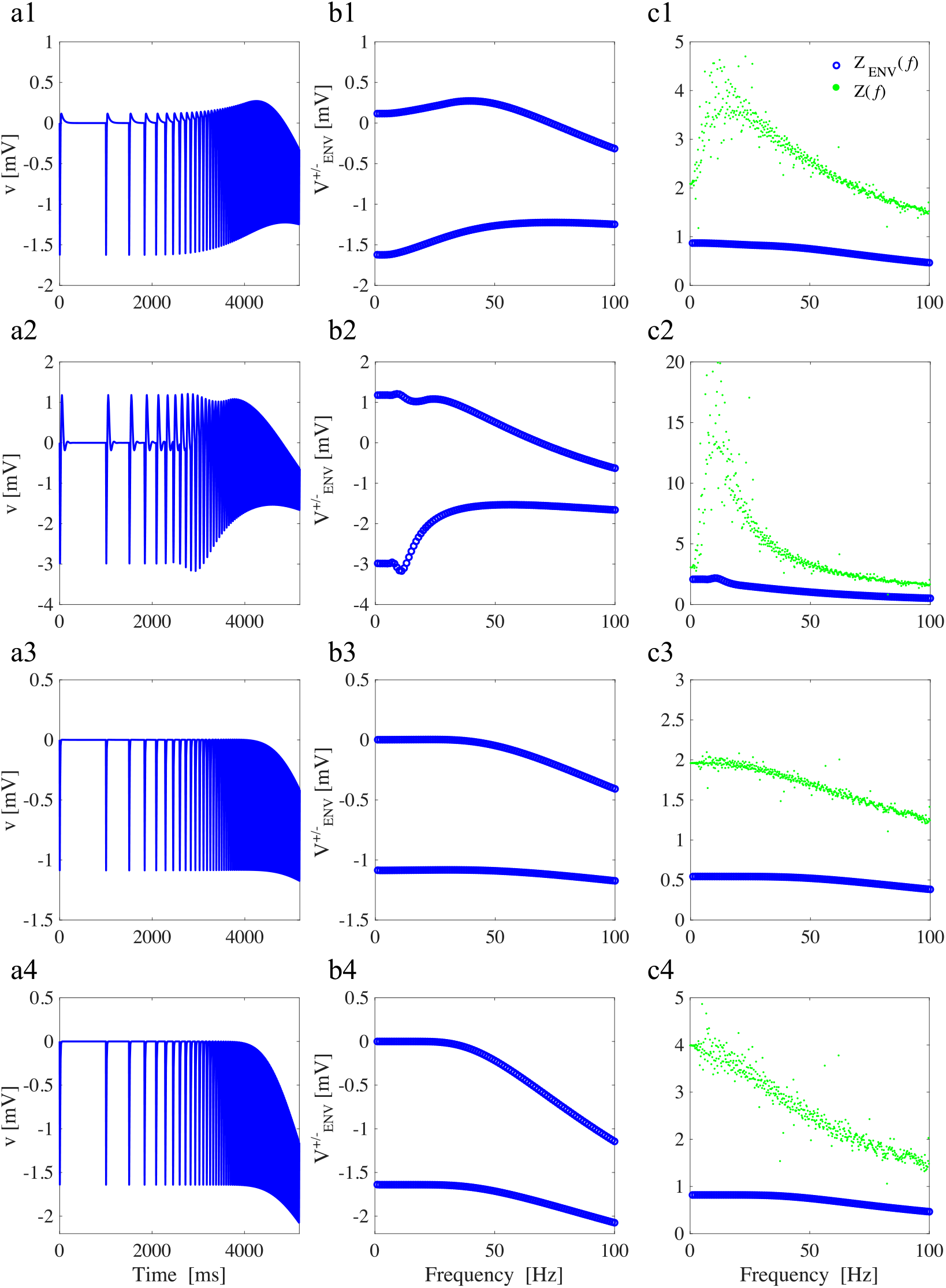
Neuronal response upon application of inhibitory synaptic-like inputs (linear model). (a) Voltage traces. (b) Voltage-response envelopes in the frequency-domain. (c) *Z*(*f*) (frequency-content) and *Z_ENV_*(*f*) (envelope) impedance. (a1, b1, and c1) Overshoot, parameter values: *C* = 1, *g*_L_ = 0.25, *g*_1_ = 0.25, *τ*_1_ = 100 ms, and *A*_in_ = 1 (as in Fig. 4). (a2, b2, and c2) Subthreshold oscillations, parameter values: *C* = 1, *g*_L_ = 0.05, *g*_1_ = 0.3, *τ*_1_ = 100 ms, and *A*_in_ = 1 (as in Fig. 5). (a3, b3, c3) No overshoot or subthreshold oscillations, parameter values: *C* = 1, *g*_L_ = 0.5, *g*_1_ = 0.0105, *τ*_1_ = 100 ms, and *A*_in_ = 1 (as in Fig. S10). (a4, b4, c4) Parameter values: *C* = 1, *g*_L_ = 0.25, *g*_1_ = 0, *τ*_1_ = 100 ms, and *A*_in_ = 1 (as in Fig. 9).

**Figure S5:**
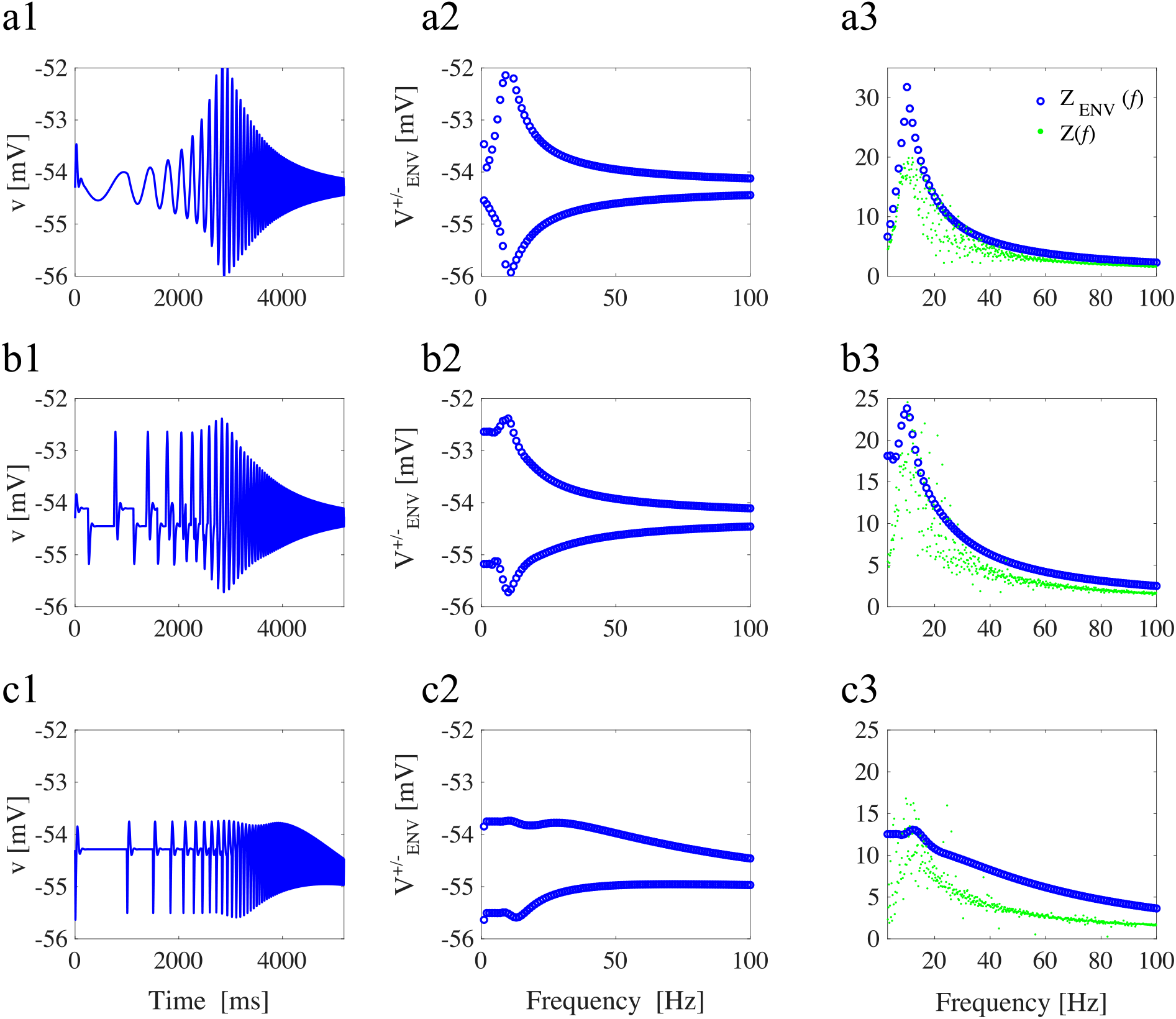
Neuronal response upon application three different inhibitory inputs (*I*_Nap_ + *I*_h_ conductance-based F-cell model). Same parameters as in Fig S22(a) and Fig. S23. (a) Sinusoidal chirp. (b) Square-wave chirp. (c) Synaptic-like chirp. (a1,b1, and c1) voltage traces. (a2, b2, and c2) Voltage-response envelopes in the frequency-domain. (a3, b3, and c3) *Z*(*f*) (frequency-content) and *Z_ENV_*(*f*) (envelope) impedance. Parameter values are: *v*_*p*,1/2_ = −38 mV, *v_p,slp_* = 6.5 mV, *v*_*r*,1/2_ = −79.2 mV, *v_r,slp_* = 9.78 mV, *C* = 1 *μ*F/cm^2^, *E*_L_, = −65 mV, *E*_Na_ = 55 mV, *E*_h_ = −20 mV, *g*_L_ = 0.5 mS/cm^2^, *g_p_* = 0.5 mS/cm^2^, *g*_h_ = 1.5 mS/cm^2^, *I*_app_ = −2.5 *μ*A/cm^2^, *τ_r_* = 80 ms, *A*_in_ = −0.1 in (a), *A*_in_ = −0.07 in (b), and *A*_in_ = −0.5 in (c).

**Figure S6:**
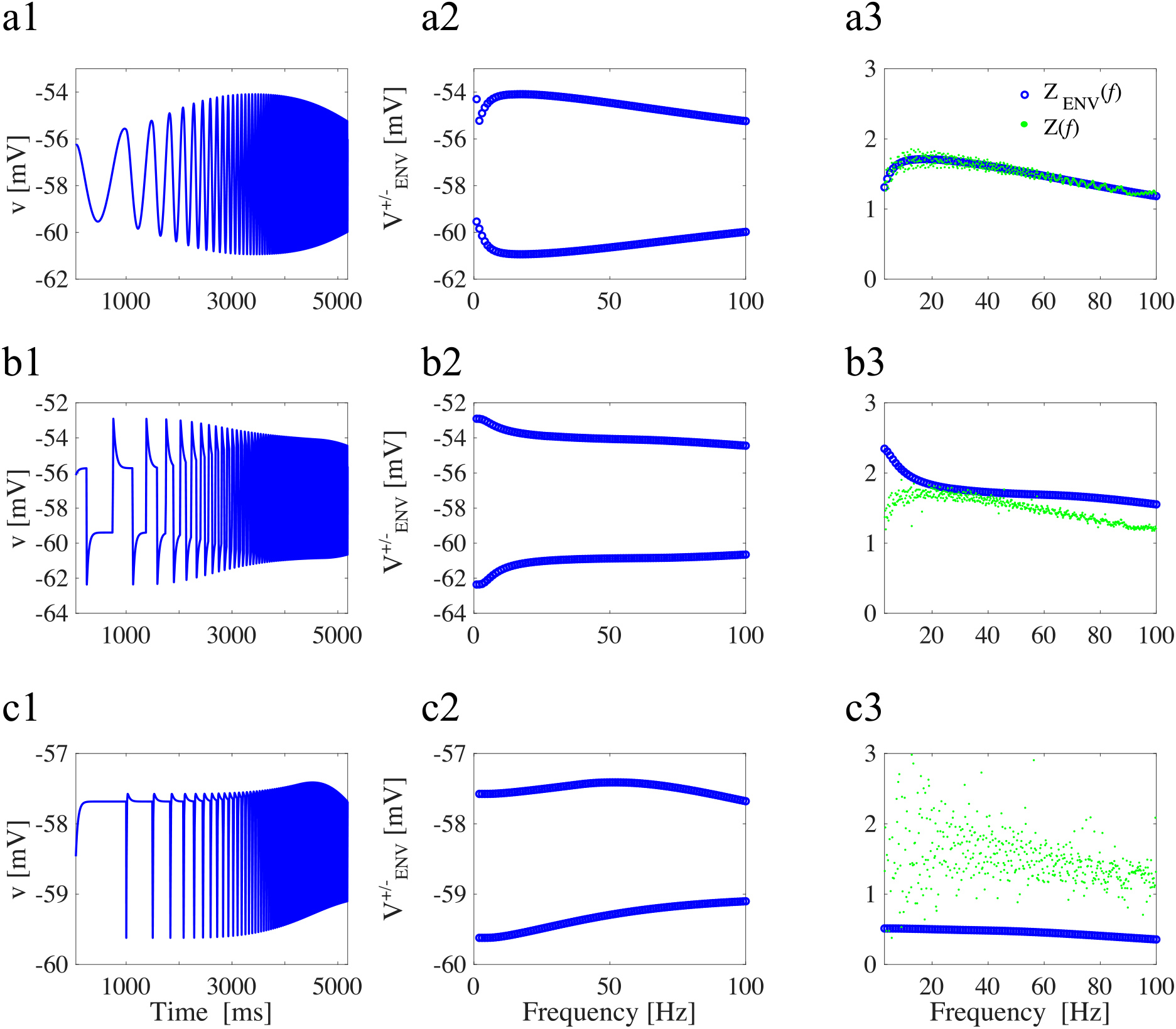
Neuronal response upon application three different inhibitory inputs (*I*_Nap_ + *I*_h_ conductance-based N-cell model). Same parameters as in Fig S22(b) and Fig. S4. (a) Sinusoidal chirp. (b) Square-wave chirp. (c) Synaptic-like chirp. (a1,b1, and c1) voltage traces. (a2, b2, and c2) Voltage-response envelopes in the frequency-domain. (a3, b3, and c3) *Z*(*f*) (frequency-content) and *Z_ENV_*(*f*) (envelope) impedance. Parameter values are: *v*_*p*,1/2_ = −38 mV, *v_p,slp_* = 6.5 mV, *v*_*r*,1/2_ = −79.2 mV, *v_r,slp_* = 9.78 mV, *C* = 1 *μ*F/cm^2^, *E*_L_, = −65 mV, *E*_Na_ = 55 mV, *E*_h_ = −20 mV, *g*_L_, = 0.5 mS/cm^2^, *g_p_* = 0.1 mS/cm^2^, *g*_h_ = 1.5 mS/cm^2^, *I*_app_ = −2.5 *μ*A/cm^2^, *τ_r_* = 80 ms, and *A*_in_ = −2.

**Figure S7:**
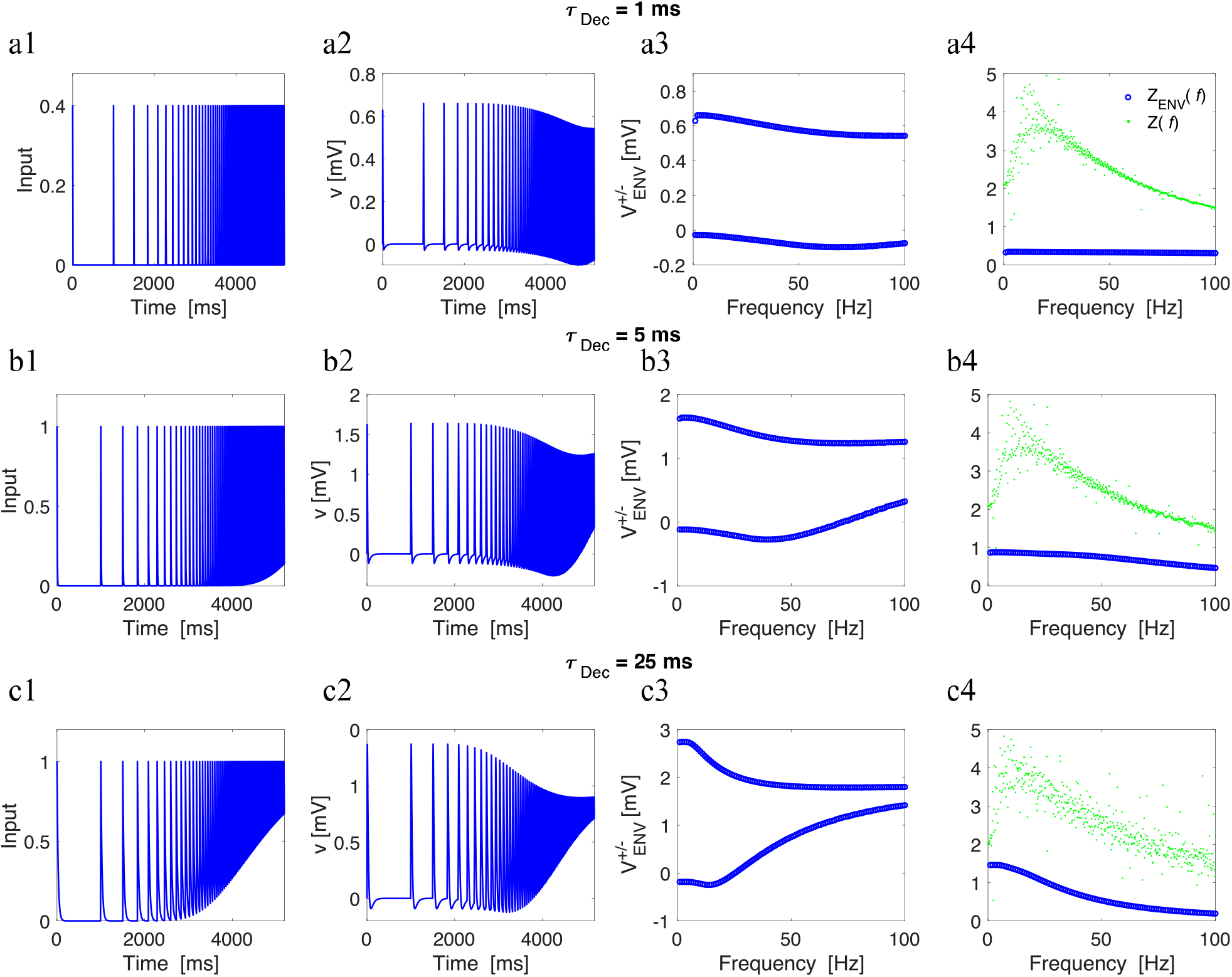
Neuronal responses to synaptic-like inputs are affected by the summation effect produced by the synaptic time decay (linear model, overshoot). (a) *τ*_Dec_ = 1 ms. (b) *τ*_Dec_ = 5 ms. (c). *τ*_Dec_ = 25 ms. (a1, b1, and c1) Time-dependent input. (a2, b2, and c2) Voltage traces. (a3, b3, and c3) Voltage-response envelopes. (a4, b4, and c4) *Z*(*f*) (frequency-content) and *Z_ENV_*(*f*) (envelope) impedance. We used the following parameter values: *C* =1, *g*_L_ = 0.25, *g*_1_ = 0.25, *τ*_1_ = 100 ms, and *A*_in_ = 1, same model as in Fig. 4.

**Figure S8:**
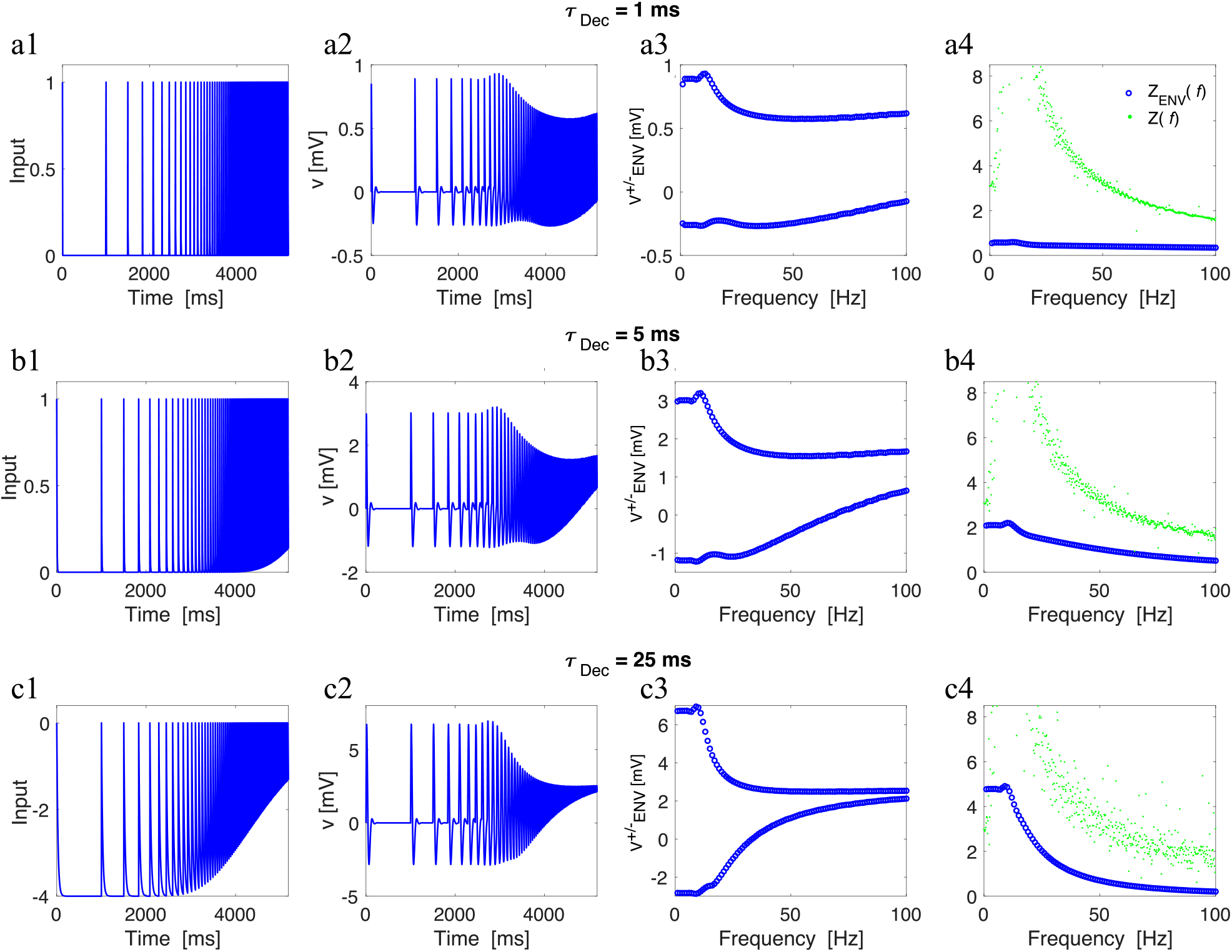
Neuronal responses to synaptic-like inputs are affected by the summation effect produced by the synaptic time decay (linear model, subthreshold oscillations). (a) *τ*_Dec_ = 1 ms. (b) *τ*_Dec_ = 5 ms. (c). *τ*_Dec_ = 25 ms. (a1, b1, and c1) Time-dependent input. (a2, b2, and c2) Voltage traces. (a3, b3, and c3) Voltage-response envelopes. (a4, b4, and c4) *Z*(*f*) (frequency-content) and *Z_ENV_*(*f*) (envelope) impedance. We used the following parameter values: *C* = 1, *g*_L_ = 0.05, *g*_1_ = 0.3, *τ*_1_ = 100 ms, and *A*_in_ = 1, same model as in Fig. 5.

**Figure S9:**
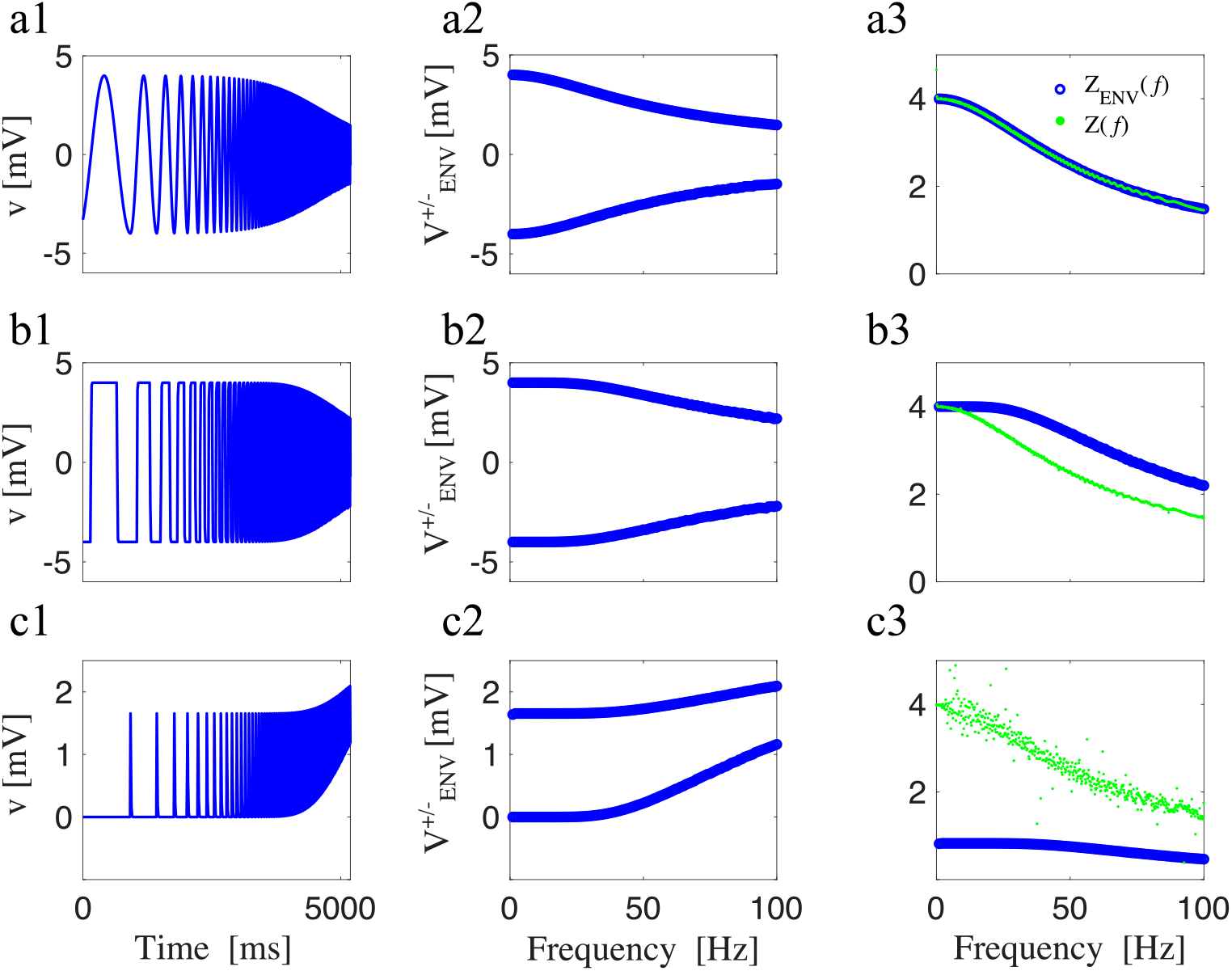
Neuronal response upon application of three different inputs (1D-linear model; *g*_1_ = 0). (a) Sinusoidal chirp. (b) Square-wave chirp. (c) Synaptic-like chirp. (a1,b1, and c1) voltage traces. (a2, b2, and c2) Voltage-response envelopes in the frequency-domain. (a3, b3, and c3) *Z*(*f*) (frequency-content) and *Z_ENV_*(*f*) (envelope) impedance. We used the following parameter values: *C* = 1, *g*_L_ = 0.25, *g*_1_ = 0, *τ*_1_ = 100 ms, and *A*_in_ = 1.

**Figure S10:**
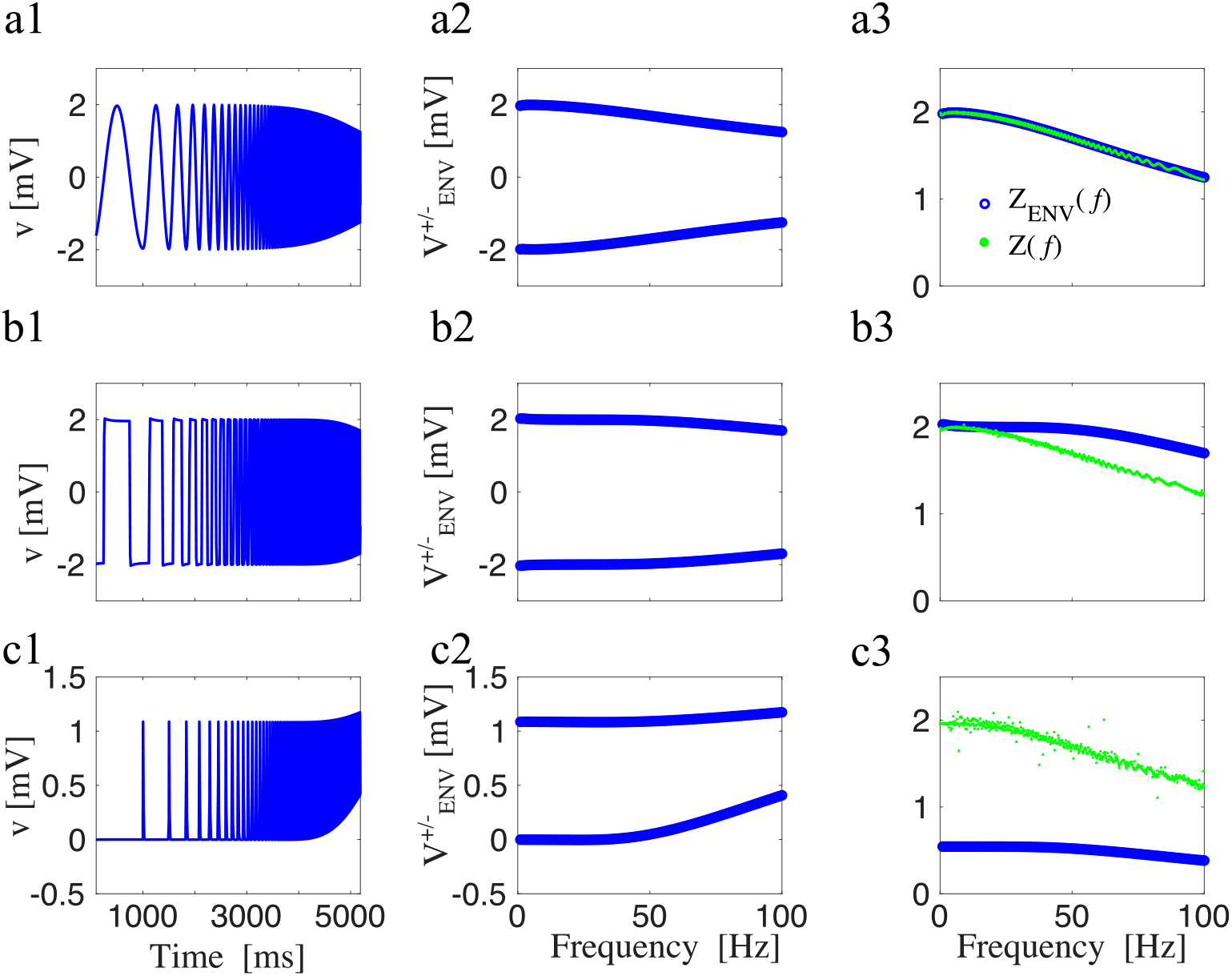
Neuronal response upon application of three different inputs (linear model, no overshoot or subthreshold oscillations). (a) Sinusoidal chirp. (b) Square-wave chirp. (c) Synaptic-like chirp. (a1,b1, and c1) voltage traces. (a2, b2, and c2) Voltage-response envelopes in the frequency-domain. (a3, b3, and c3) *Z*(*f*) (frequency-content) and *Z_ENV_*(*f*) (envelope) impedance. We used the following parameter values: *C* = 1, *g*_L_ = 0.5, *g*_1_ = 0.0105, *τ*_1_ = 100 ms, and *A*_in_ = 1.

**Figure S11:**
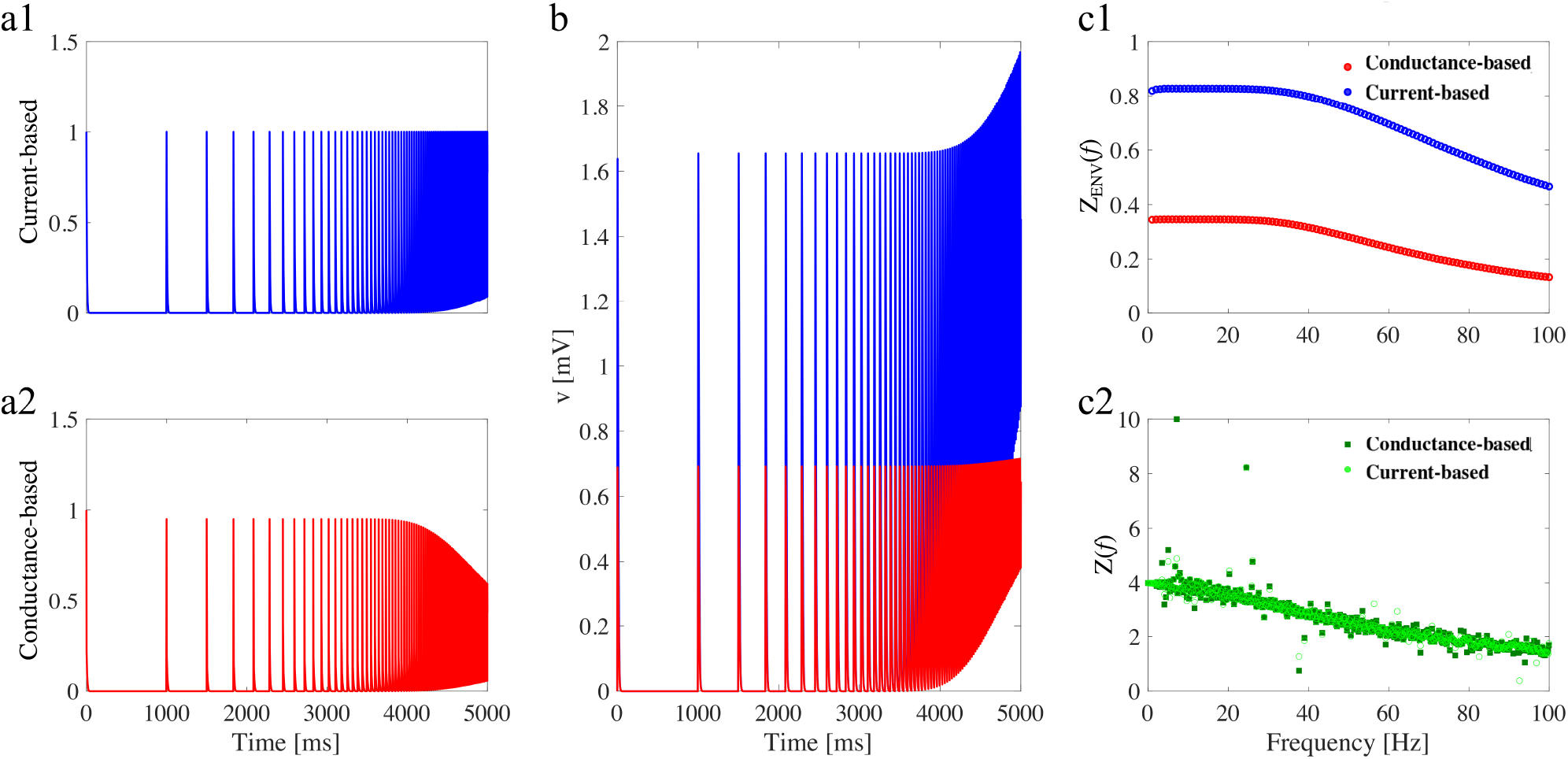
Comparison between conductance-based and current-based inputs (1D-linear model, *g*_1_ = 0). (a1) Current-based input. (a2) Conductance based input. (b) Voltage traces (same colors as in a1 and a2). (c1) *Z_ENV_*(*f*) (envelope) impedance. (c2) *Z*(*f*) (frequency-content). We used the following parameter values: *C* = 1, *g*_L_ = 0.25, *g*_1_ = 0, *τ*_1_ = 100 ms, and *A*_in_ = 1, same model as in Fig. S9.

**Figure S12:**
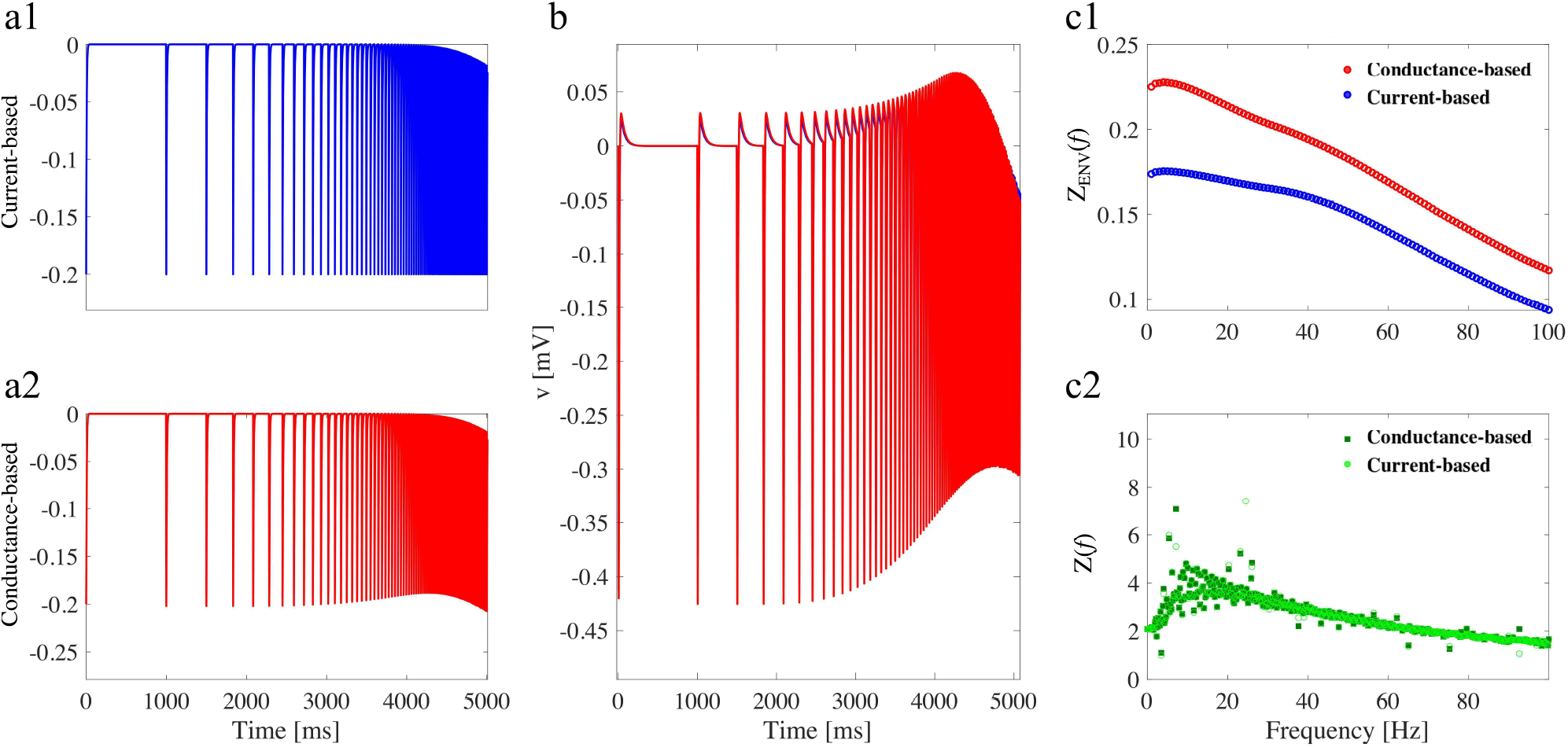
Comparison between conductance-based and current-based inputs (linear N-cell model and inhibitory responses). (a1) Current-based input. (a2) Conductance based input. (b1) Voltage traces (same colors as in a1 and a2). (c1) *Z_ENV_*(*f*) (envelope) impedance. (c2) *Z*(*f*) (frequency-content). We used the following parameter values: *C* = 1, *g*_L_ = 0.25, *g*_1_ = 0.25, *τ*_1_ = 100 ms, and *A*_in_ = −0.2, same model as in Fig. 4.

**Figure S13:**
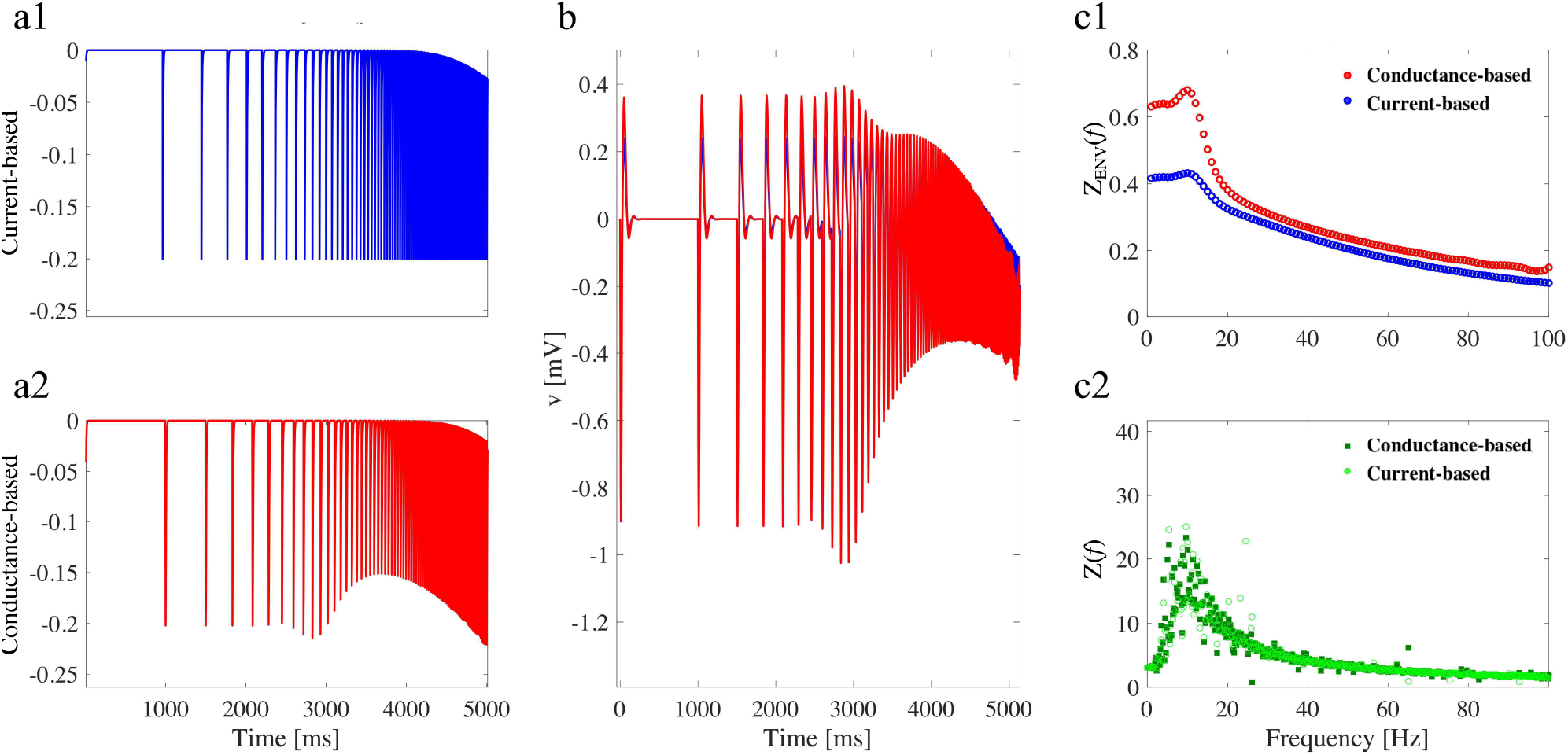
Comparison between conductance-based and current-based inputs (linear F-cell model and inhibitory responses). (a1) Current-based input. (a2) Conductance based input. (b1) Voltage traces (same colors as in a1 and a2). (c1) *Z_ENV_*(*f*) (envelope) impedance. (c2) *Z*(*f*) (frequency-content). We used the following parameter values: *C* = 1, *g*_L_ = 0.05, *g*_1_ = 0.3, *τ*_1_ = 100 ms, and *A*_in_ = −0.2, same model as in Fig. 5.

**Figure S14:**
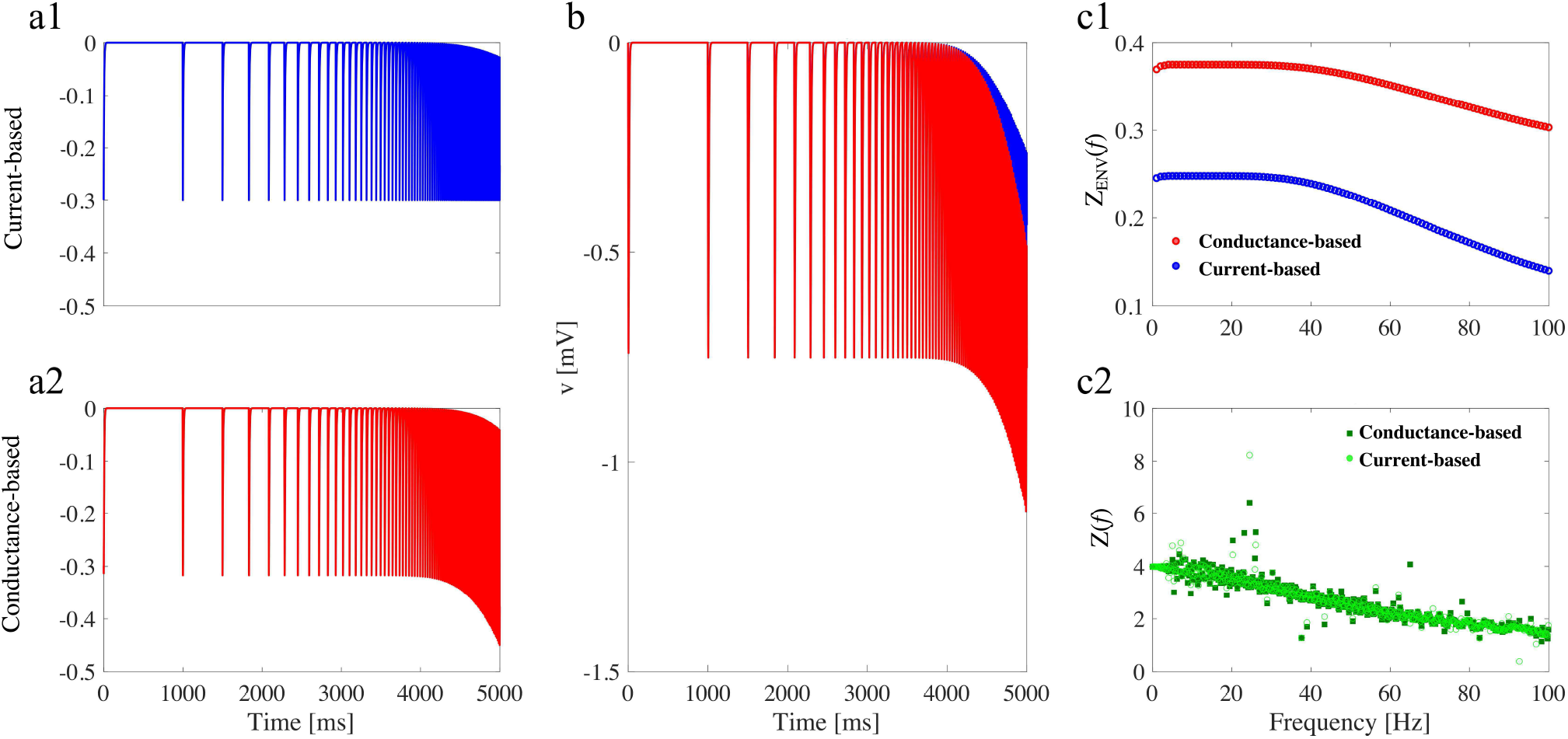
Comparison between conductance-based and current-based inputs (1D-linear model and inhibitory responses, *g*_1_ = 0). (a1) Current-based input. (a2) Conductance based input. (b1) Voltage traces (same colors as in a1 and a2). (c1) *Z_ENV_*(*f*) (envelope) impedance. (c2) *Z*(*f*) (frequency-content). We used the following parameter values: *C* =1, *g*_L_ = 0.25, *g*_1_ = 0, *τ*_1_ = 100 ms, and *A*_in_ = −0.3, same model as in Fig. 9.

**Figure S15:**
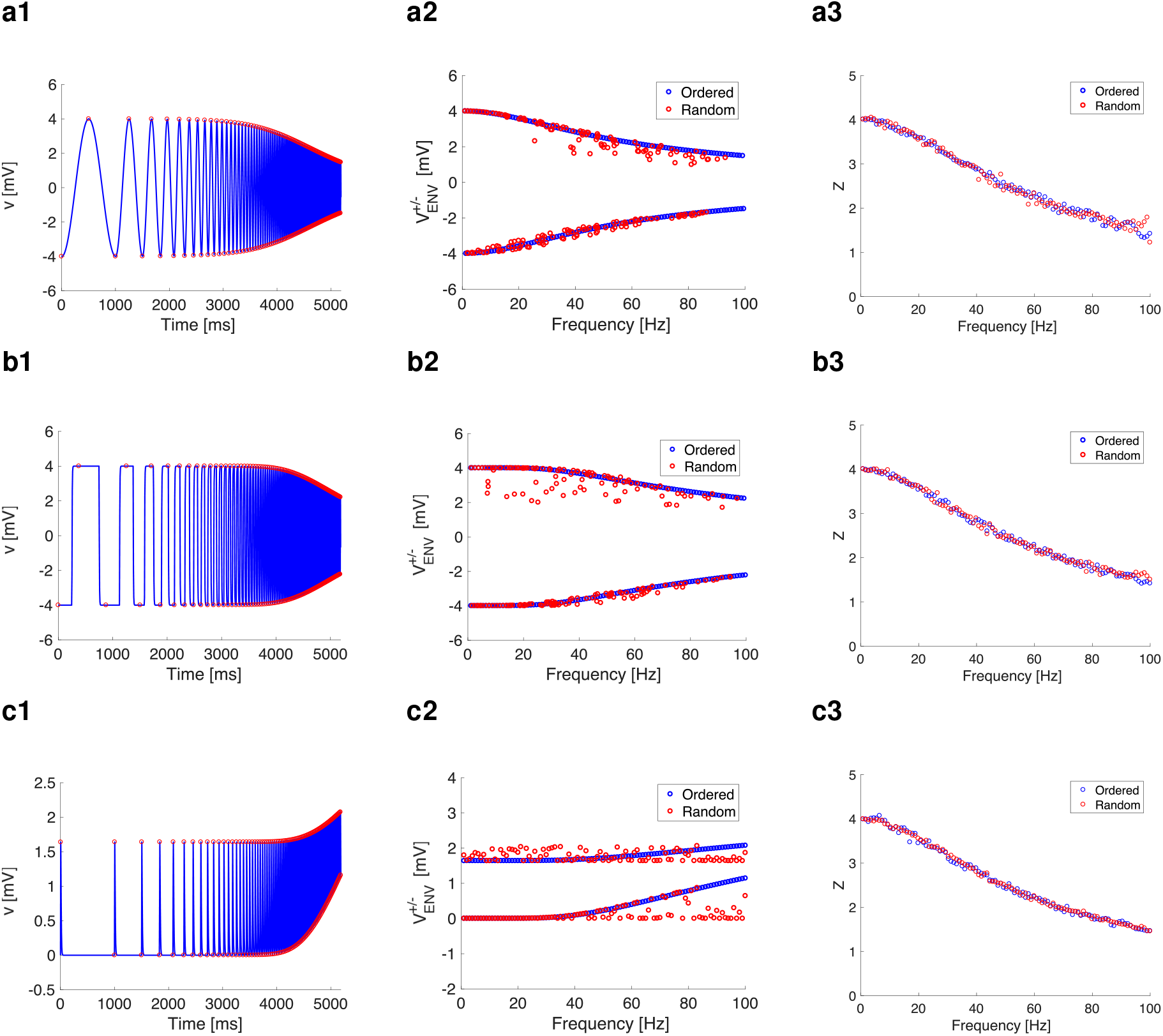
Comparison of neuronal response between ordered and random input (1D-linear model, *g*_1_ = 0). (a) Sinusoidal chirp. (b) Square-wave chirp. (c) Excitatory synaptic-like chirp. (a1, b1, and c1) voltage traces with peaks and troughs marked by red circles. (a2, b2, and c2) Voltage-response envelopes in the frequency domain (blue is ordered input; red is shuffled input as in Fig. 1b). (a3, b3, and c3) *Z*(*f*) (frequency-content) for ordered and shuffled inputs. In this model, we used the following parameter values: *C* = 1, *g*_L_ = 0.05, *g*_1_ = 0, *τ*_1_ = 100 ms, and *A*_in_ = 1.

**Figure S16:**
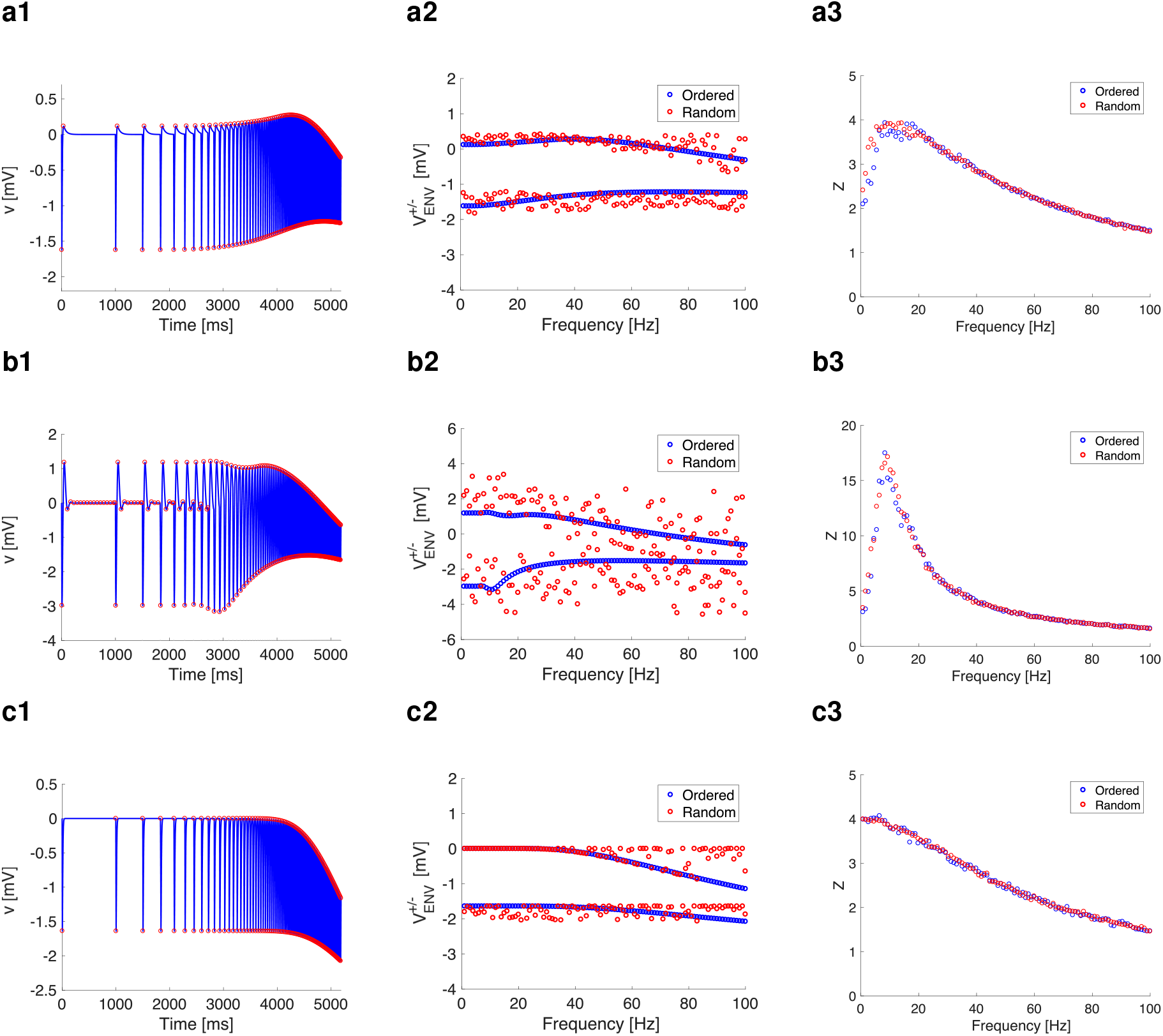
Comparison of neuronal response of the individual cells upon inhibitory inputs. Same models as in Figs. 8, 9 and S15. (a1, b1, and c1) voltage traces with peaks and troughs marked by red circles. (a2, b2, and c2) Voltage-response envelopes in the frequency domain (blue is ordered input; red is shuffled input as in Fig. 1b). (a3, b3, and c3) *Z*(*f*) (frequency-content) for ordered and shuffled inputs. *C* = 1. *A_in_* = 1, *G_syn_* = 1, *E_syn_* = −1. **a.** *g_L_* = 0.25, *g*_1_ = 0.25, *τ*_1_ = 100. **b.** *g_L_* = 0.05, *g*_1_ = 0.3, *τ*_1_ = 100. **c.** *g_L_* = 0.25, *g*_1_ = 0.

**Figure S17:**
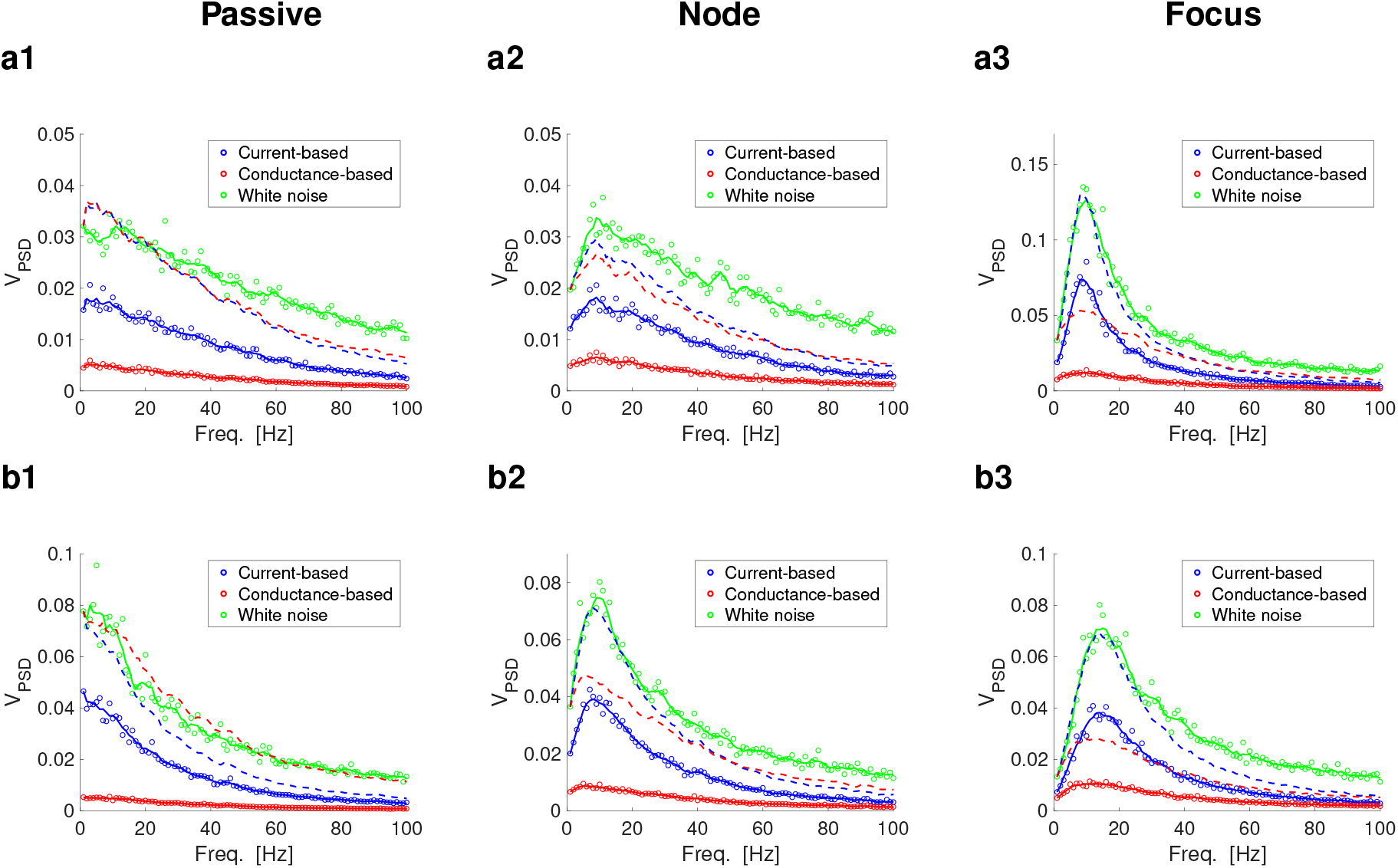
PSD for the *V* response to Poisson synaptic inputs trains (rate = 100 Hz) for current-and conductance-based models. For current-based synaptic-like inputs we used eqs. (1)–(2). For conductance-based synaptic-like inputs we used the linear component of eqs. (3)–(4). The parameter values are as in Figs. 10 and 11. Poisson inputs (refractory time = 0.2 ms) were generated for a total duration of 1,000,000 ms. White noise had a variance 2*D* with *D* = 1. Blue dots and solid curves represent the PSD in response to current-based synaptic-like inputs. Red dots and solid curves represent the PSD in response to conductance-based synaptic-like inputs. Green dots and solid curves represent the PSD in response to white noise. The solid curves are a smoothed version (“moving”, 13 points) of the corresponding dots. The dashed curves are rescaled versions of the dots/solid curves to **Column 1.** Passive cells.of **Column 2.** Node (N-) cells. **Column 3.** Focus (F-) cells. **a1.** *g_L_* = 0.25 and *g*_1_ =0 (*f_nat_* = *f_res_* = 0). **a2.** *g_L_* = 0.25 and *g*_1_ = 0.25 (*f_nat_* = 0 and *f_res_* = 9). **a3.** *g_L_* = 0.05 and *g*_1_ = 0.3 (*f_nat_* = 8.1 and *f_res_* = 8). **b1.** *g_L_* = 0.1 and *g*_1_ =0 (*f_nat_* = *f_res_* = 0). **b2.** *g_L_* = 0.1 and *g*_1_ = 0.2 (*f_nat_* = 0 and *f_res_* = 7). **b3.** *g_L_* = 0.1 and *g*_1_ = 0.8 (*f_nat_* = 12.3 and *f_res_* = 14). We used the additional parameter values: C = 1, *τ*_1_ = 100, *A_in_* = 1, *G_syn_* = 1, *E_syn_* = 1, *τ*_Dec_ = 5.

**Figure S18:**
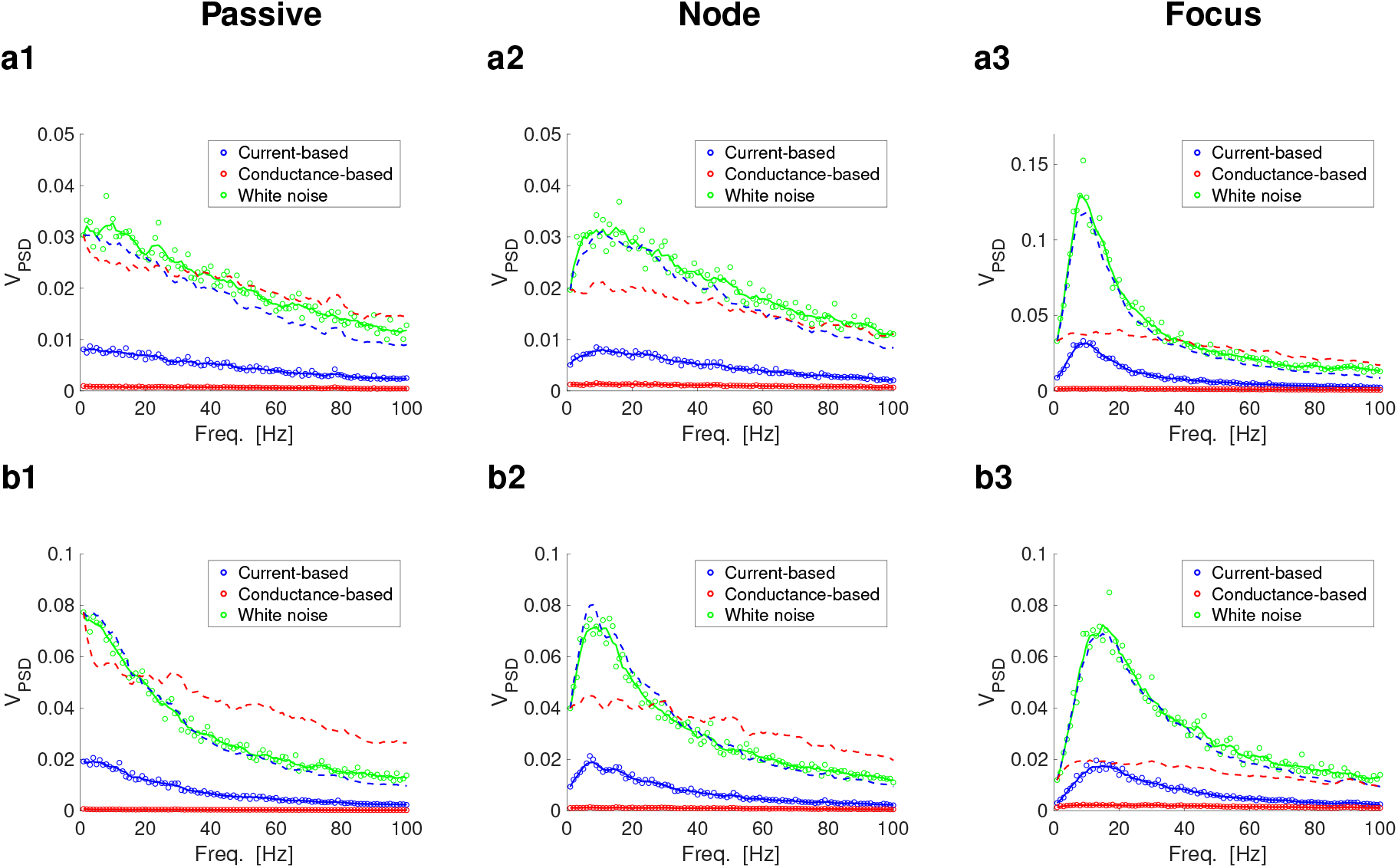
PSD for the *V* response to Poisson synaptic inputs trains (rate = 500 Hz) for current-and conductance-based models. For current-based synaptic-like inputs we used eqs. (1)–(2). For conductance-based synaptic-like inputs we used the linear component of eqs. (3)–(4). The parameter values are as in Figs. 10 and 11. Poisson inputs (refractory time = 0.2 ms) were generated for a total duration of 1,000,000 ms. White noise had a variance 2*D* with *D* = 1. Blue dots and solid curves represent the PSD in response to current-based synaptic-like inputs. Red dots and solid curves represent the PSD in response to conductance-based synaptic-like inputs. Green dots and solid curves represent the PSD in response to white noise. The solid curves are a smoothed version (“moving”, 13 points) of the corresponding dots. The dashed curves are rescaled versions of the dots/solid curves to **Column 1.** Passive cells.of **Column 2.** Node (N-) cells. **Column 3.** Focus (F-) cells. **a1.** *g_L_* = 0.25 and *g*_1_ =0 (*f_nat_* = *f_res_* = 0). **a2.** *g_L_* = 0.25 and *g*_1_ = 0.25 (*f_nat_* = 0 and *f_res_* = 9). **a3.** *g_L_* = 0.05 and *g*_1_ = 0.3 (*f_nat_* = 8.1 and *f_res_* = 8). **b1.** *g_L_* = 0.1 and *g*_1_ =0 (*f_nat_* = *f_res_* = 0). **b2.** *g_L_* = 0.1 and *g*_1_ = 0.2 (*f_nat_* = 0 and *f_res_* = 7). **b3.** *g_L_* = 0.1 and *g*_1_ = 0.8 (*f_nat_* = 12.3 and *f_res_* = 14). We used the additional parameter values: *C* = 1, *τ*_1_ = 100, *A_in_* = 1, *G_syn_* = 1, *E_syn_* = 1, *τ*_Dec_ = 5.

**Figure S19:**
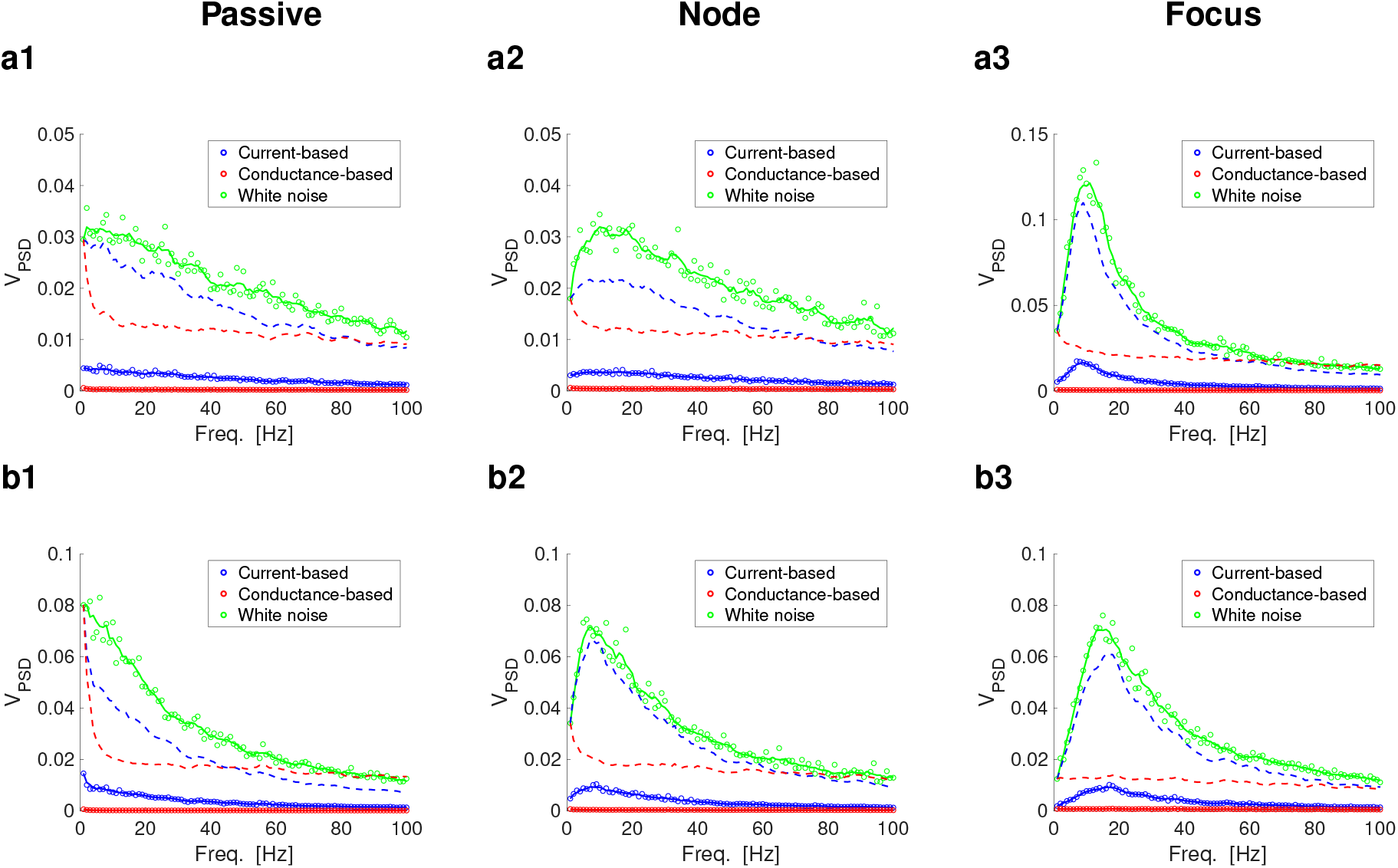
PSD for the *V* response to Poisson synaptic inputs trains (rate = 1000 Hz) for current- and conductance-based models. For current-based synaptic-like inputs we used eqs. (1)–(2). For conductance-based synaptic-like inputs we used the linear component of eqs. (3)–(4). The parameter values are as in Figs. 10 and 11. Poisson inputs (refractory time = 0.2 ms) were generated for a total duration of 1,000,000 ms. White noise had a variance 2*D* with *D* = 1. Blue dots and solid curves represent the PSD in response to current-based synaptic-like inputs. Red dots and solid curves represent the PSD in response to conductance-based synaptic-like inputs. Green dots and solid curves represent the PSD in response to white noise. The solid curves are a smoothed version (“moving”, 13 points) of the corresponding dots. The dashed curves are rescaled versions of the dots/solid curves to **Column 1.** Passive cells.of **Column 2.** Node (N-) cells. **Column 3.** Focus (F-) cells. **a1.** *g_L_* = 0.25 and *g*_1_ =0 (*f_nat_* = *f_res_* = 0). **a2.** *g_L_* = 0.25 and *g*_1_ = 0.25 (*f_nat_* = 0 and *f_res_* = 9). **a3.** *g_L_* = 0.05 and *g*_1_ = 0.3 (*f_nat_* = 8.1 and *f_res_* = 8). **b1.** *g_L_* = 0.1 and *g*_1_ = 0 (*f_nat_* = *f_res_* = 0). **b2.** *g_L_* = 0.1 and *g*_1_ = 0.2 (*f_nat_* = 0 and *f_res_* = 7). **b3.** *g_L_* = 0.1 and *g*_1_ = 0.8 (*f_nat_* = 12.3 and *f_res_* = 14). We used the additional parameter values: *C* = 1, *τ*_1_ = 100, *A_in_* = 1, *G_syn_* = 1, *E_syn_* = 1, *τ*_Dec_ = 5.

**Figure S20:**
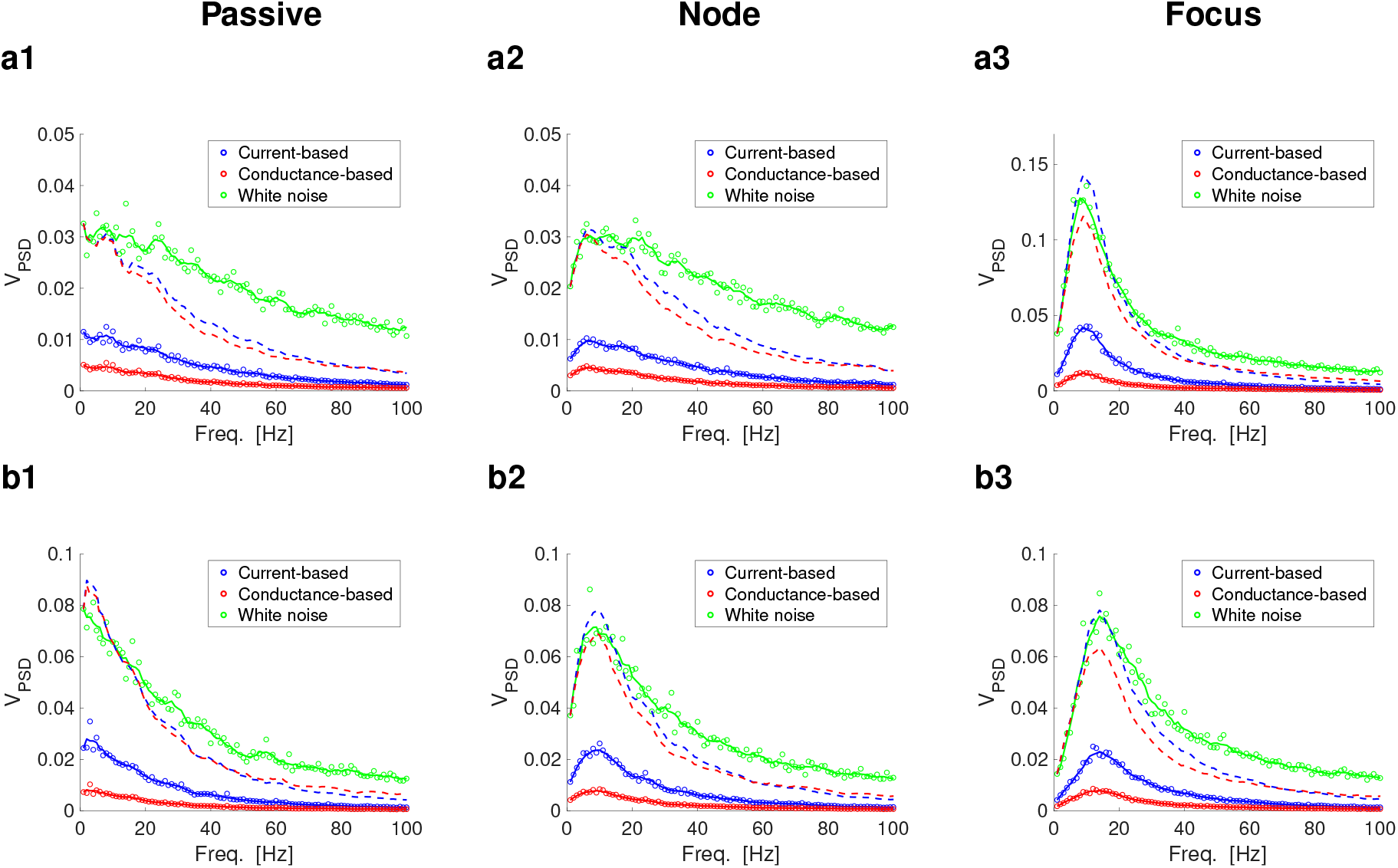
PSD for the *V* response to Poisson synaptic inputs trains (rate = 10 Hz) for current- and conductance-based models. For current-based synaptic-like inputs we used eqs. (1)–(2). For conductance-based synaptic-like inputs we used the linear component of eqs. (3)–(4). The parameter values are as in Figs. 10 and 11. Poisson inputs (refractory time = 0.2 ms) were generated for a total duration of 1,000,000 ms. White noise had a variance 2*D* with *D* = 1. Blue dots and solid curves represent the PSD in response to current-based synaptic-like inputs. Red dots and solid curves represent the PSD in response to conductance-based synaptic-like inputs. Green dots and solid curves represent the PSD in response to white noise. The solid curves are a smoothed version (“moving”, 13 points) of the corresponding dots. The dashed curves are rescaled versions of the dots/solid curves to **Column 1.** Passive cells.of **Column 2.** Node (N-) cells. **Column 3.** Focus (F-) cells. **a1.** *g_L_* = 0.25 and *g*_1_ =0 (*f_nat_* = *f_res_* = 0). **a2.** *g_L_* = 0.25 and *g*_1_ = 0.25 (*f_nat_* = 0 and *f_res_* = 9). **a3.** *g_L_* = 0.05 and *g*_1_ = 0.3 (*f_nat_* = 8.1 and *f_res_* = 8). **b1.** *g_L_* = 0.1 and *g*_1_ =0 (*f_nat_* = *f_res_* = 0). **b2.** *g_L_* = 0.1 and *g*_1_ = 0.2 (*f_nat_* = 0 and *f_res_* = 7). **b3.** *g_L_* = 0.1 and *g*_1_ = 0.8 (*f_nat_* = 12.3 and *f_res_* = 14). We used the additional parameter values: *C* = 1, *τ*_1_ = 100, *A_in_* = 1, *G_syn_* = 1, *E_syn_* = 1, *τ*_Dec_ = 5.

**Figure S21:**
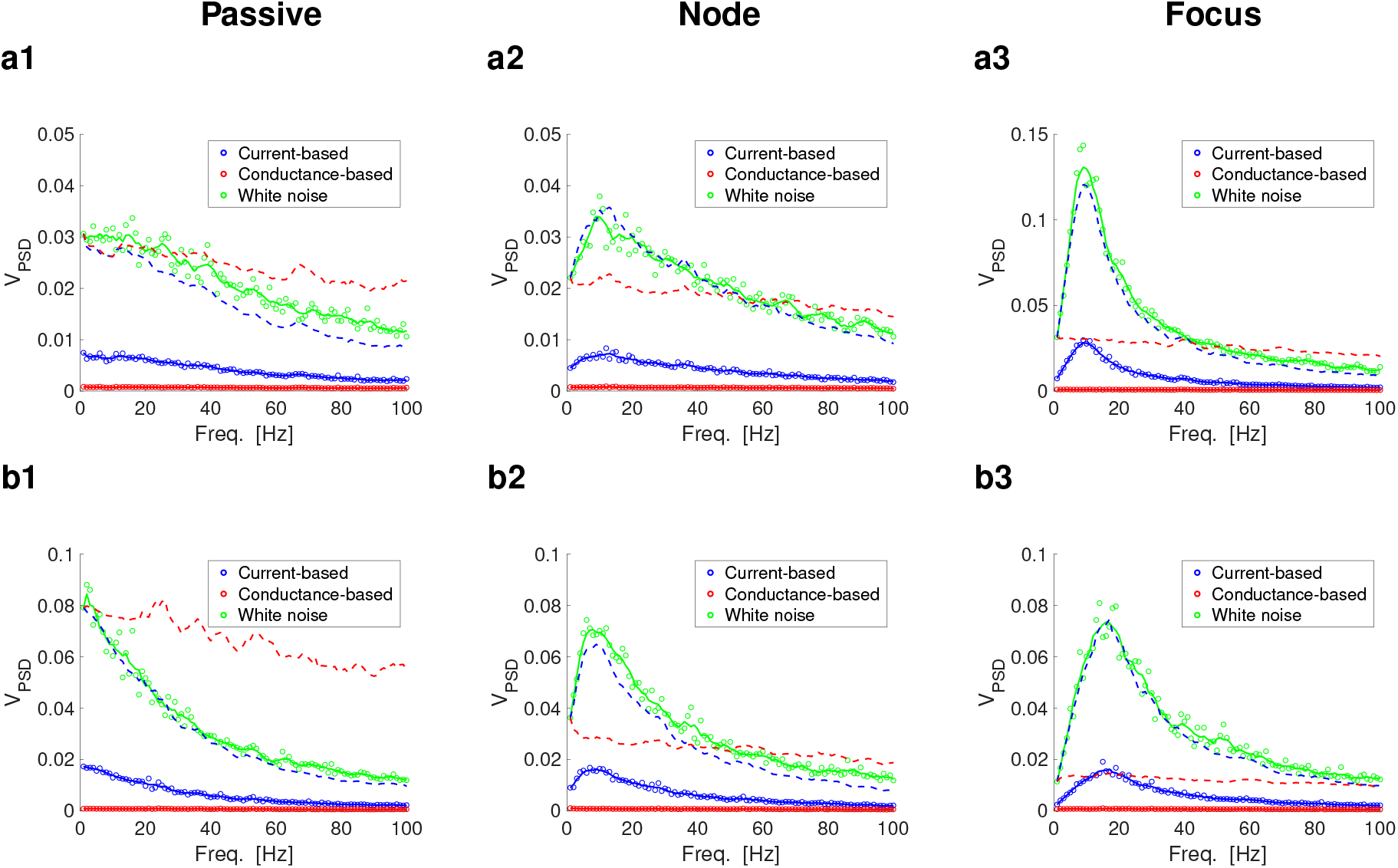
PSD for the *V* response to Poisson synaptic inputs trains (excitatory rate = 1000 Hz, inhibitory rate = 500 Hz) for current- and conductance-based models. For current-based synaptic-like inputs we used eqs. (1)–(2). For conductance-based synaptic-like inputs we used the linear component of eqs. (3)–(4). The parameter values are as in Figs. 10 and 11. Poisson inputs (refractory time = 0.2 ms) were generated for a total duration of 1,000,000 ms. White noise had a variance 2*D* with *D* = 1. Blue dots and solid curves represent the PSD in response to current-based synaptic-like inputs. Red dots and solid curves represent the PSD in response to conductance-based synaptic-like inputs. Green dots and solid curves represent the PSD in response to white noise. The solid curves are a smoothed version (“moving”, 13 points) of the corresponding dots. The dashed curves are rescaled versions of the dots/solid curves to **Column 1.** Passive cells.of **Column 2.** Node (N-) cells. **Column 3.** Focus (F-) cells. **a1.** *g_L_* = 0.25 and *g*_1_ = 0 (*f_nat_* = *f_res_* = 0). **a2.** *g_L_* = 0.25 and *g*_1_ = 0.25 (*f_nat_* = 0 and *f_res_* = 9). **a3.** *g_L_* = 0.05 and *g*_1_ = 0.3 (*f_nat_* = 8.1 and *f_res_* = 8). **b1.** *g_L_* = 0.1 and *g*_1_ =0 (*f_nat_* = *f_res_* = 0). **b2.** *g_L_* = 0.1 and *g*_1_ = 0.2 (*f_nat_* = 0 and *f_res_* = 7). **b3.** *g_L_* = 0.1 and *g*_1_ = 0.8 (*f_nat_* = 12.3 and *f_res_* = 14). We used the additional parameter values: *C* =1, *τ*_1_ = 100, *A_in_* = 1, *G_syn,ex_* = 1, *G_syn,in_* = 1 (*G_syn,in_* = −1 for current-clamp), *E_syn,ex_* = 1 *E_syn,in_* = −0.5, *τ*_Dec_ = 5.

**Figure S22:**
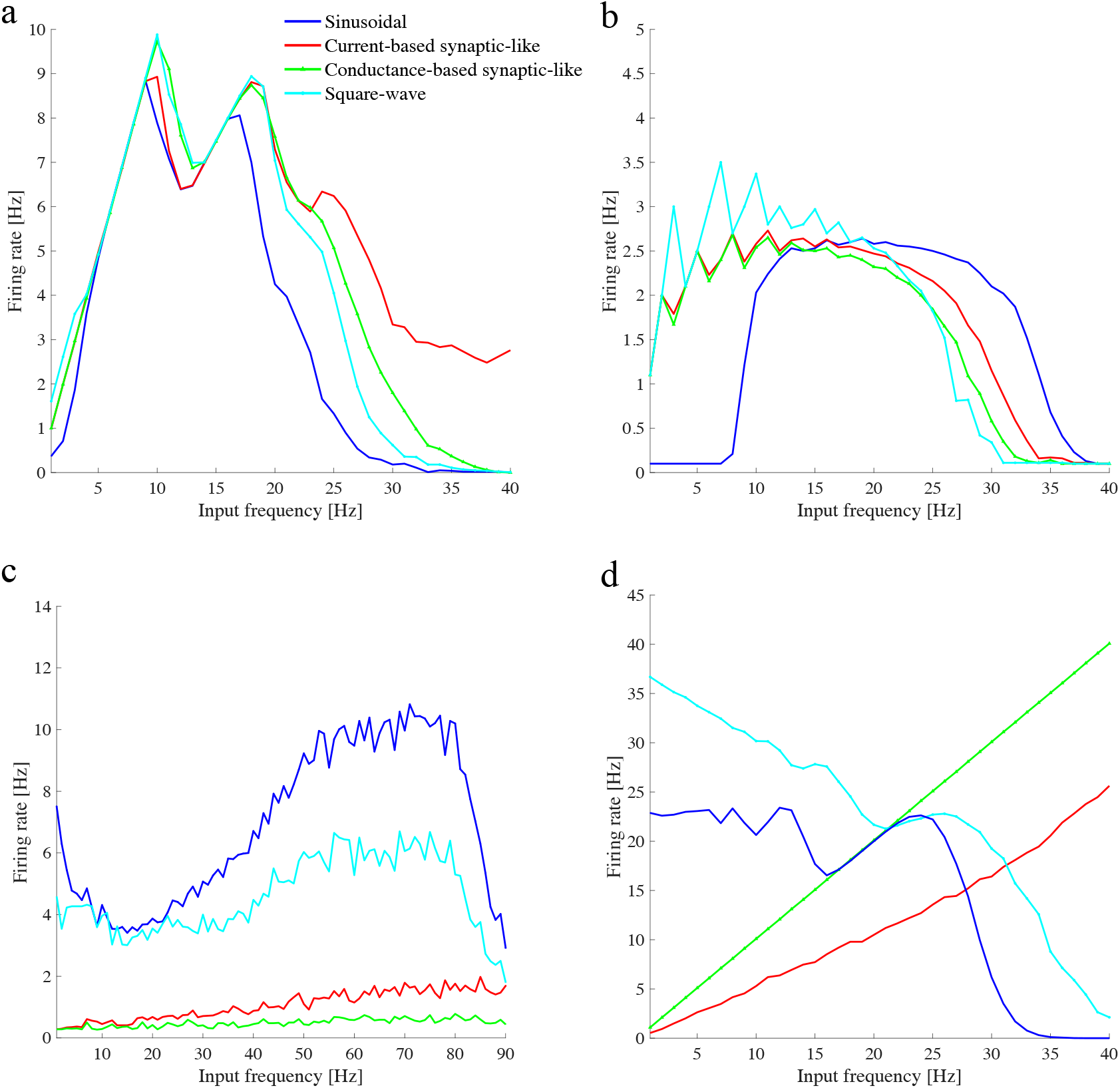
Comparisons between the firing rate responses to different frequency-dependent inputs for the *I*_Nap_ + *I*_h_-model. The curves represent the firing rate of the neuron for each input frequency and input type (see legend). (a) F-cell (same model as in Fig. S23 and Fig. S1). (b) N-cell (same model as in Fig. S2 and Fig. S4). (c) F-cell with resonance at higher frequencies (same model as in Fig. S20). (d) LIF model (1D subthreshold linear dynamics, same parameters as in Fig. 9). Parameter values for are: (a) *v*_*p*,1/2_ = −38 mV, *v_p,slp_* = 6.5 mV, *v*_*r*,1/2_ = −79.2 mV, *v_r,slp_* = 9.78 mV, *C* =1 *μ*F/cm^2^, *E*_L_ = −65 mV, *E*_Na_ = 55 mV, *E*_h_ = −20 mV, *g*_L_ = 0.5 mS/cm^2^, *g_p_* = 0.5 mS/cm^2^, *g*_h_ = 1.5 mS/cm^2^, *I*_app_ = −2.5 *μ*A/cm^2^, *τ_r_* = 80 ms, *v_th_* = −45 mV, *V_rst_* = −55 mV, *r_rst_* = 0, *E*_ex_ = −51 mV, and *τ_Dec_* = 5 ms. *A*_in_ = 0.2 for sinusoidal and square-wave inputs and *A*_in_ = 0.3 for synaptic-like inputs. (b) Same as in (a) except for *g_p_* = 0.1 mS/cm^2^, *v_th_* = −57.63 mV, *V_rst_* = −57.74 mV, *E*_ex_ = −56.65 mV. *A*_in_ = 0.03 for sinusoidal and square-wave inputs and *A*_in_ = 0.06 for synaptic-like inputs. (c) Same as in (a) except for *τ_r_* =4 ms, *v_th_* = −53 mV, and *V*_rst_ = −55 mV. (d) *C* = 1, *g*_L_ = 0.25, *g*_1_ = 0, *τ*_1_ = 100 ms, *v_th_* = 3.5, and *w_r_* = *v_r_* = 0. All simulations in this figure were run for 10 s over 10 trials where noise was added by 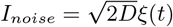.

**Figure S23:**
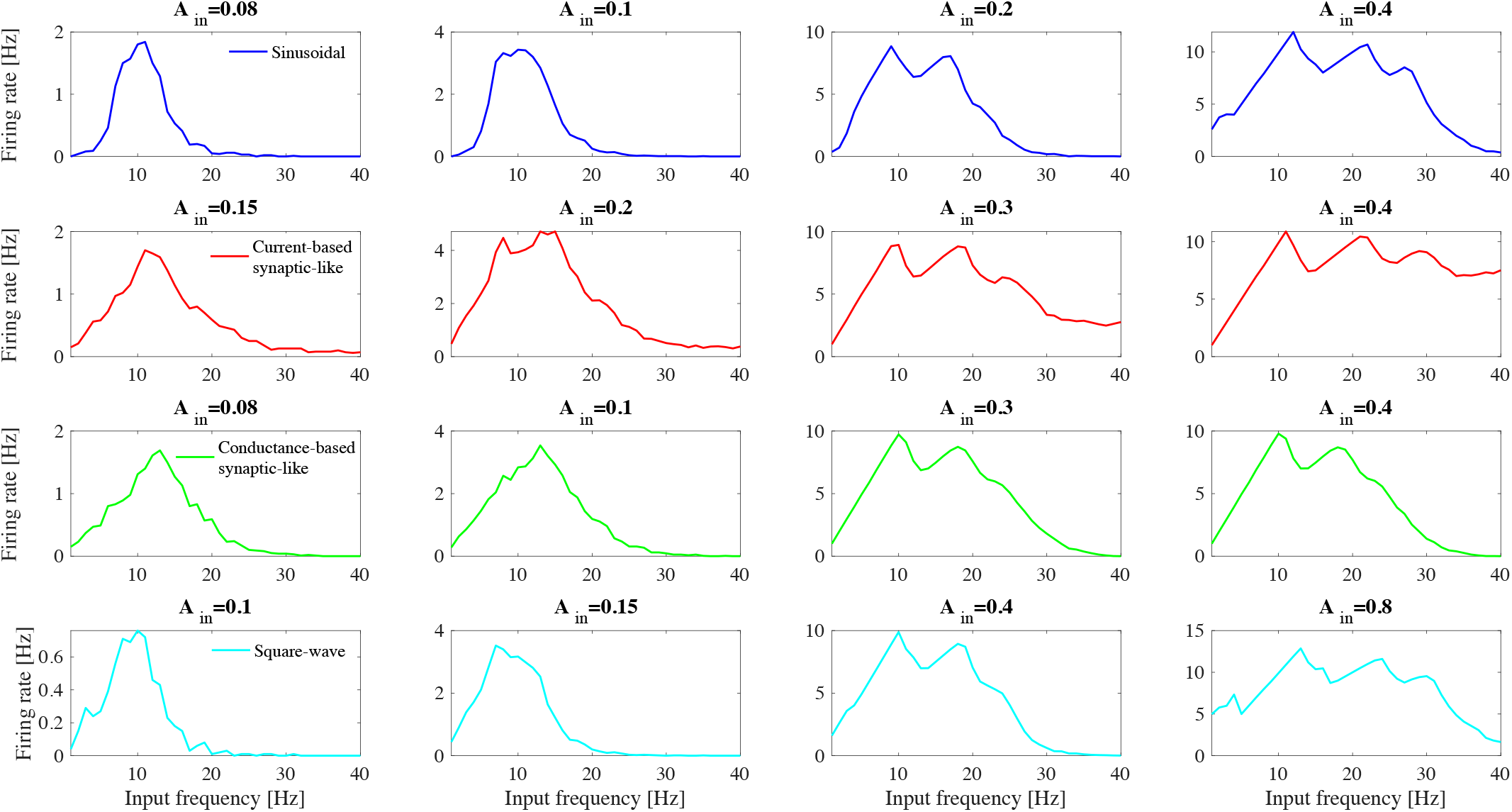
Comparisons between the firing rate responses to different frequency-dependent inputs for the *I*_Nap_ + *I*_h_-model as input amplitude is increased (F-cell). The curves represent the firing rate of the neuron for each input frequency and input type (see legend). Same parameters as in Fig. S22(a) and Fig. S1 (example with subthreshold oscillations). Parameter values are: *v*_*p*,1/2_ = −38 mV, *v_p,slp_* = 6.5 mV, *v*_*r*,1/2_ = −79.2 mV, *v_r,slp_* = 9.78 mV, *C* = 1 *μ*F/cm^2^, *E*_L_ = −65 mV, *E*_Na_ = 55 mV, *E*_h_ = −20 mV, *g*_L_ = 0.5 mS/cm^2^, *g_p_* = 0.5 mS/cm^2^, *g*_h_ = 1.5 mS/cm^2^, *I*_app_ = −2.5 *μ*A/cm^2^, *τ_r_* = 80 ms, *v_th_* = −45 mV, *V_rst_* = −55 mV, *r_rst_* = 0, *E*_ex_ = −51 mV, and *τ_Dec_* = 5 ms. Values of *A_in_* atop each panel. All simulations in this figure were run for 10 s over 10 trials where noise was added by 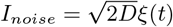.

**Figure S24:**
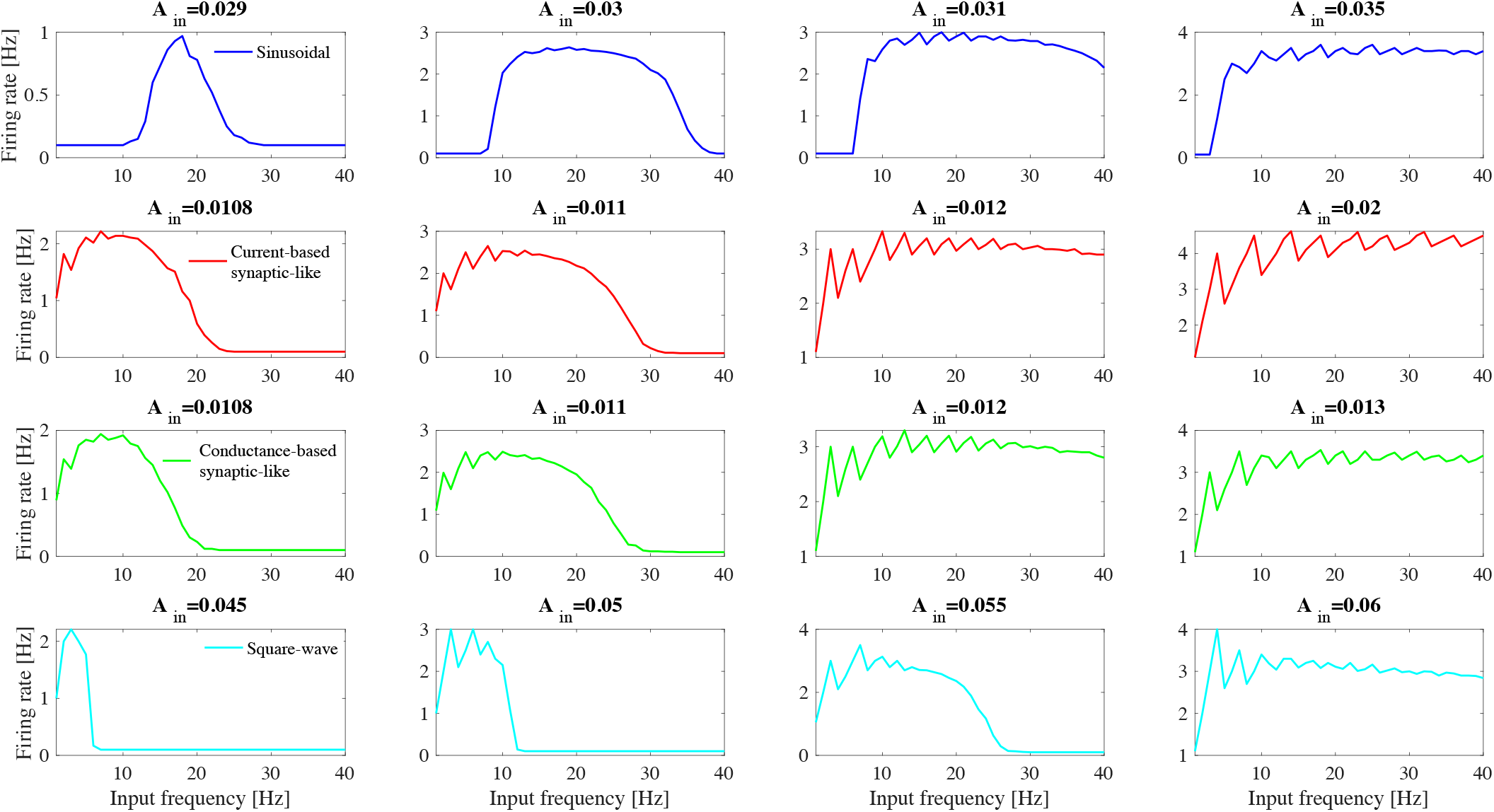
Comparisons between the firing rate responses to different frequency-dependent inputs for the *I*_Nap_ + *I*_h_-model as input amplitude is increased (N-cell). The curves represent the firing rate of the neuron for each input frequency and input type (see legend). Same parameters as in Fig. S2 and Fig. S22(b). Parameters are the same as in Fig. S22(a) except for *g_p_* = 0.1 mS/cm^2^, *v_th_* = −57.63 mV, *V_rst_* = −57.74 mV, and *E*_ex_ = −56.65 mV. Values of *A*_in_ atop each panel. All simulations in this figure were run for 10 s over 10 trials where noise was added by 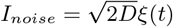.

**Figure S25:**
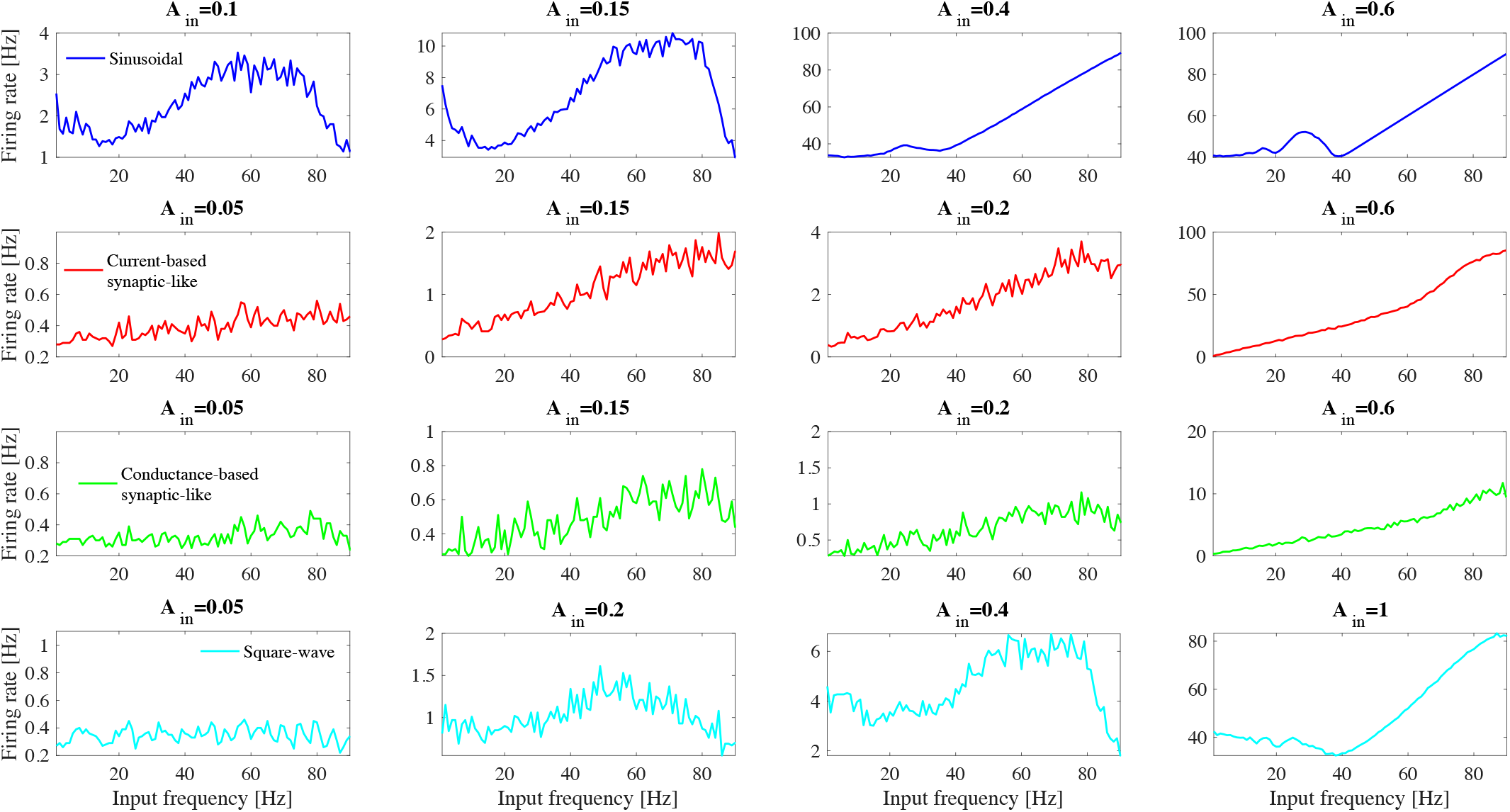
Comparisons between the firing rate responses to different frequency-dependent inputs for the *I*_Nap_ + *I*_h_-model as input amplitude is increased (F-cell, resonance at higher frequencies). The curves represent the firing rate of the neuron for each input frequency and input type (see legend). Same parameters as in Fig. S22(c). Parameters are the same as in Fig. S22(a) except for *τ_r_* =4 ms, *v_th_* = −53 mV, *V_rst_* = −55 mV, and *D* = 0.2. Values of *A*_in_ atop each panel. All simulations in this figure were run for 10 s over 10 trials where noise was added by 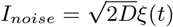.

**Figure S26:**
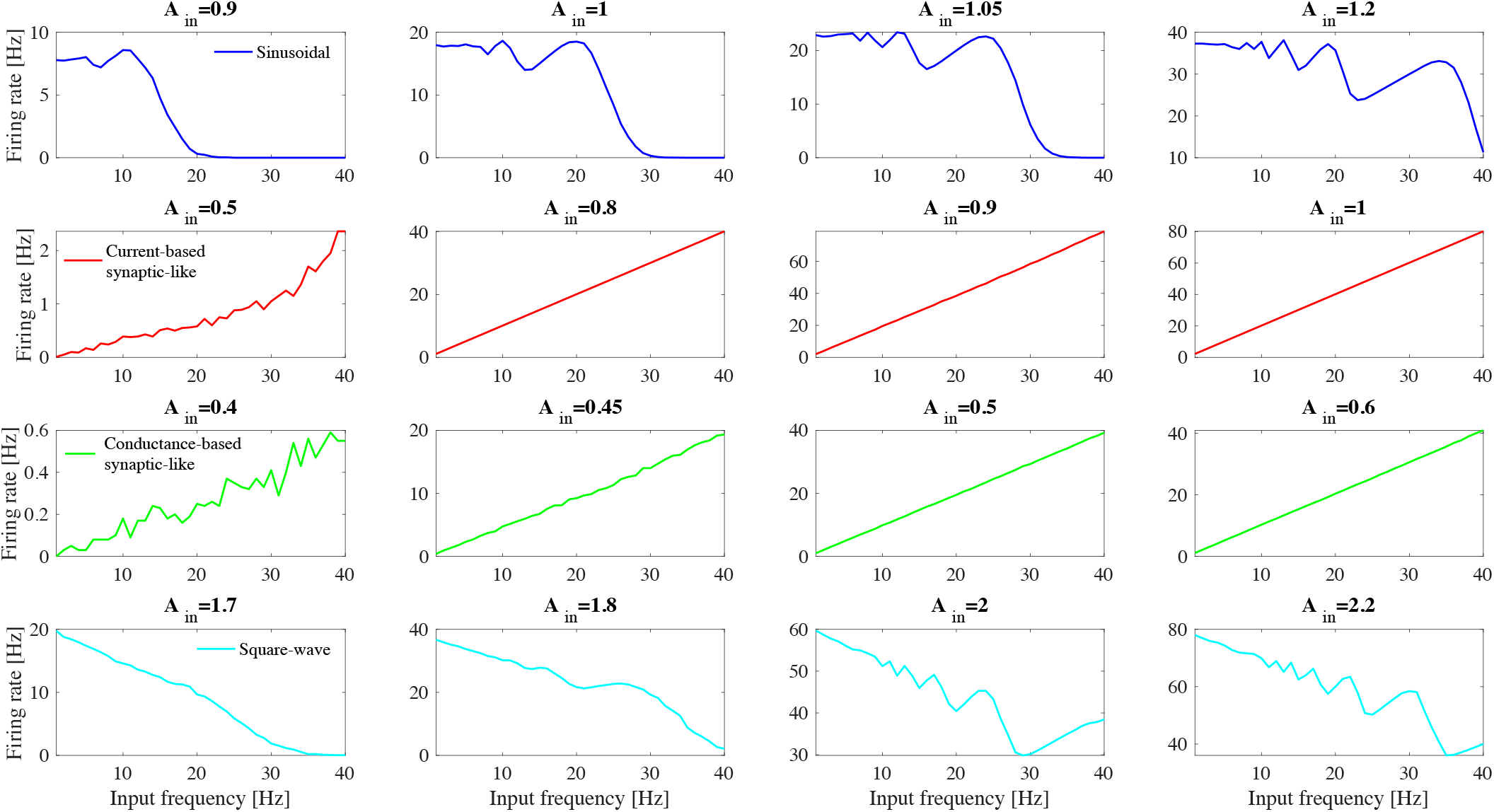
Comparisons between the firing rate responses to different frequency-dependent inputs for the LIF model (1D subthreshold linear dynamics). The curves represent the firing rate of the neuron for each input frequency and input type (see legend). Same parameters as in Fig. 9 and Fig. S22(d). Parameters are *C* =1, *g*_L_ = 0.25, *g*_1_ = 0, *τ*_1_ = 100 ms, *v_th_* = 3.5, and *w_r_ = *v_r_** = 0. Values of *A*_in_ atop each panel. All simulations in this figure were run for 10 s over 10 trials where noise was added by 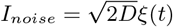.

